# A wheat tandem kinase sensor activates an NLR helper to trigger immunity

**DOI:** 10.1101/2024.08.30.610287

**Authors:** Renjie Chen, Jian Chen, Megan A. Outram, Oliver R. Powell, Taj Arndell, Karthick Gajendiran, Yan L. Wang, Michael A. Ayliffe, Cheryl Blundell, Melania Figueroa, Jana Sperschneider, Thomas Vanhercke, Dingzhong Tang, Yang Xu, Guitao Zhong, Catherine Gardener, Guotai Yu, Spyridon Gourdoupis, Łukasz Jaremko, Oadi Matny, Brian J. Steffenson, Willem H. P. Boshoff, Wilku B. Meyer, Stefan T. Arold, Peter N. Dodds, Brande B. H. Wulff

## Abstract

Most plant resistance genes encode membrane-anchored receptor-like proteins or intracellular nucleotide-binding and leucine-rich repeat (NLR) receptors. In wheat and barley, tandem kinases (TKs) have emerged as a new class of resistance determinants. To understand the modus operandi of the wheat stem rust resistance protein Sr62^TK^, we identified two genetic interactors— a host gene required for Sr62^TK^ function and the corresponding fungal AvrSr62 effector. We discovered that the *SR62* locus consists of a digenic module encoding Sr62^TK^ and an NLR (Sr62^NLR^). AvrSr62 binds to the N-terminal kinase of Sr62^TK^. This triggers displacement of the C-terminal kinase allowing it to recruit Sr62^NLR^ for activation of immune responses. Understanding the mechanism of this two-component resistance complex will help engineering and breeding for durable resistance.

## Main Text

Plant diseases cause about 20-30% losses in crop production annually (*1*). Limiting such losses without recourse to chemicals relies heavily on breeding crops for disease resistance traits. Most known plant resistance (*R*) genes encode nucleotide-binding leucine-rich repeat (NLR) immune receptor proteins, which recognize pathogen effector proteins that are delivered into host cells during infection (*2, 3*). In this context, immune-recognized effectors are known as Avirulence (Avr) proteins, since their recognition triggers immune responses that prevent disease. Many NLRs containing an N-terminal coiled coil (CC) domain function as autonomous units that both detect Avr proteins and then trigger immune responses by forming an oligomeric resistosome that can act as a Ca^2+^ channel (*4–6*). Other NLRs function in pairs, usually encoded by adjacent genes at a digenic resistance locus, wherein one NLR acts as a ‘sensor’ to detect an Avr protein, while the second acts as a ‘helper’ or ‘executor’ NLR to trigger downstream responses (*7, 8*). In some species, complex networks of sensor and helper NLRs have evolved to confer resistance to a variety of pathogens (*9*). In dicotyledonous plants, another class of NLRs contains an N-terminal TIR domain that upon activation catalyzes the production of small signaling molecules that ultimately lead to activation of downstream helper CC-NLRs via the EDS1 family signaling pathway (*2, 10*).

Recently a new class of immune receptor has emerged in *Triticeae* species, with 18 of the 85 (20%) cloned wheat and barley *R* genes found to encode kinase fusion proteins (*11, 12*), including *Sr43* (*13*), *Sr60* (*14*), *Sr62* (*15*), *Yr15* (*16*), *Lr9* (*17*), *Pm24* (*18*), and *Rwt4* (*19*) in wheat, and *Rpg1* (*20*) in barley. Most of these proteins consist of two kinase domains and are known as tandem kinase (TK) proteins, although a few, like Sr43, contain a single kinase domain fused to other domains. It is not known how TK proteins confer pathogen recognition or induce immune responses. Corresponding Avr proteins are unknown, except for RWT4, which recognizes PWT4 from *Pyricularia oryzae* to confer wheat blast resistance (*19, 21*). *Sr62* is derived from the wheat relative *Aegilops sharonensis* (*15*) and confers resistance to stem rust disease caused by the fungus *Puccinia graminis* f. sp. *tritici* (*Pgt*), including against highly virulent *Pgt* strains, such as Ug99, that have caused significant losses worldwide (*22, 23*). Mutational analysis showed that a TK gene at the *Sr62* locus (*Sr62^TK^*) was required for resistance in the Zahir-1644 introgression line, and transgenic expression of *Sr62^TK^*was sufficient to confer resistance in the susceptible wheat cultivar Fielder (*15*). In this study, we investigated how the Sr62^TK^ protein functions to confer resistance in wheat, uncovering a sensor-helper relationship in which Sr62^TK^ detects a corresponding AvrSr62 protein from *Pgt* and activates a helper CC-NLR, also encoded at the *SR62* locus.

### Sr62^TK^ recognizes AvrSr62 protein variants encoded at a complex locus in *Pgt*

To identify corresponding *Avr* genes, we used *Sr62^TK^* to screen a *Pgt* effector library (*24*) by co-expression in wheat protoplasts. Four effector constructs (clone #s 0469, 0472, 0483, 0490) showed significantly reduced expression in the presence of *Sr62^TK^* relative to an empty vector control (fig. S1), suggesting immune recognition-induced protoplast cell death. These four *Avr* gene candidates are members of a larger gene family located at a complex locus on chromosome 5 of the *Pgt* strain Pgt21-0 (Fig. 1A; fig. S2). The chromosome 5A haplotype encodes four gene family members in a ∼70 kbp region, while three genes are present on the chromosome 5B haplotype spanning ∼35 kbp. The three additional genes at the locus were present in the effector library (clone #s 482, 486 and 1059) but were not detected as recognized by *Sr62^TK^* in the screen. All seven *AvrSr62* gene family members were screened individually for recognition and cell death activation by *Sr62^TK^* in wheat protoplasts. Expression of *Sr62^TK^* alone resulted in reduced accumulation of a YFP reporter, indicating some autoactive cell-death induction by this protein (fig. S3). Nevertheless, co-expression of the four positively identified *AvrSr62* variants resulted in enhanced *Sr62^TK^*-dependent cell death, while two other variants (# 0486 and 1059) did not induce *Sr62^TK^*-dependent cell death and the third (# 0482) produced an intermediate response (Fig. 1B). These data confirmed specific recognition of members of this gene family by *Sr62^TK^*and we designated the genes as *AvrSr62-1* to *-7*.

**Fig. 1.**
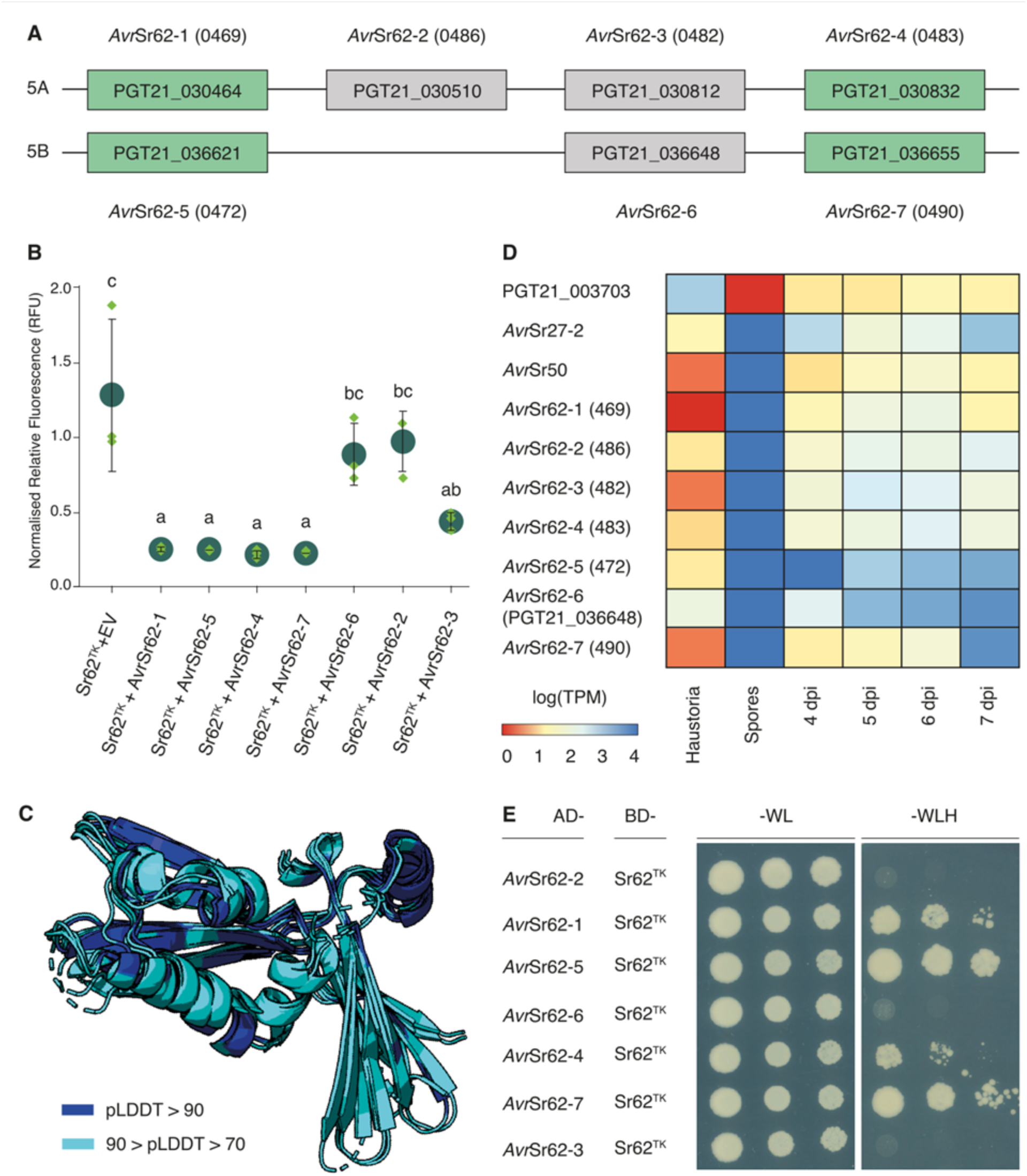
Identification of candidate avirulence effectors for Sr62^TK^ using a high-throughput effector library screening platform in wheat Fielder protoplasts. (**A**) Genomic arrangement of *AvrSr62* genes in the Pgt21-0 genome assembly (*23*). Genes shown in green are recognised by *Sr62^TK^*, while those shown in grey are not recognised. (**B**) Protoplasts from wheat cv. Fielder were co-transformed with YFP and combinations of individual *AvrSr62* genes, *Sr62^TK^* and an empty vector (EV = pTA22) as indicated. YFP fluorescence (y-axis) was measured after 24 hours. Results represent the means (dark green circles) of three biological replicates (green diamonds) with error bars indicating the standard error. Samples marked by identical letters in the plots do not differ significantly (P < 0.05; ANOVA, posthoc Tukey test). (**C**) Superimposed AlphaFold2 structure predictions of the seven AvrSr62 proteins shown in cartoon representation and coloured according to pLDDT (predicted local distance difference test) confidence score as indicated. Only regions with pLDDT >70 are shown. For full models see Figure S5. (**D**) Heatmap of transcript expression levels determined from RNA-Seq data (*27*) for *AvrSr62* genes in comparison to *AvrSr50* and *AvrSr27-2* and a spore-expressed gene (PGT21-003703) in isolated haustoria, germinated spores and infected plants from 2 to 7 days post infection (dpi). Colours correspond to log(Transcript Per Million) as indicated. (**E**) Growth of yeast strains co-transformed with AvrSr62 variants fused to the GAL4 activation domain (AD) and Sr62^TK^ fused to GAL4 binding domain (BD). Yeast suspensions at an OD_600_ = 1.0 and two serial dilutions of 1/10 and 1/100 were spotted on synthetic medium lacking tryptophan and leucine (-WL, growth control) and on synthetic medium lacking tryptophan, leucine and histidine (-WLH, interaction selection). Pictures were taken after 5 days of growth.

The AvrSr62-1 to -7 protein sequences are related (fig. S4) and their high-confidence (average pLDDT > 74.5) AlphaFold predicted structures are highly similar (Fig. 1C, fig. S5). The predicted structures consist of two anti-parallel β-sheets and four α-helices with no predicted disulfide bonds or metal binding pockets as found in some other *Pgt* Avr proteins (*25*). No structural similarities to other proteins were detected in the AlphaFold (Foldseek server) or PDB (Dali server) databases. EffectorP3.0 (*26*) predicts the AvrSr62 proteins as cytoplasmic effectors and RNA-Seq data (*23*) indicates that the *AvrSr62* genes are preferentially expressed in haustoria and during *in planta* infection, but not in germinated spores, similar to other known *Pgt* Avr genes (*27*) (Fig. 1D). However, *AvrSr62-6* shows low expression in all infection stages.

Yeast and *in planta* two-hybrid assays (*28*) showed interaction between Sr62^TK^ and the four recognized AvrSr62 proteins, but not the non-recognized variants, while co-immunoprecipitation assays detected an interaction between Sr62^TK^ and AvrSr62-5 only (Fig. 1E; fig. S6). These data suggest that Sr62^TK^ is the primary receptor for AvrSr62 proteins. However, co-expression of *Sr62^TK^* and *AvrSr62* variants in *Nicotiana benthamiana* leaves or oat (*Avena sativa*) protoplasts did not cause cell death (fig. S7). In contrast, wheat NLR-type *Sr* genes can induce cell death in *N. benthamiana* and oat when co-expressed with corresponding Avr proteins (fig. S7) (*24, 29, 30*). This suggests that the Sr62^TK^-induced immune response in wheat may require other proteins that are not conserved in these heterologous plants.

### The *SR62* locus comprises a digenic module encoding a tandem kinase and an NLR

To identify other wheat genes involved in *Sr62* resistance, we re-examined seven putative EMS-derived susceptible mutants, previously identified amongst a mutagenic population of the wheat-*Aegilops sharonensis* introgression line Zahir-1644, that appeared to maintain wildtype *Sr62^TK^* sequences based on RNA-Seq analysis (*15*). We re-evaluated the phenotypes and sequences of these mutants aided by additional whole genome shotgun-sequences and a high-quality Zahir-1644 genome assembly (Contig N50, 21 Mb) (table S1 and table S2). Four partially or completely susceptible mutants were confirmed to encode intact *Sr62^TK^* open reading frames (mutants 119d, 267d, 200h and 1298e; Fig. 2A, tables S3 and S4, and dataset S1). Previously, we observed an NLR-encoding gene 20.4 kbp distal to *Sr62^TK^* (*15*). The gene encodes a predicted 1040-amino acid protein with domains resembling a coiled-coil, a nucleotide-binding site with two NB-ARC domains, and 12 leucine-rich repeats (Fig. 2B-D and fig. S8 and table S5). We identified missense mutations (119d and 1298e) or nonsense mutations (267d and 200h) in this NLR gene in each of the four mutants (Fig. 2C, table S4, and dataset S1). We backcrossed the four mutant lines with Zahir-1644 and identified BC_1_F_2_ susceptible and resistant plants (table S6). These were genotyped by bulk exome-capture sequencing and/or PCR and Sanger sequencing of individual plants. In all cases, we observed complete co-segregation between susceptibility and the NLR mutations (table S7 and dataset S1). No other instances of a gene containing mutations in three or four mutants were identified (table S8).

**Fig. 2.**
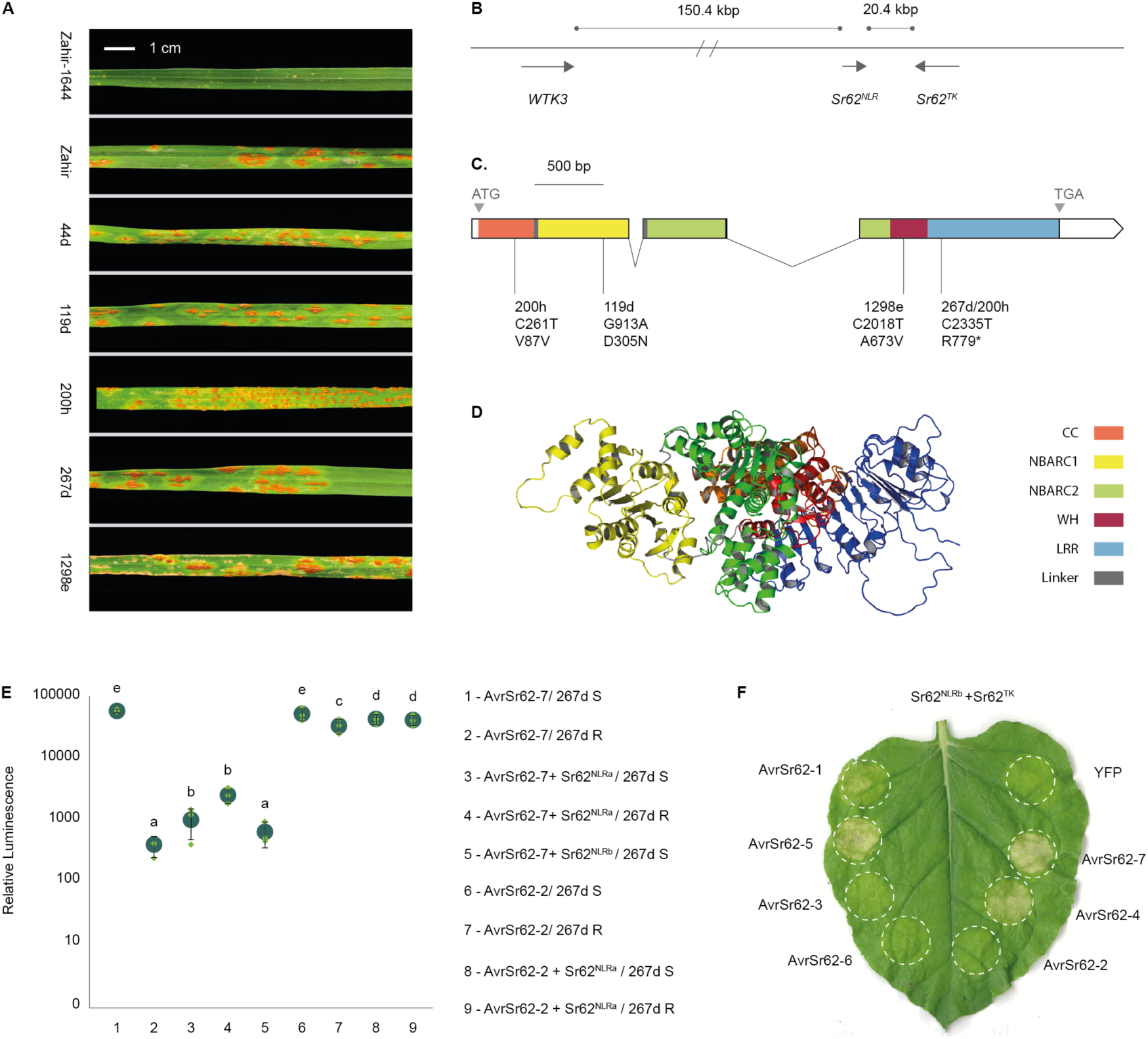
Discovery of an NLR essential for the function of the *Sr62* tandem kinase. (**A**) Infection types of the wheat-*Ae. sharonensis* introgresion line Zahir-1644, the recurrent parent cultivar Zahir, and EMS-derived mutants inoculated with *Puccinia graminis* f. sp. *tritici* (*Pgt*) race TTKSK. Scale bar, 1 cm. (**B**) The *Sr62^NLR^*is flanked by the paralogous *Sr62^TK^* and *WTK3* (*Pm24*/*Rwt4* orthologues). (**C**) *Sr62^NLR^* gene structure with predicted nucleotide changes caused by EMS-derived loss-of-function mutations (top two lines) and corresponding amino acid changes (bottom line). Colored and white boxes represent predicted translated and untranslated exons, respectively, while black connecting lines represent introns. The predicted structural domains encoded by the exons are colored and labelled: coiled coil (CC, orange), Nucleotide-Binding Adaptor Shared by APAF-1, R proteins, and CED-4 (NBARC)-1 (yellow), NBARC2 (green), winged helix (WH, red) and leucine-rich repeat (LRR, blue). (**D**) AlphaFold-predicted structure of Sr62^NLR^ coloured by domains as in (C). (**E**) Protoplasts of BC1F2 lines derived from the Zahir-1644 *Sr62^NLR^* 267d mutant line and either homozygous susceptible (267d S) or resistant (267d R) were co-transformed with a *Luciferase* construct and either *AvrSr62-2* or *-7*, with or without *Sr62^NLR^*. Graph shows the mean (dark green circle) luminescence of three replicates (green diamonds) with the standard error indicated (black line) on a log scale. Samples marked by identical letters in the plots do not differ significantly (P < 0.05; ANOVA, posthoc Tukey test). *Sr62^NLRa^* and *Sr62^NLRb^* denote *Ae. sharonensis* and *T. aestivum* cv. Fielder chromosome 1D alleles, respectively. (**F**) Co-expression of *Sr62^TK^*, *Sr62^NLRb^* and *AvrSr62* induces cell death in *N. benthamiana*. Sr62^TK^ and Sr62^NLR^ are fused to 3xHA tag at the C-terminus, AvrSr62 effectors are fused to a YFP tag at the N-terminus.

Transient expression of *AvrSr62-7* induced cell death in wildtype Zahir-1644 protoplasts, but not in protoplasts of NLR mutant Zahir-1644 lines (Fig. 2E; fig. S9). However, co-expression of the wild type NLR gene with *AvrSr62-7* did induce cell death in protoplasts from 267d, 1298e and 200h mutant lines (Fig. 2E; fig. S9). These experiments confirm the requirement of this NLR for *Sr62* function. Previously, we transformed *Sr62^TK^* into wheat cv. Fielder and recovered disease resistant transgenics (*15*), suggesting that the orthologue of this NLR gene on chromosome 1D of Fielder (94.6% amino acid identity to its *Ae. sharonensis* counterpart) supports *Sr62^TK^* resistance function. We confirmed this by transient complementation with the Fielder NLR in protoplasts from the 267d mutant line (Fig. 2E). Finally, transient co-expression of *Sr62^TK^*, the Fielder *NLR* and recognized *AvrSr62* variants resulted in cell death in *N. benthamiana* and *N. tabacum* (Fig. 2E; fig. S10), demonstrating heterologous recapitulation of the interaction specificity observed in wheat. In conclusion, the *SR62* locus comprises a digenic module encoding a tandem kinase and a linked NLR, which we designated *Sr62^TK^* and *Sr62^NLR^*, respectively.

### Sr62^TK^ and Sr62^NLR^ work as a sensor-helper complex in AvrSr62 detection

The interaction of Sr62^TK^ with AvrSr62 suggests that Sr62^TK^ acts as a sensor while Sr62^NLR^ may function as a helper NLR. Split luciferase complementation assays (*31*) showed that co-expression of AvrSr62-7 with Sr62^TK^ and Sr62^NLR^ fused to the N- and C-terminal domains of luciferase, respectively, induced an interaction resulting in strong luminescence (Fig. 3A). Co-expression experiments using recognized AvrSr62 proteins also led to enhanced co-immunoprecipitation of Sr62^NLR^ by Sr62^TK^ (fig. S11). These results suggest that recognized AvrSr62 proteins promote an interaction between the TK and the NLR that activates immunity.

**Fig. 3.**
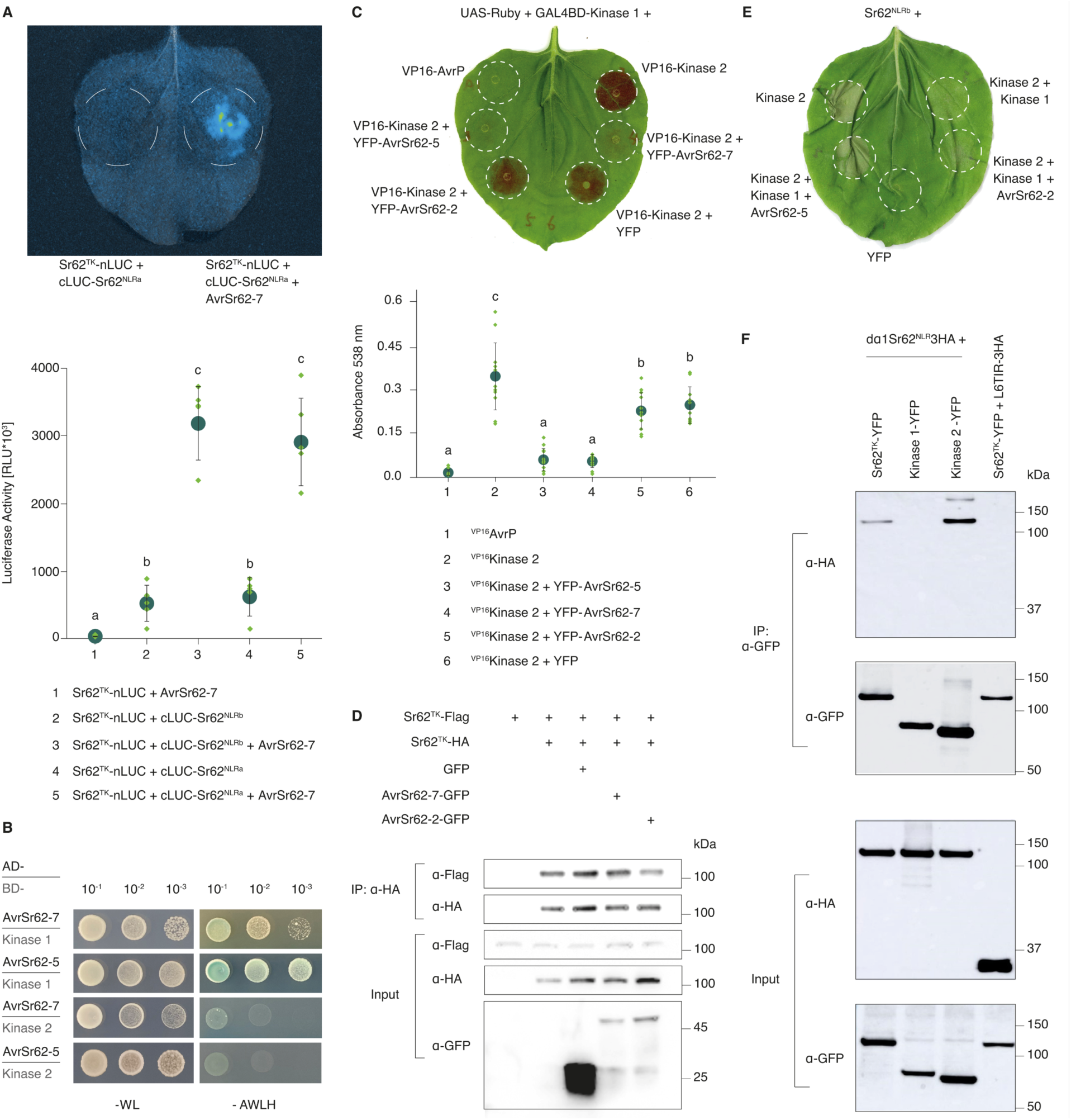
Sr62^TK^ Kinase 1 and 2 regulate AvrSr62-induced recruitment of Sr62^NLR^ to induce cell death. (**A**) Split-luciferase complementation assay. *N. benthamiana* leaves were co-infiltrated with Agrobacterial strains containing different pairs of constructs. Luciferase images were captured at 48 hpi (example, top) and the luminescence quantified (bottom). Graph shows mean (dark green circle) luminescence of three replicates (green diamonds) with the standard error indicated. Samples marked by identical letters in the plots do not differ significantly (P < 0.05; ANOVA, posthoc Tukey test). (**B**) Yeast two-hybrid assay. The AvrSr62-5 and AvrSr62-7 coding sequences were fused to the Gal4 transactivation domain (AD) whereas the *Sr62^TK^* Kinase 1 and Kinase 2 coding sequences were fused to the Gal4 DNA binding domain (BD). Pairs of constructs were co-transformed into AH109, plated as a serial dilution on selective media lacking tryptophan and Leucine (-WL, left) and media lacking tryptophan, leucine, histidine and adenine and with the chromogenic indicator X-Gal (-AWLH, right). Photographed 4 dpi. (**C**) Plant two-hybrid assay. Betalain accumulation was observed in leaves expressing UAS-Ruby with Kinase 1 fused to BD and Kinase 2 fused to VP16 and co-expressed with AvrSr62 variants. VP16-AvrP was used as negative control. Leaves were photographed 3 days after infiltration (top). Extracted betalain was quantified by its absorbance at 538 nm (Right). Graph (bottom) presents results of at least six biological replicates (green diamonds) with the average (green circle) and standard error indicated. Samples marked by identical letters in the plots do not differ significantly (P < 0.05; ANOVA, posthoc Tukey test). (**D**) Co-immunoprecipitation (Co-IP) assay. Flag-tagged *Sr62^TK^* was transiently co-expressed in *N. benthamiana* with HA-tagged *Sr62^TK^*, GFP, and GFP-tagged AvrSr62-7 and AvrSr62-2. Total protein extracts immunoprecipitated with an anti-HA antibody were immunoblotted with anti-GFP, anti-Flag, or anti-HA antibodies. (**E**) Sr62^NLR^ was transiently expressed in *N. benthamiana* along with Kinase 1, Kinase 2, AvrSr62-2, AvrSr62-5 or YFP in the indicated combinations. Sr62^NLR^ and Kinase 1, Kinase 2 are fused to 3xHA tag at the C-terminus, AvrSr62 effectors are fused to a YFP tag at the N-terminus. (**F**) Co-IP assay. YFP-tagged Sr62^TK^, Kinase 1 and Kinase 2 were transiently co-expressed in *N. benthamiana* with HA-tagged Sr62^NLRb^ lacking the first alpha helix (dα1Sr62^NLR^). Total protein extracts were immunoprecipitated with anti-GFP beads (IP), followed by immunoblotting with anti-GFP, and anti-HA antibodies. Sr62^TK^-YFP co-expression with L6TIR E135A (*38*) (no cell death) with HA tag was used as negative control.

Yeast two-hybrid experiments showed that recognized AvrSr62 variants interact with the N-terminal Kinase 1 domain of Sr62^TK^, but not with the C-terminal Kinase 2 domain (Fig. 3B), which was further confirmed by *in planta* two-hybrid and Co-IP assays (fig. S12 and 13). No interaction was detected between any AvrSr62 proteins and the NLR protein in yeast or *in planta* two-hybrid experiments (fig. S14). We also observed interaction between the separated Kinase 1 and Kinase 2 domains in Co-IP and *in planta* two-hybrid assays (Fig. 3C and fig. S15), as well as self-association of the full-length Sr62^TK^ and of the Kinase 1 domain (Fig. 3D and fig. S16). Co-expression of recognized AvrSr62-5 or -7 proteins suppressed the association between Kinase 1 and Kinase 2 fragments, while the interaction was maintained in the presence of the non-recognized AvrSr62-2 protein (Fig. 3C and fig. S15 and fig. S17). In contrast, TK self-association was not affected by the AvrSr62 proteins, while Kinase 1 self-association was enhanced in the presence of AvrSr62-5, but not AvrSr62-2 (fig. S16).

We next investigated molecular interactions between the TK and the NLR. Co-expression of Kinase 2, but not Kinase 1, with Sr62^NLR^ was sufficient to induce a strong cell death response in *N. benthamiana* and *N. tabacum* (Fig. 3E; fig. S18A). This effect was dependent on an intact alpha-1 helix in Sr62^NLR^ (fig. S18B) and was not observed with unrelated NLRs Sr33 and Sr50 (fig. S18C). However, when Kinase 1 and Kinase 2 were both expressed in the presence of the NLR, this cell death was strongly reduced or absent (Fig. 3E and fig. S19), suggesting that Kinase 1 negatively regulates Kinase 2-induced cell death. In the presence of the recognized AvrSr62-5, but not the unrecognized AvrSr62-2, the cell death response was re-established indicating that AvrSr62-5 detection releases Kinase 1-mediated inhibition of Kinase 2 (Fig. 3E and fig. S19). Similar results were observed in wheat protoplast assays (fig. S19) and Co-IP experiments showed interaction between Kinase 2 and Sr62^NLR^ (Fig. 3F). Overall, these data suggest a model in which interaction of a recognized AvrSr62 protein with the Kinase 1 domain disrupts an intramolecular interaction in Sr62^TK^ which then allows Kinase 2 to interact with Sr62^NLR^ to activate immunity.

Subsequently, we explored the 3D protein structures and their interactions *in silico* using AlphaFold (*32*). Based on the predicted scores for the structures and their interactions (fig. S20 and table S9), AlphaFold supported our experimentally-derived model by confidently predicting Kinase 1 homodimerization, Kinase 1 and 2 heterodimerization, and Kinase 2 and Sr62^NLR^ heterodimerization (Fig. 4, fig. S21 A to D, L to N, and table S10). AlphaFold also predicted associations between Kinase 1 and the recognized AvrSr62-1, -4, -5 and -7 effectors, whereas the predictions with the other family members showed weaker scores (fig. S21 E to H vs I to K, and table S10). AlphaFold models indicated that Kinase 1 contains all features needed for catalysis (including p-loop and catalytic residues Lys86 and Glu83), whereas key catalytic features were absent in Kinase 2, suggesting it is a pseudokinase (*13*).

**Fig. 4.**
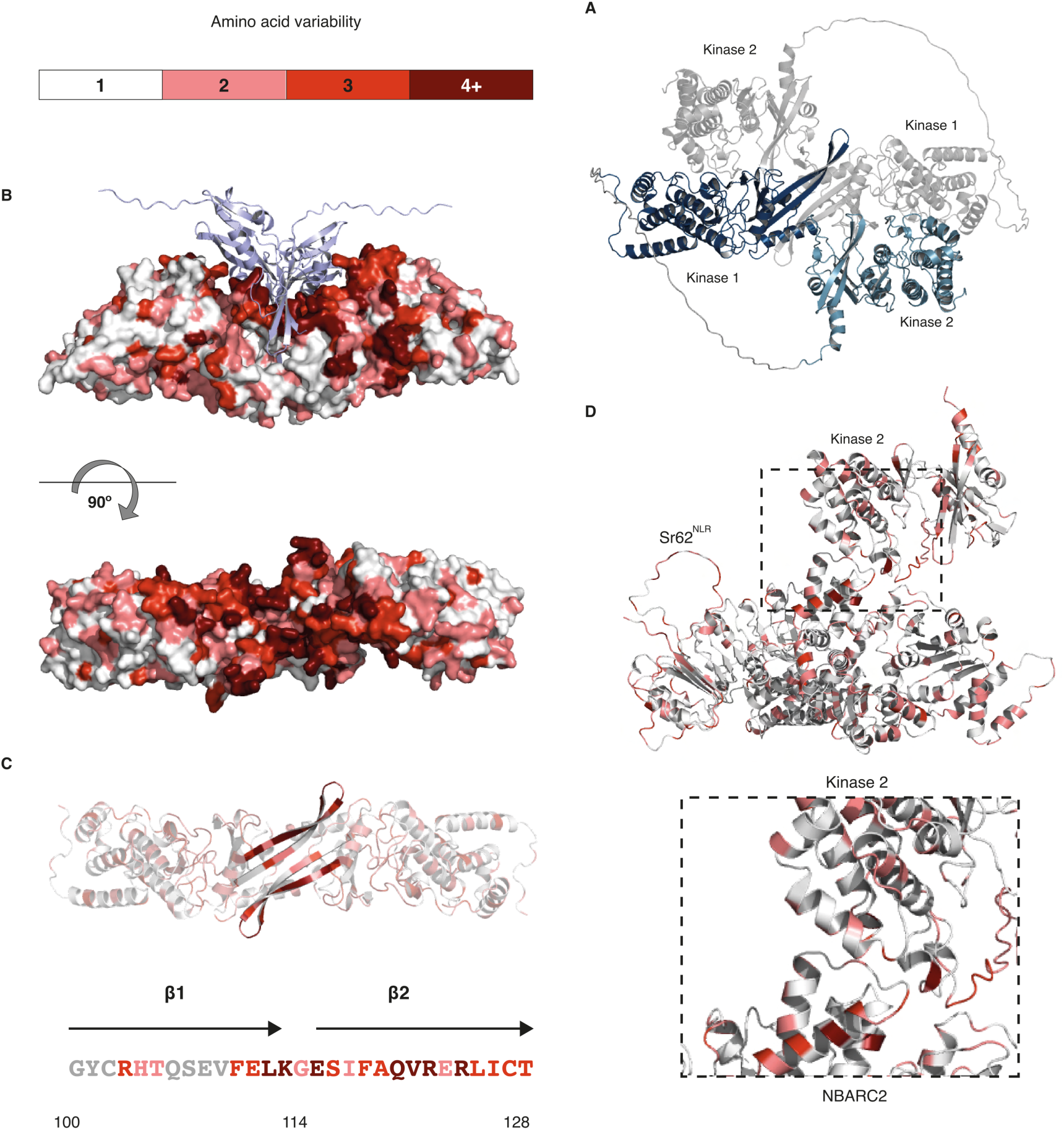
Predicted structures, interactions, and amino acid variability in the ‘Sr62 complex’. (A) AlphaFold-predicted structure of the Sr62^TK^ homodimer, with each monomer subunit represented in grey or blue. Sr62^TK^ dimerization is mediated by Kinase 1/Kinase 2 and Kinase 1/Kinase 1 interactions. **(B)** Interaction of AvrSr62 (light purple) with the Sr62^TK^ Kinase 1 homodimer colored according to amino acid variability as indicated in the scale. (**C)** The extended ß-finger structure mediating Sr62^TK^ homodimerization through the conserved ß1 and effector interaction through the variable ß2. **(D)** Interaction of Sr62^NLR^ with Sr62^TK^ Kinase 2 through the APAF-1 domain of NBARC2 in the Sr62^NLR^ and the phosphotransferase domain of Sr62^TK^. The distribution of amino acid variability (panels B to D) is based on sequence-comparison of *Triticum* and *Aegilops* homologues.

Compared to canonical kinase folds, the last two β-strands of the predicted Kinase 1 N-lobe extend by eight residues each, forming an unusual β-hairpin finger structure (termed β-finger; residues 100–128). (Fig. 4B to C). The first strand of this β-finger contributes to the predicted homodimerization interface of Kinase 1, whereas the second strand plays a key role in the predicted binding of the effector molecules through a mostly hydrophobic interaction (fig. S21L and fig. 4.3). The hydrophobicity of the effector residues at this site correlates with their experimentally observed capability to interact with Sr62^TK^ (fig. S22). The position of Kinase 2 in the highest-scoring AlphaFold Sr62^TK^ homodimer predictions was largely overlapping with the position of the effectors (fig. S21 L and N), supporting our model that recognized AvrSr62 effectors compete with Kinase 2 for the same site on Kinase 1. AlphaFold further predicted that Kinase 2 uses overlapping surfaces to bind to both Kinase 1 and the NB-ARC2 of Sr62^NLR^, consistent with both events being mutually exclusive.

Mapping of the EMS-induced mutations onto the 3D structures (fig. S23) revealed that mutation E10K in the 44d Sr62^TK^ mutant is predicted to disrupt the putative homodimer interaction surface, while T627M in the 905c mutant appears to perturb the Kinase 2 interaction surface that binds to Kinase 1 and Sr62^NLR^, underscoring the importance of these putative interaction sites (fig. S23). We also examined natural variation in Sr62^TK^ and Sr62^NLR^ homologues in 76 genome assemblies across the *Triticum* and *Aegilops* genera, identifying 19 and 25 unique variants, respectively (fig. S24, fig. S25 and table S11). A polymorphic hotspot in Kinase 1 coincides with the putative AvrSr62 interaction surface (Fig. 4B to C), suggesting that effector variation drives selection for variation at this site. In contrast, we found Kinase 2 and the NLR to be largely conserved (Fig. 4D). Collectively, the computational and experimental analyses support a molecular mechanism whereby Sr62^TK^ adopts a dimeric ‘closed’ autoinhibited conformation in the absence of recognized effector molecules. Effector binding to the β-finger of Kinase 1 releases Kinase 2 for interactions with Sr62^NLR^ NB-ARC2 leading to activation of immune signaling (Fig. 5).

**Fig. 5.**
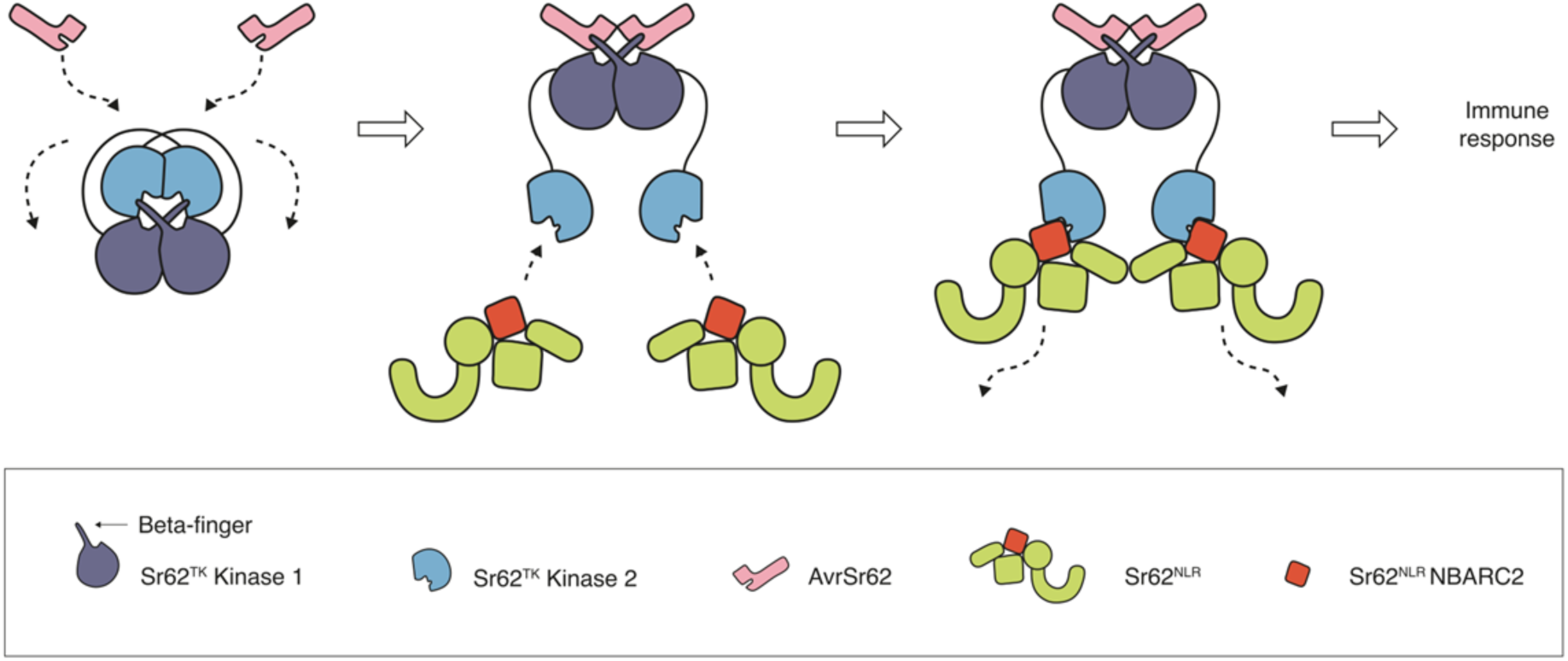
Proposed mechanism of Sr62^NLR^ activation upon detection of AvrSr62 by Sr62^TK^. In the resting state, Sr62^TK^ forms a homodimer in which the C-terminal Kinase 2 overlaps and binds the other molecule’s N-terminal Kinase 1 at the β-finger. Upon AvrSr62 effector binding to the Kinase 1 β-finger the Kinase 1 interaction with Kinase 2 is disrupted. Release of Kinase 2 allows Kinase 2 to recruit Sr62^NLR^ through the NLR NBARC2 domain.

## Discussion

Kinases play important roles in plant signaling, regulating growth, metabolism, and responses to the environment (*33*). In PAMP-triggered immunity, kinase cascades lead to the phosphorylation of transcription factors and defense proteins, triggering defense gene expression and cell wall reinforcement (*34*). This central role of kinases makes them prime targets of pathogen virulence effectors (*35*). In some cases, these effectors can be directly recognized by NLRs or indirectly by kinase decoys guarded by NLRs to induce immunity (*2, 3*). The present study of the wheat stem rust resistance locus *SR62* are consistent with the workings of a novel multi-step immune-switch. In our model, Sr62^TK^ exists as a homodimer in which the N- and C-terminal kinases sit next to each other in the inactive state. Upon AvrSr62 effector binding to the N-terminal kinase β-finger (“sensor”), the N-terminal kinase (“repressor”) releases the C-terminal pseudo-kinase (“activator”). This allows the C-terminal kinase to recruit Sr62^NLR^ (“executor”) to trigger NLR immune signaling. Our model has analogies with the ZAR1/ZED1/PBL2 system in Arabidopsis, although in this system the kinases are encoded in trans, and the PBL2 kinase sensor is recruited to the ZAR1 (NLR)/ZED1 (pseudokinase) complex upon bacterial effector activation (*4*). Given that the kinase catalytic site mutation D177N disrupts Sr62^TK^ function (*15*) (fig. S23), phosphorylation events may also contribute to the activation process.

*Sr62^TK^* is located 20.4 kbp proximal to the *Sr62^NLR^*. The allelic powdery mildew and wheat blast resistance genes *Pm24* and *Rwt4*, respectively, are *Sr62^TK^* paralogues that reside 151.4 kbp distal to the *Sr62^NLR^* (*18, 19*) (Fig. 2B). Strikingly, Lu and colleagues recently found that Pm24 and Rwt4 signal through the same NLR as Sr62^TK^ (this issue). This is akin to the tomato Prf /Pto system, in which the *Prf* NLR-encoding gene is embedded within a complex locus of *Pto* kinase homologues (*36*). It is not known whether other wheat kinase fusion proteins require NLR helpers for their function, or whether such helpers are always encoded by physically linked genes.

We observed extensive haplotype variation at the *SR62* locus in wheat (fig. S24 and fig. S25). Possibly not all the *Sr62^NLR^* helper variants would support *Sr62^TK^* function. In natural populations and breeding programs, digenic modules ensure co-inheritance of functionally co-adapted genes. By extension, when using two-component resistance determinants in multi-resistance gene stacks (*37*), it will be important to ensure inclusion of both components in the stack. We found that Sr62^TK^ recognizes multiple AvrSr62 variants, with two recognized variants encoded in each of the two haplotypes of the Pgt21-0 and Ug99 strains (fig S2). This reduces the risk of mutation to virulence and along with the broad-spectrum resistance conferred by *Sr62* (*15*), highlights its value in resistance breeding. In conclusion, Sr62^TK^ and Pm24/Rwt4 represent a novel mechanism characterized by the interaction of kinase fusion proteins and NLRs.

## Materials and Methods

### Plant material

Wheat germplasm used in this study include wheat cv. Zahir, cv. Fielder, cv. Sonora, wheat introgression line Zahir-1644 and Zahir-1644 mutants (table S3). Wheat plants were grown in a growth room at 22° C under a 16 h light/8 h dark photoperiod.

### Plasmid construction design and molecular cloning

Plasmids used in this study are listed in table S12. Oligonucleotide primers are listed in table S13. The coding sequence (CDS) of the *Sr62^TK^* was synthesised in pDONR207 and cloned into the pTA22-GW vector via Gateway LR reaction (Invitrogen). The CDS of *Sr62-^NLRa^* (*Ae. sharonensis* allele) and *Sr62^NLRb^* (Fielder allele) were synthesized in the TWIST pENTR Kozak vector (Twist Biosciences, San Francisco, USA). Other constructs were generated using Gateway or restriction cloning as detailed in table S12 and described below. The Kinase 1 and Kinase 2 fragments were derived from amino acids 1-389 and 380-740 of Sr62^TK^, respectively. For transient expression and Co-IP assays in *Nicotiana spp.*, *Sr62^TK^*, Kinase 1, Kinase 2 and *Sr62^NLR^*constructs were generated in the pAM-PAT vector, while effector gene expression constructs were in the pBIN19 vector. For in plant two-hybrid assays, GAL4BD and VP16 fusion vectors are as described previously (*28*). All vectors were sequenced to confirm correct insertions.

### Protoplast isolation from wheat and oat

Seeds (∼15) of wheat (*Triticum aestivum*) lines or oat (*Avena sativa*) cultivar ‘Swan’ were planted in 13 cm pots containing Martins Seed Raising and Cutting Mix supplemented with 3 g/L osmocote. Seedlings were grown in a growth cabinet at 24°C on a cycle of 12 hours light (∼100 µmol m^-2^ s^-1^) and 12 h dark, for 7-8 days. The first leaf of wheat or second leaf of oat was selected for protoplast generation. Protoplast isolation and transformation was carried out as described previously (*24, 39*). Released protoplasts were resuspended to a final concentration of either 4.0 × 10^5^ cells/mL using MMG solution (4 mM MES-KOH [pH 5.7], 0.4 M mannitol, 15 mM MgCl_2_) for library screening experiments or 3.0 × 10^5^ cells/mL for individual transformations.

### Stem rust effector library screening in wheat protoplasts

A library of 696 effector candidates from *Pgt* was described previously (*24*). An additional 677 candidates were synthesised and two library pools (part A and part B, containing 718 and 655 clones respectively) were prepared (*40*). Pooled libraries were were propagated in ElectroMAX™ Stbl4™ competent cells and DNA was isolated using the QIAGEN EndoFree Plasmid Giga kit, as described previously (*24*). The reporter (pTA22-*YFP*), empty vector (pTA22), and pTA22-*Sr62^TK^* plasmids were isolated from *E. coli* DH5α using the Macherey Nagel NucleoBond Xtra Maxi Plus EF kit. A total of 250,000 protoplast cells of wheat cultivar Sonora were transformed with with a pooled effector library at a multiplicity of transfection (MOT) of 0.14 million molecules per cell for each construct along with either the empty vector pTA22 or pTA22-*Sr62^TK^* (MOT = 36 million molecules per cell, 45 and 66 μg, respectively) in triplicate as described (*24*). A control protoplast sample (50,000 cells) was transformed with pTA22-*YFP* (10 μg; MOT = 36 million molecules per cell) alone to assess transfection efficiency by flow cytometry.

Messenger (m)RNA was extracted from transformed protoplasts after 24 hours using the NEB Magnetic mRNA Isolation Kit (#S1550S). Library-specific cDNA synthesis and PCR was carried out on 30 ng mRNA using the Invitrogen SuperScript IV One-Step RT-PCR System with ezDNase kit (Cat. # 12595100) with the forward and reverse primers ZmUbi1_5UTR_F3b (5’- GCACACACACACAACCAG-3’) and FS_cDNA_R (5’-TGCTAGATCTCGACAGTACG-3’) as previously described (*24*). Illumina libraries were generated for each cDNA sample (three replicates of each treatment and control) and the pooled effector library before and after propagation using the Illumina DNA Prep Kit (Cat. # 20060059) and IDT for Illumina DNA/RNA UD Indexes Set A (Cat. # 20027213) following the manufacturer’s protocol. Illumina libraries were sequenced using the NextSeq2000 at the ACRF Biomolecular Resource Facility, The John Curtin School of Medical Research, Australian National University using the NextSeq P1 300 cycles kit, with 150 bp paired end reads.

Differential expression analysis was carried out as described (*24*). RNA sequencing reads were cleaned using fastp 0.22.0 (*41*) (--length_required 20) specifying the UTR sequences common to all library transcripts as well as the Illumina DNA Prep adapter sequence as adapters. The clean reads were aligned to the coding sequences of the effector candidates with HISAT2 2.2.1 (*42*) (--very-sensitive; --sp 1,1; --no-spliced-alignment). Mappings where the read pairs map to different transcripts were dismissed. Salmon 1.8.0 (*43*) was used to quantify expression from the HISAT2 alignments. Read counts were imported into DESeq2 (*44*) with tximport (type = “salmon”) for differential expression analysis using default parameters followed by lfcShrink (type=“apeglm”) to compare the *Sr62^TK^* and empty vector treatments. DESeq2 uses the Wald test to compare expression between two samples and reports p-values adjusted for multiple testing using the Benjamini and Hochberg method (BH-adjusted p values). Volcano plots were produced with EnhancedVolcano (https://github.com/kevinblighe/EnhancedVolcano). Syntenic regions in *Pgt* genome assemblies were called with minimap2 (-X -N 50 -p 0.1 -c) (*45*) and plots were produced with gggenomes (https://github.com/thackl/gggenomes.)

### YFP fluorescence-based cell death assays in protoplasts

YFP fluorescence of protoplast suspensions was measured 24 hours after transformation with pTA22-YFP and various combinations of R/Avr gene constructs as described above.

Protoplast suspensions were transferred from 24-well culture plates to 2 mL tubes and allowed to settle via gravity for 30 min before removing 350 μL of solution from each sample. Samples were gently rotated to generate a homogenous protoplast suspension and 130 μL (in duplicate) was transferred to a black Cliniplate 96-well plate (9502767). YFP fluorescence was measured using a FluoStar Omega plate reader (excitation filter = 485-12, emission filter = 520). Gain was set to 60% of the well with the highest fluorescence (pTA22-YFP with pTA22 empty vector) and fluorescence was normalized against either Sr62 or the empty vector sample, as indicated in the figure captions.

### Luciferase-based cell death assays in protoplasts

Wheat protoplasts were isolated from 10-day-old Zahir-1644 and mutant plants and transformed (15 μg plasmid per 50,000 protoplasts) as described (*46*) with constructs pICH47811 (Addgene plasmid # 48008) with ubiquitin:*AvrSr62-7* C-terminally tagged by full length luciferase and/or 35S:*Sr62^NLR^* in pCambia1300 backbone. The transfected protoplasts were aliquoted into a 96-well plate with 200 nM D-luciferin/well with 3 biological replicates and then measured with a luminometer (Promega Max NAVIGATOR) every hour for 16 h.

### Transient expression in *Nicotiana benthamiana* and *N*. *tabacum*

*N. tabacum* and *N. benthamiana* plants were grown in a growth chamber at 23 °C with a 16-hour light period and used for agroinfiltration at the 3-to 4-week-old stage. *Agrobacterium tumefaciens* cultures were grown at 28°C overnight with shaking at 200 RPM in Luria-Bertani liquid medium with appropriate antibiotic selections. Cells were pelleted by centrifugation and resuspended in infiltration mix (10 mM MES pH 5.6, 10 mM MgCl_2_, 150 μM acetosyringone). The cells for each construct were diluted to a final concentration of OD_600_ = 0.5 except where stated otherwise and incubated at room temperature for two hours before infiltration. For documentation of cell death, leaves were photographed or scanned 3 to 5 days after infiltration. For Co-IP and western blot, leaves were harvested 24 hours after infiltration.

### *In planta* two-hybrid assays

Plant two-hybrid assays were performed as previously described (*28*). In brief, *N. benthamiana* leaves were co-infiltrated with *Agrobacterium* cultures containing pUAS-Ruby, pGAL4BD and pVP16 fusion constructs at OD_600_ = 0.5. Viral silencing-suppressor protein V2 was used to enhance protein expression. Leaves were photographed 3 to 5 days after infiltration and decolorized by immersing in absolute ethanol overnight. Three leaf disks (0.8 cm diameter) from three independent leaves were pooled into 0.5 ml water to extract betalain overnight and the absorbance of betalain in solution was measured at 538 nm in a spectrophotometer (Biochrom WPA).

### Luciferase luminescence assays in *N. benthamiana*

*N. benthamiana* leaves expressing desired construct combinations were cut into 3 mm x 3 mm pieces 2 days post infiltration and floated in a Petri dish containing water overnight. The leaf pieces were transferred into a 96-well plate with 100 µL of 200 nM D-luciferin/well with 5 replicates for each construct combination. The substrate was infiltrated into the leaf pieces by vacuum for 1 min. Luminescence of an integral of 10 s was measured with a luminometer (Promega Max NAVIGATOR). The *Sr62^NLR^* and *Sr62^TK^* CDS were cloned into pCambia1300-Cluc/Nluc vectors (*31*) respectively with the sites *Kpn* I and *Sal* I to generate constructs encoding Sr62^TK^ with a C-terminal NLUC fusion and Sr62^NLR^ with an N-terminal CLUC fusion. For luminescence imaging of whole *N. bethamiana* leaves, those expressing the desired construct combination for 3 days were pressure-infiltrated with 200 nM D-luciferin, followed by 2 min incubation at room temperature. Leaves were imaged by a VILBER FUSION FX SPECTRA under chemiluminescence detection mode with an exposure time of 3 min.

### Protein extraction, immunoblot, and co-immunoprecipitation

For immunoblot analysis, total proteins from infiltrated leaves were extracted using Laemmli buffer (0.125 M Tris-HCl, pH 7.5, 4% SDS, 20% glycerol, 0.2 M DTT, 0.02% bromophenol blue) and separated by SDS-PAGE and transferred to nitrocellulose membranes. Membranes were blocked with 5% skimmed milk and then incubated in the primary and secondary antibodies, Details for antibodies used in this study were previously described (*47*). Membranes were incubated with Anti-HA-Peroxidase, High Affinity (Roche, reference 12013819001) or mouse anti-GFP (Roche, reference 11814460001) and followed by goat anti-mouse antibodies conjugated with horseradish peroxidase (Biorad, reference 170–5047), or anti-GAL4BD (Sigma-Aldrich, reference G3042) and anti-VP16 (Sigma-Aldrich, reference V4388) antibodies followed by anti-rabbit horseradish peroxidase (Sigma) to detect HA, YFP, GAL4BD and VP16 tagged proteins, respectively. Protein loading was visualized by staining membranes with Ponceau S and protein signals were developed using the SuperSignal West Femto chemiluminescence kit (Pierce).

Co-immunoprecipitation experiments were performed as described (*48*). Total proteins were extracted in 800 μl protein extraction buffer (25 mM Tris-HCl, pH 7.5, 150 mM NaCl, 1 mM EDTA, 10 mM DTT, 1 mM PMSF, and 0.1% NP-40, protease inhibitor cocktail (11836170001, Roche) and 0.5% polyvinylpolypyrrolidone). 600 μL of the supernatant were transferred to a 2.0 mL tube and an equal volume of protein extraction buffer with 5 μL GFP-Trap Magnetic Agarose beads (ChromoTek) were added. Samples were incubated for 60-90 min at 4°C with gentle rotation on a wheel. Magnetic beads were separated and washed once with protein extraction buffer and four times with wash buffer (25 mM Tris-HCl, pH 7.5, 150 mM NaCl, 1 mM EDTA).

### Yeast two-hybrid assays

Yeast two-hybrid assays were performed using the MATCHMAKER GAL4 system (Clontech, USA) according to the manufacturer’s instructions. The coding sequences (CDS) of the full length and truncated *Sr62^NLR^* were cloned into pGADT-7 vector (Clontech, USA, PT3249-5) with the sites *Eco*RI and *Bam*HI. The CDS of effectors were cloned into pGBKT-7 vector (Clontech, USA, PT3248-5) with the sites *Eco*RI and *Bam*HI. The CDS of full length *Sr62^TK^*and its truncated fragments were cloned into pGADT-7 and pGBKT-7 vectors, respectively. For Figure 1E, the *Saccharomyces cerevisiae* HF7c strain was transformed with BD-*Sr62^TK^* and AD-*AvrSr62* fusions in pGBKT-7 and pGADT-7 plasmids respectively. Single colonies were grown in 5 ml of liquid SD medium synthetic dropout (SD) medium without tryptophan (W) and leucine (L) (SD-WL) for 16 h at 30°C. Yeast cultures were adjusted to OD_600_ = 1.0 and then diluted 1/10 and 1/100 times. Aliquots (5 μl) of each sample were inoculated onto SD-WL and selective medium lacking tryptophan, leucine and histidine (SD-WLH) and incubated at 30°C for 3-5 days. For Figure 3B and S14, constructs were co-transformed into yeast strain AH109 and then incubated on SD-Trp/-Leu medium. Three single clones from SD-Trp/Leu medium were transferred to SD-Trp/-Leu/-His/-Ade medium and incubated at 30° C for 5 days to estimate the interactions of different pairs.

### Zahir-1644 PacBio genome and transcriptome sequencing

Seeds of Zahir-1644 1B (longhand 1S^sh^S.1S^sh^L-1BL (DPRM0081, BW_27246; table S3)) were stratified at 4° C for 3 days and placed in soil, 4 plants per pot in 1L pots in greenhouse conditions. Plants intended for DNA extraction were placed in the dark for 36 hours prior to tissue harvest. Two-week old, clean plant material was flash frozen in liquid nitrogen. DNA and RNA extraction, CCS and IsoSeq library preparation, and PacBio sequencing were outsourced to Arizona Genomics Institute, Arizona, USA.

### Zahir-1644 genome assembly, quality assessment and validation

The Zahir-1644 raw sequencing reads from the Pacific Biosciences (PacBio) Sequel II platform were processed using the hifiasm (v.0.19) assembler (*49*). The raw reads were assembled with default parameters without error correction. The assembly was performed on a high-performance computing cluster (at King Abdullah University of Science and Technology, Saudi Arabia) with the following specifications for execution and job management on the cluster: 1 node, 45 tasks per CPU and 500 GB memory. The assembled genome was assessed using the Quality Assessment Tool for Genome Assemblies (QUAST v.5.2) (*50*). Additionally, Benchmarking Universal Single-Copy Orthologs (BUSCO v.5.1) was performed to evaluate the completeness of the Zahir-1644 assembly using the ‘poales’ lineage dataset (*51*). The raw read and assembly metrics can be found in table S1.

### Zahir-1644 isoform processing

Full-length cDNA from Zahir-1644 was sequenced using the PacBio Sequel II system. The raw data from two SMRT cells were processed using the IsoSeq pipeline (v.3; https://github.com/PacificBiosciences/IsoSeq). In brief, the subreads were converted to Circular Consensus Sequencing (CCS) with minimum read quality of 0.9 to ensure retaining high quality reads. The CCS reads were trimmed and adapters and primers were removed using the lima tool. The full-length non-chimeric reads (flnc) were refined using the IsoSeq refine function. After refining, a list of flnc reads were created and used to generate the consensus isoforms by using the IsoSeq cluster function. The high-quality clustered reads were then mapped to the assembled Zahir-1644 genome using the pbmm2 align tool. Finally, the aligned reads were collapsed into a non-redundant or unique set of transcript sequences from the Zahir-1644 assembly using the IsoSeq3 collapse function. The IsoSeq raw and processed read metrics can be found in table S5.

### Seedling assessment of BC_1_F_2_ populations

Seeds were sown in sterilized Mikskaar Professional Potting Soil 70 (Hygrotech, Pretoria, SA) in 10-cm-diameter pots, 25 seeds per pot, and placed in a growth chamber at 25°C. At emergence, seedlings were placed at 22°C (±1°C) in a rust-free greenhouse under natural light. Seedlings were fertilized twice, prior and post inoculation, with 0.2% (w/v) Multifeed-Classic water-soluble fertilizer (Effekto®, NPK Analysis 19:8:16). With their primary leaves fully exposed and the second leaf halfway out, 8-day-old seedlings were inoculated with a (±1 mg/ml) suspension of urediniospores, freshly collected from wheat line Federation*4/Kavkaz, of *Pgt* race PTKST (North American race classification, isolate UVPgt60) in Soltrol® 130 (Chevron Philips Chemical Company, The Woodlands, TX) light mineral oil. Following drying for 30 min in growth cabinets (200 μE/m^2^/s light; 25°C), seedlings were placed in a dark dew chamber at 18°C (±1°C) for 16 h. Upon, removal, seedlings were dried off for 3 h in growth cabinets and returned to the greenhouse under the conditions described above. Seedlings were assessed 12 days post-inoculation using standard infection types (ITs, 0 to 4 scale) (*52*) and considered as either resistant (low ITs) or susceptible (high ITs) per population. The stem rust susceptible entries Line 37-07 and Zahir were included as controls.

### DNA extraction from BC_1_F_2_ plants

Genomic DNA was extracted from freeze-dried leaf material using a cetyltrimethyl ammonium bromide (CTAB) DNA extraction protocol (*53*) by CenGen Pty Ltd (South Africa). The DNA integrity and quantity was assessed with a NanoDrop^®^ Spectrophotometer ND-1000, and by running an aliquot on a 0.8% agarose gel at 50 Volts for 50 minutes alongside a 100 ng Lambda phage control.

### BC_1_F_2_ bulk segregant analysis

Equimolar DNA bulks from susceptible BC_1_F_2_ families (table S6), respectively, were sequenced using the myBaits® Expert Wheat Exome capture panel which targets >250 Mb CDS and CDS-proximate regions of the wheat genome (https://arborbiosci.com/wp-content/uploads/2020/01/myBaitsExpert_WheatExome_Product_Sheet.pdf). Raw whole exome sequencing reads from bulked families were aligned to the wildtype assembly of Zahir-1644 using BWA (v0.7.0) (*54*). The aligned reads were converted to mpileup format using SAMtools (v1.16.0) (*55*) mpileup with parameters, “-BQ0”. The candidate locus was identified using a version of the MutantHunter pipeline (*56*) called MutOatSeq adapted for WGS data (https://github.com/steuernb/oat_mutseq). In short, the Zahir-1644 assembly was scaffolded to the 1S chromosome of the *Ae. sharonensis* 1644 assembly (*15*) using RagTag (*57*). The AGP file produced by RagTag and the single nucleotide variant (SNV) calls for each mutant, filtered to retain only EMS-transition type mutations, were used to establish the order and relative position of SNVs on the 1S chromosome. The region with the highest number of SNVs on the 1S chromosome for each mutant was identified using a sliding window approach. SNVs were counted in a rolling 10 Mbp window with a 5 Mbp overlap across the entire chromosome. SNP counts were normalized between mutants by dividing the number of plants in each bulked susceptible BC_1_F_2_ family. The analysis was performed using Python (v3.11.5) (*58*) and the library Pandas (v2.1.4) (*59*). A plot of the SNV count across the 1S chromosome was generated for each mutant using Matplotlib (v3.8.0) (*60*).

Mapping and variant calling of RNA-Seq, WGS, and exome-capture data to Zahir-1644 Before mapping, the raw reads of RNA-Seq and WGS sequencing reads (table S2) were trimmed to remove adapters and low-quality sequences using trimmomatic (v.0.39) using the following settings: ILLUMINACLIP:2:30:10:2: True LEADING:3 TRAILING:3 MINLEN:36.

After trimming, the reads were mapped using different tools. RNA-Seq reads were mapped to the Zahir-1644 assembly using HISAT2 (v.2.2.1) (*61*). Post-alignment, SAMtools (v.1.16) were used to convert SAM to BAM, sorting and removing duplicates and indexing. Finally, variant calling was performed using bcftools mpileup (v.1.16). For WGS data and exome-capture data, BWA-MEM (v.0.7) was used for mapping against the reference genome Zahir-1644 (*62*). For converting SAM to BAM, removing, and sorting duplicates, indexing, and merging the sorted BAM files SAMtools were used as mentioned above. Variant calling for WGS data was performed using the BCFtools (v1.16) suite and QuickSNP, an in-house Java-based pipeline developed for exome-capture data (https://github.com/steuernb/oat_mutseq). This light-weight SNP caller counts the allele frequency at a position based on the reads mapping at that position. It has filters for coverage and frequency of the reference allele.

### Protein structure modelling

Protein structures and complexes were predicted using AlphaFold (version 2.3.1.) (*32*) installed on the KAUST IBEX cluster. Predictions were run using the in-house Python wrapper, modified to produce additional scores (iPTM, pDockQ, interface pLDDT, number of interface contacts) in addition to pLDDT and pTM (*63*) (fig. S20). For each protein complex prediction, three or more prediction series were run, with 5 models produced for each. The numbers in the Supplementary Tables reporting the model scores correspond to the best-scoring prediction for each complex type (table S9 and table S10). Models were visualized using PyMOL (pymol.org).

Highest scoring Sr62^TK^ Kinase 1:Kinase 2 association within the dimer featured an interaction *in trans*, between the Kinase 1 of one chain and the Kinase 2 of the other chain of the dimer. The topology of this interaction was reproduced exactly by complexes based on separate Kinase 1 and Kinase 2 sequences, reflecting the case where the Kinase 1 and Kinase 2 association was observed in planta (fig. S21A-B and table S9). For the association between Sr62^NLR^ and Sr62^TK^ Kinase 2, two possible interaction topologies were produced by AlphaFold with similar frequencies and interactions scores (fig. S1C-D and table S9). In both, the interaction site was located on the Sr62^NLR^ NB-ARC2. However, in one topology, Kinase 2 docked onto the NB domain, whereas in the other, onto the ARC domain. Given that Sr62^TK^ binds as a homodimer to Sr62^NLR^, it is possible that each of the two Kinase 2 domains binds to one Sr62^NLR^ site, forming a 2:1complex.

In AlphaFold predictions the effectors bind with the same topology to monomeric or dimeric Sr62^TK^, preserving the key interactions between the Kinase 1 β-finger and the β-sheet formed by β-strands contributed from the effector’s N and C-termini (fig. S21 E to K and fig S22, and table S9). In several models, the conserved lysine 6 from the N-terminal extension of the effectors reaches back into the Sr62^TK^ active site (fig. S22).

### Identifying Sr62^NLR^ natural sequence variation in *Triticum* and *Aegilops*

Seventy-six publicly available genomes spanning the *Triticum* and *Aegillops* genera were analyzed to identify *Sr62^NLR^* homologs. For each genome, candidate exons and gene homolog loci were identified with the custom-built Semi-Automatic BLAST Annotation Toolkit (SABAT) Exon Annotation Nextflow Pipeline (https://github.com/Orpowell/SABAT_NF) with the following parameters: --b2b_exons 3, --b2b_coverage 0.85, and --b2b_locus_size 4000 and the CDS of *Sr62^NLR^*from *Aegillops sharonensis* accession 1644. Predicted *Sr62^NLR^*gene loci and exons for each genome were visualized and manually curated using IGV (*64*) to ensure all potential gene loci had been identified. The SABAT (https://github.com/Orpowell/SABAT) command *assemble-locus* was used to assemble candidate gene loci into CDS and protein sequences with the parameter: *-f 30*. Candidate protein sequences were clustered into groups with 100% sequence identity using CD-HIT (*65*) with the parameter: *c -1*. A multiple sequence alignment was generated with a representative sequence from each cluster and the Sr62^NLR^ sequence from *Ae. sharonensis* accession 1644 using MUSCLE (*66*) (v5.1) and visualized using Python and the library Biotite (*67*) (v 0.39.0). Natural sequence variation in Sr62^NLR^ homologs was mapped onto the predicted AlphaFold structure of the protein using a custom Python script and visualized in PyMol (http://www.pymol.org/pymol).

### Identifying Sr62^TK^ natural sequence variation in *Triticum* and *Aegilops*

The same 76 publicly available *Triticum* and *Aegilops* genomes as above were analyzed to identify homologs of *Sr62^TK^*. The SABAT Exon Annotation Pipeline was run with the following parameters: --b2b_exons 11, --b2b_coverage 0.85, and --b2b_locus_size 7500 and the CDS of *Sr62^TK^*from *Ae. sharonensis* accession 1644. Predicted gene loci and exons were visualized and manually curated using IGV (*64*). In several cases, gene loci could not be predicted by the pipeline but a candidate homolog could be identified from the predicted exons. The position of each candidate *Sr62^TK^* homolog was compared to the predicted position of the *Sr62^NLR^* homolog in the same genome (see above) to ensure that the predicted gene loci were homologs of *Sr62* and not the highly similar *WTK3* (*Pm24*/*Rwt4*). Candidates that were not within 30 kbp of a candidate *Sr62^NLR^* homolog were discarded.

The SABAT command *assemble-locus*, with default parameters, was used to assemble candidate genes from genomes with predicted gene loci into CDS and protein sequences. Genomes with no predicted gene loci were manually curated to identify exons belonging to the gene and assembled using the SABAT command *assemble-exons* with default parameters. Candidate protein sequences were clustered into groups with 100% sequence identity using CD-HIT with the parameter: *c -1*. A multiple sequence alignment was generated with a representative sequence from each cluster and the Sr62^TK^ sequence from *Ae. sharonensis* accession 1644 using MUSCLE (v5.1) and visualized using Python and the library Biotite (*67*) (v 0.39.0). Natural sequence variation in Sr62^TK^ homologs was mapped onto the predicted AlphaFold structure of the protein using a custom Python script (https://github.com/Orpowell/sr62_analysis) and visualized in PyMol. In *T. diccocoides* accession 210222, the final exon was not predicted by the SABAT exon annotation pipeline. Instead, the exon sequence was manually identified in IGV and added to the protein sequence.

### Calculating percentage identity of Sr62^NLR^ homologs

The experimentally derived NLR sequences from chromosome 1S of *Aegilops sharonensis* accession 1644 and chromosome 1D of *Triticum aestivum* cvs. Zahir and Fielder, and the chromosome 1B of cv. Fielder (predicted by SABAT, see above) were aligned using MUSCLE (v5.1) with default parameters. The resulting alignment was converted into a percentage identity matrix (PIM) using Clustal Omega (v1.2.4) (*68*) with following command: *clustalo -i alignment.fasta --percent-id --distmat-out=pim.txt --full –force* .

## Acknowledgments

This research used the Ibex HPC cluster and the Shaheen Supercomputer managed by the Supercomputing Laboratory at King Abdullah University of Science and Technology (KAUST). We are grateful to Zhiyong Liu (Chinese Academy of Sciences) and Peter Brodersen (Copenhagen University) for helpful discussions, Tobin Florio (www.Flozbox-Science.com) for figure artwork, Haitao Cui (Shandong Agricultural University) for Golden Gate plasmids, Burkhard Steuernagel (JIC) for bioinformatics advice, Ana Belén Perrera Rodríguez (KAUST) for greenhouse assistance, Guo (Cherry) Huijuan (Novogene) and Dario Copetti (Arizona Genomics Institute) for NGS services, and Debbie Snyman and Renée Prins (CenGen Pty Ltd.) for DNA extractions.

## Funding

KAUST baseline and awards CRG10-2021-4735 and CRG11-2022-5087 (BBHW).

KAUST baseline and awards FCC/1/1976-33 and REI/1/4446-01 (STA).

The National Research Foundation, SARChI chair UID 8464 (WHPB).

The Lieberman-Okinow Endowment at the University of Minnesota (BJS).

The National Natural Science Foundation of China (31830077) (DT).

CSIRO Research Office, OD-213047, OD-225629, OD-227545

CSIRO SynBio Future Science Platform OD-206702

## Author contributions

Conceptualization: RC, JC, MAO, ORP, TA, MF, JS, TV, PND, BBHW.

Investigation: RC, JC, MAO, ORP, TA, KG, YLW, CB, JS, YX, GZ, CG, GY, SG, OM, WHPB, WBM, STA

Visualization: RC, JC, MAO, ORP, KG, YLW, JS, STA, PND, BBHW

Funding acquisition: MAA, MF, JS, TV, DT, LJ, BJS, WHPB, STA, PND, BBHW

Project administration: DT, LJ, BJS, WHPB, PND, BBHW Writing – original draft: RC, JC, MAO, ORP, STA, PND, BBHW

Writing – review & editing: RC, JC, MAO, ORP, TA, MAA, MF, JS, TV, STA, PND, BBHW “Authors are listed by institution and surname, except for first seven and last three authors.”

## Data and materials availability

Selected wheat cultivars and EMS-derived mutants are available from SeedStor (table S3). Bespoke code is available from GitHub repositories https://github.com/Orpowell/SABAT, https://github.com/Orpowell/SABAT_NF, https://github.com/Orpowell/sr62_analysis and https://github.com/steuernb/oat_mutseq. Sequence data is available from ENA project ID PRJEB74232, except for the wheat protoplast RNA-Seq data which can be accessed at the CSIRO data access portal (https://doi.org/10.25919/j0vh-ve59). The *Sr62^NLR^* and *AvrSr62* variant sequences and annotations are available from NCBI (GenBank accession number PP537390 and Bioproject PRJNA516922, respectively). The Zahir-1644 genome assembly can be downloaded from doi:10.5061/dryad.hdr7sqvr9.

## Supplementary Figures

**Fig. S1.**
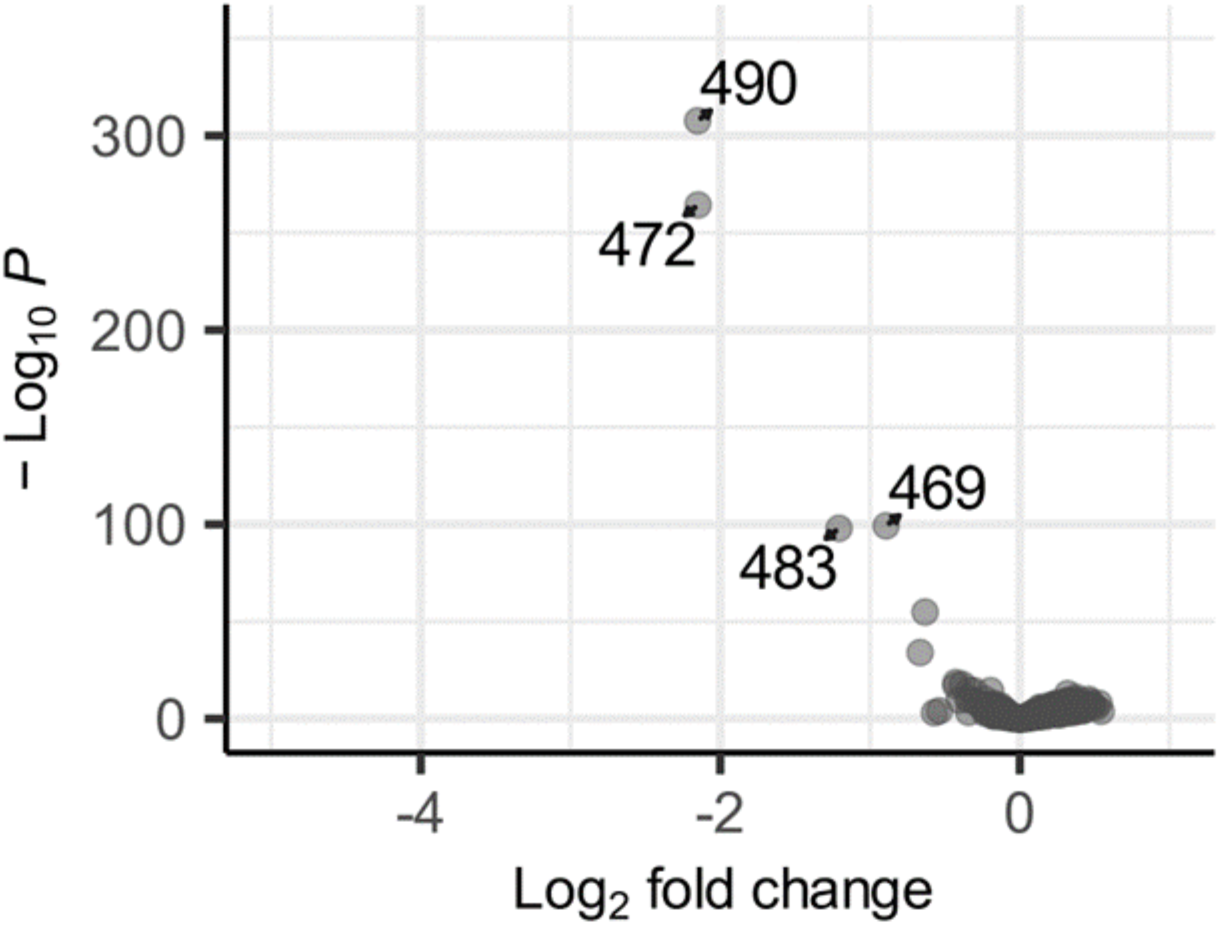
*Pgt* effector library screen with *Sr62^TK^*. Differential gene expression analysis of a pooled stem rust effector library co-transformed into wheat protoplasts with *Sr62^TK^* compared to the empty vector. Graph shows volcano plot of differential expression (X-axis) versus adjusted P value (Y-axis) for each effector construct (dots). Effector gene constructs showing significantly reduced expression (red dots) within each treatment are labelled with their library ID number.

**Fig. S2.**
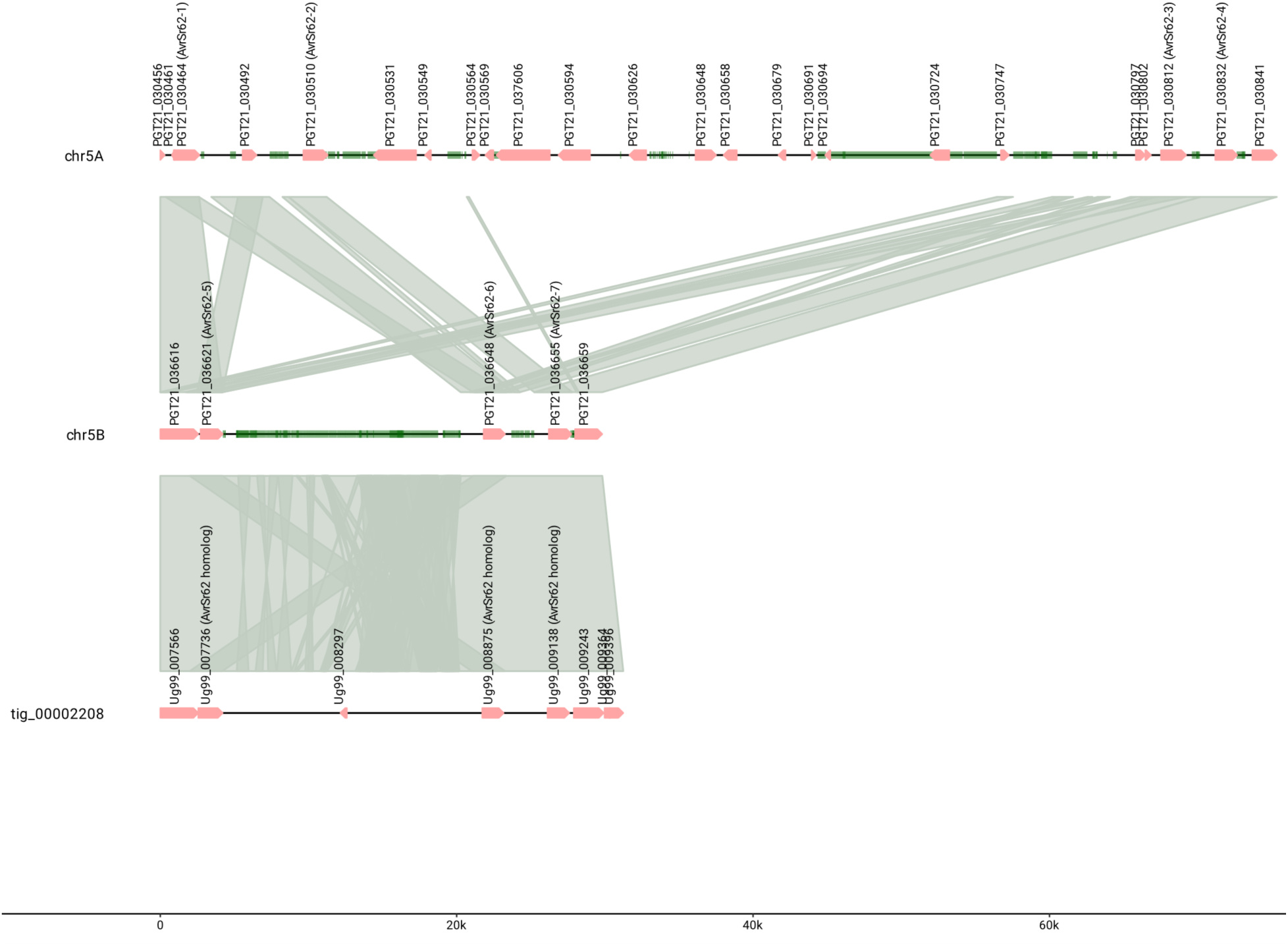
Synteny plot of the *AvrSr62* locus region on the 5A and 5B chromosomes of *Pgt*21-0 and on the C haplotype of Ug99 (tig_00002208). Annotated genes are shown in pink and labelled with their gene ID and repetitive sequences are shown in green. Highly similar sequences between haplotypes are indicated by grey shading. The Ug99 strain contains an identical A haplotype to Pgt21-0, while the C haplotype is similar to the Pgt21-0 B haplotype and contains identical gene sequences to AvrSr62-5, -6 and -7.

**Fig. S3.**
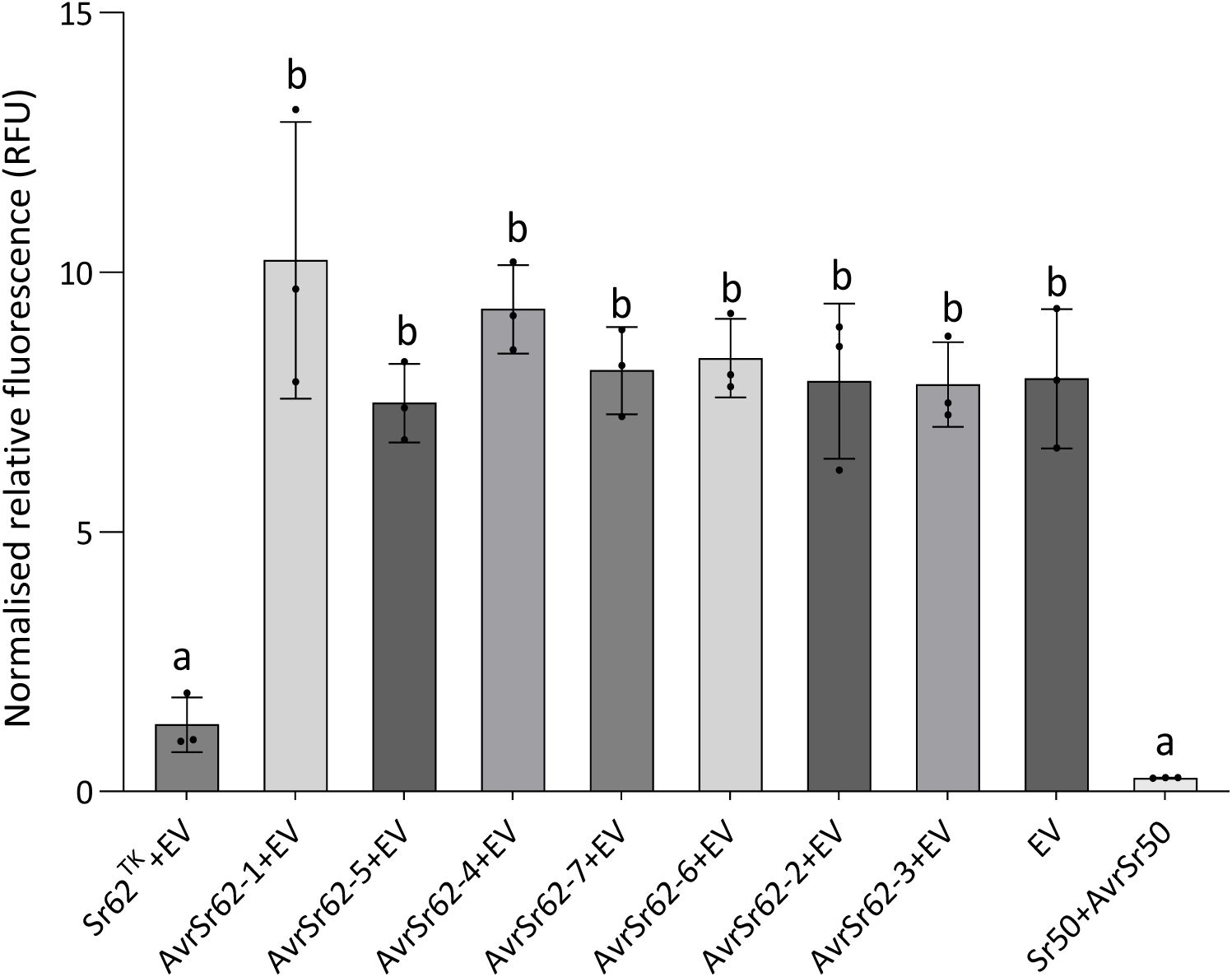
*Sr62^TK^* autoactivity in protoplasts. Protoplasts from wheat cv. Fielder were co-transformed with YFP and individual *AvrSr62* genes or *Sr62^TK^* alone as indicated. Co-expression of Sr50 with AvrSr50 was used as a positive control. YFP fluorescence (y-axis) was measured after 24 hours. Results represent the means of three biological replicates (dots) with error bars indicating the SE. Samples marked by identical letters in the plots do not differ significantly (P < 0.05; ANOVA, posthoc Tukey test).

**Fig. S4.**
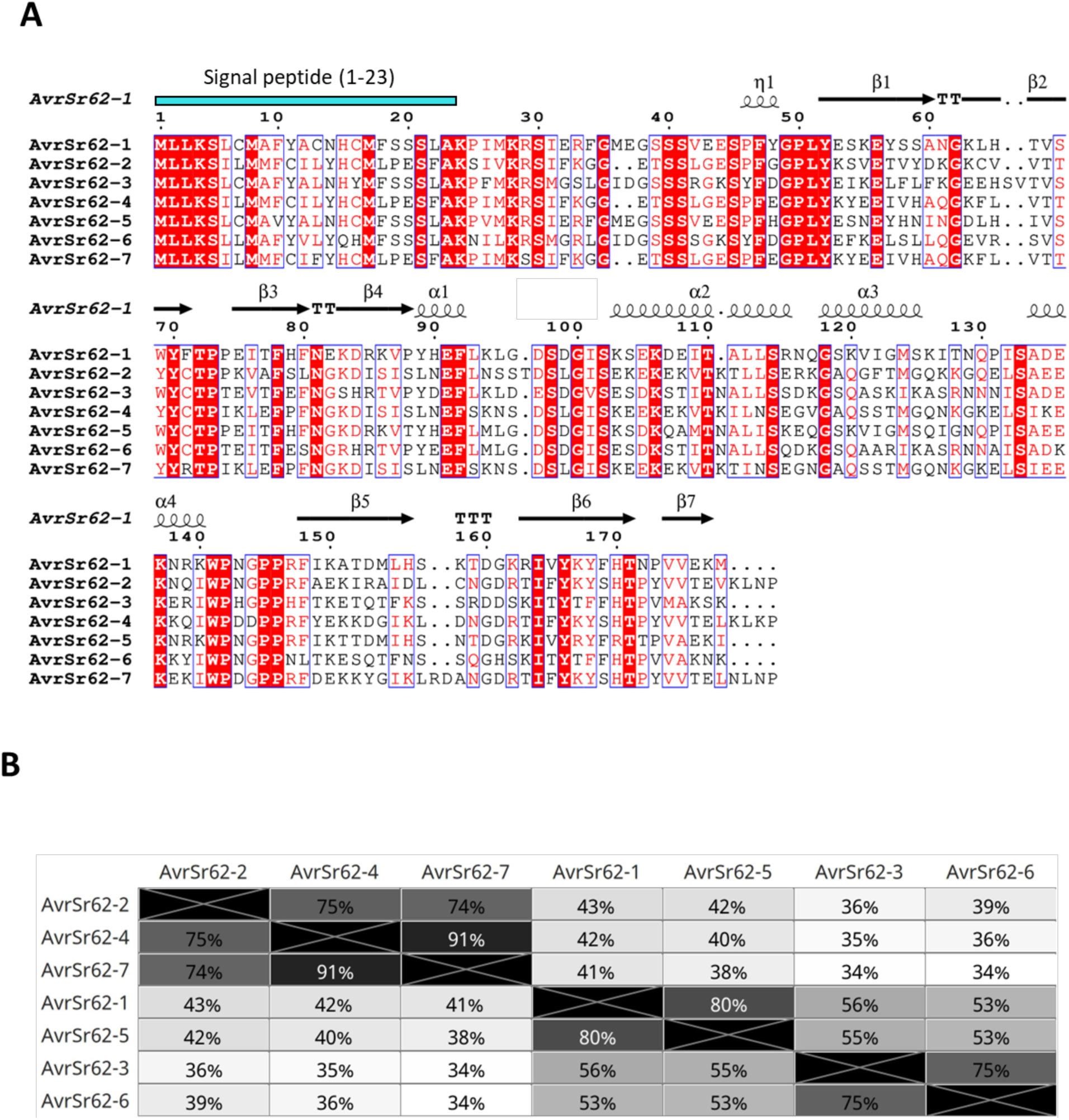
Sequence alignment of all seven AvrSr62 effector candidates. (**A**) Multiple sequence alignment of the full-length sequence of the seven AvrSr62 proteins generated using ESPript3 (*69*). The secondary structure elements of the AlphaFold2 prediction of AvrSr62-1 are displayed on the protein alignment, and signal peptide is indicated based on SignalPv4.1 prediction (-t euk -u 0.34 -U 0.34). (**B**) Pairwise amino acid identities between AvrSr62 proteins.

**Fig. S5.**
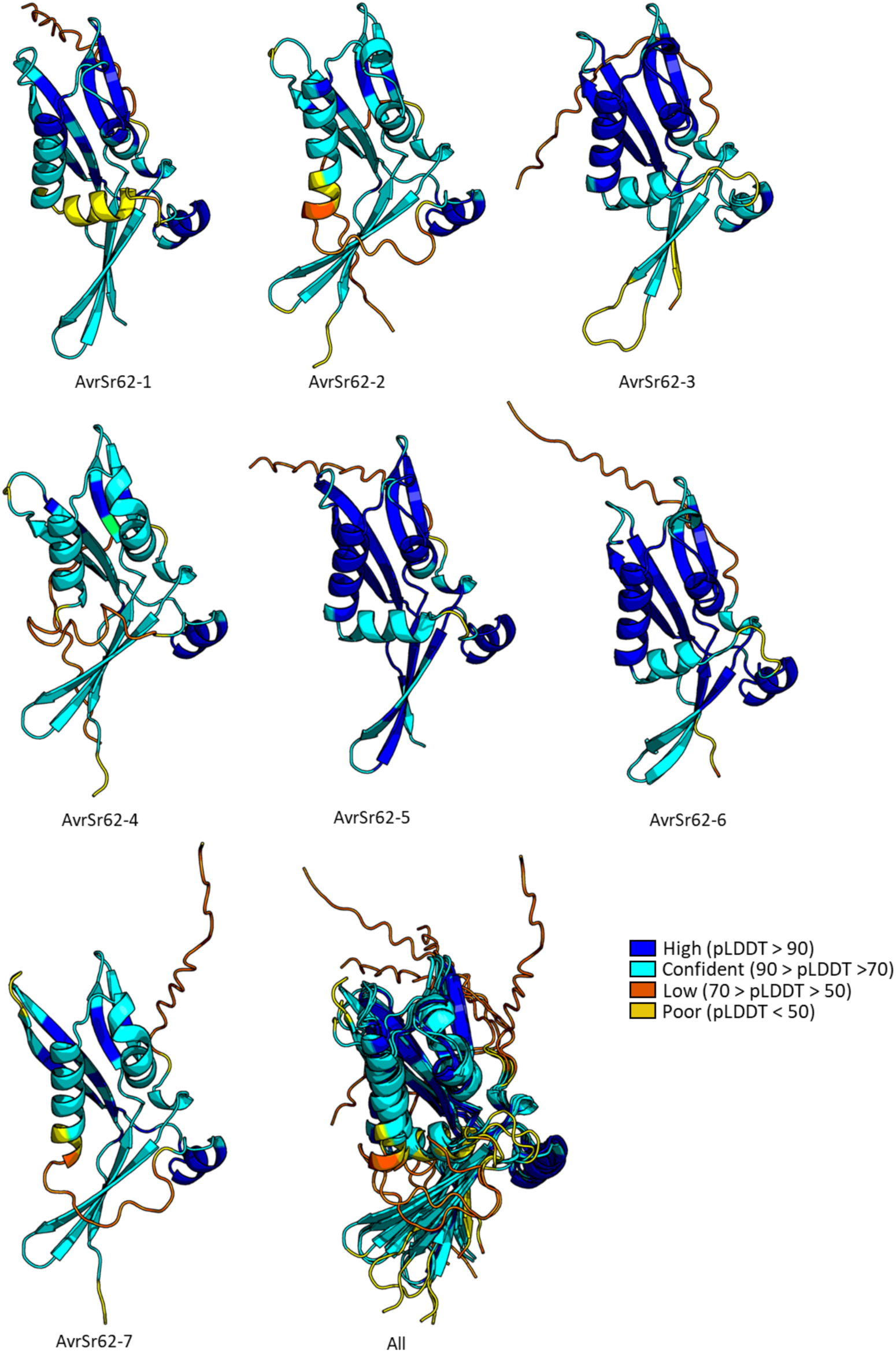
AlphaFold2 predictions of AvrSr62 family. Protein cartoon representations are colored according to confidence of the prediction (pLDDT) as indicated in the key, where blue is the most confident and yellow the least confident.

**Fig S6.**
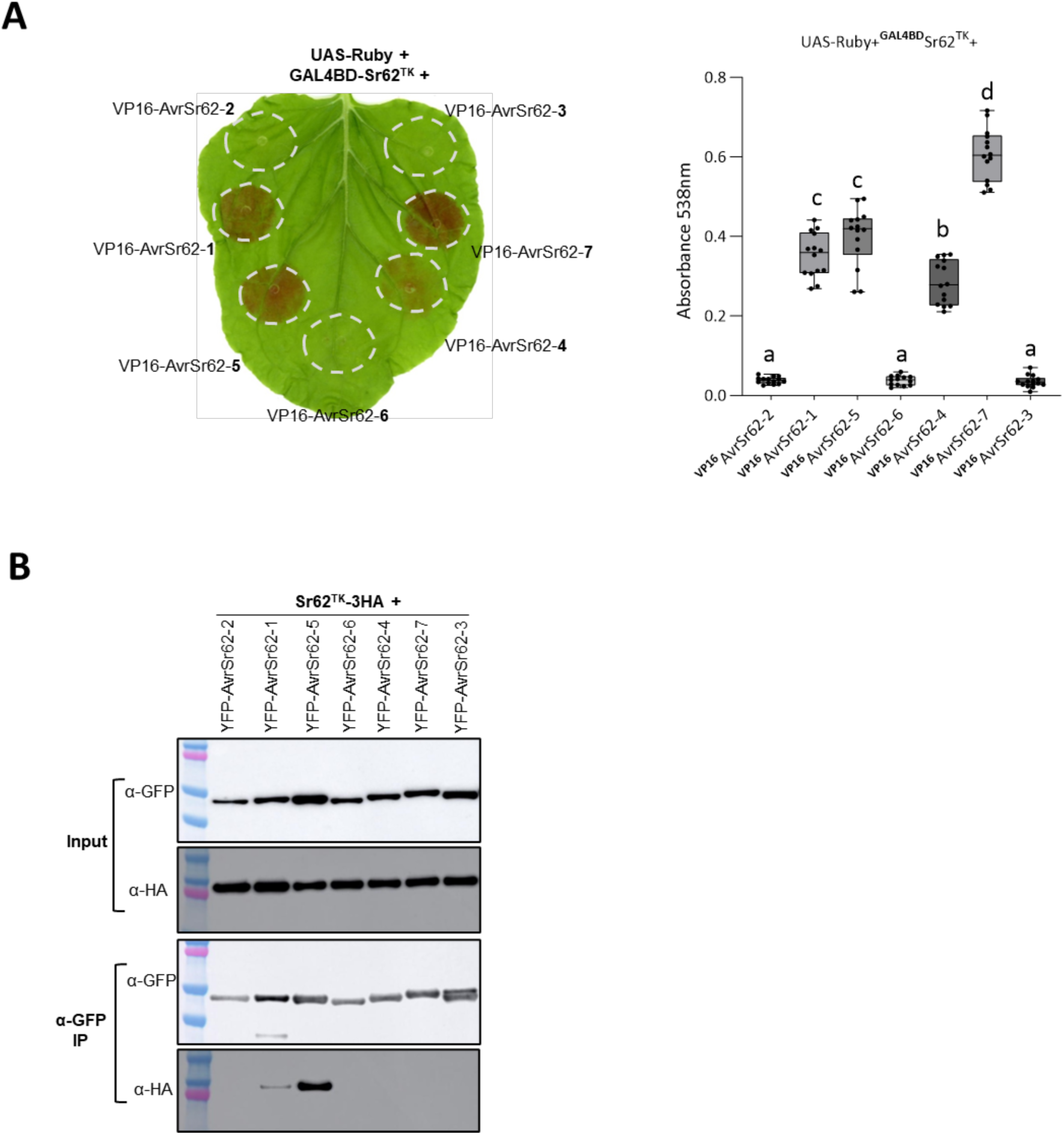
Sr62^TK^ interacts with AvrSr62 effectors. (**A**) *In planta* two hybrid assay. Betalain accumulation was observed in leaves expressing UAS-Ruby with Sr62^TK^ fused to BD and AvrSr62 variants 1 to 7 fused to VP16. Leaves were photographed 3 days after infiltration. Extracted betalain was quantified by absorbance at 538 nm (Right). Graph indicates results box and whisker plots with at least five biological replicates (dots). Samples marked by identical letters in the plots do not differ significantly (P < 0.05; ANOVA Tukey test). (**B**) Co-IP assay. YFP-tagged AvrSr62 variants 1 to 7 were transiently co-expressed in *N. benthamiana* with HA-tagged Sr62^TK^. Total protein extracts were immunoprecipitated with anti-GFP (beads), followed by immunoblotting with anti-GFP, and anti-HA antibodies.

**Fig. S7.**
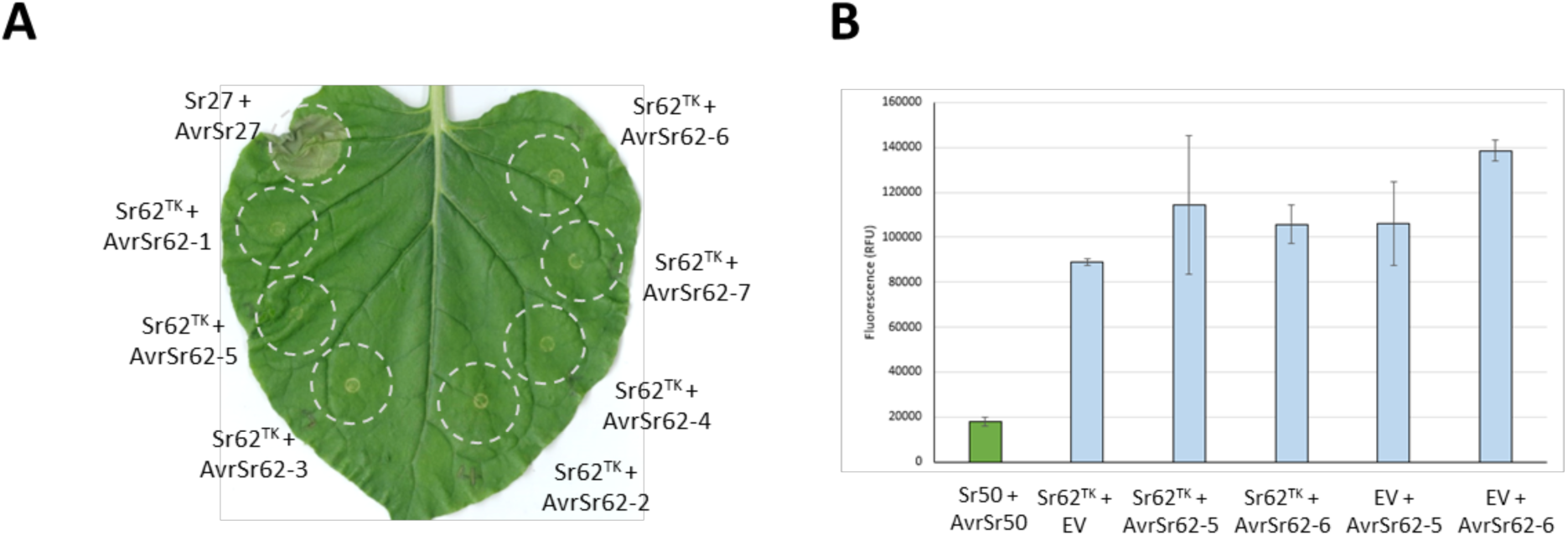
Co-expression of *Sr62^TK^* and *AvrSr62* in *N. benthamiana* leaves or oat protoplasts does not induce cell death. (**A**) *N. benthamiana*. *Sr62^TK^* fused to 3xHA tag at the C-terminus was co-expressed with AvrSr62 effectors with a N-terminal YFP tag. *Sr27*-YFP co-expression with YFP-*AvrSr27-1* was used as positive control. (**B)** Oat (*Avena sativa*) cv Swan. *Sr62^TK^* and *AvrSr62-5* or *-6* were expressed singly or in combination in oat protoplasts as indicated and cell viability measured by fluorescence of a co-transformed YFP reporter gene. Co-expression of *Sr50* and *AvrSr50* was used as positive control.

**Fig. S8.**
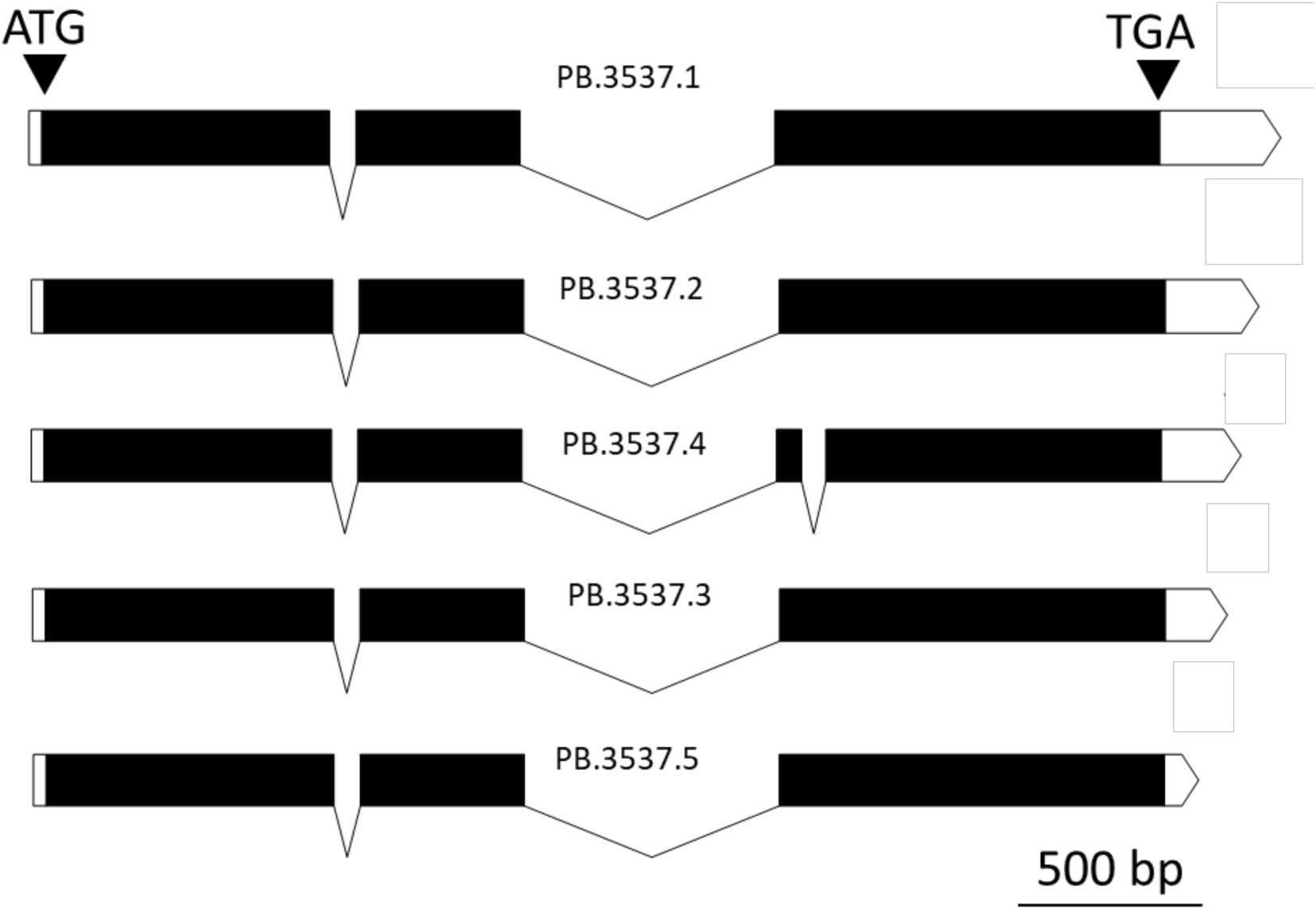
Full length transcripts of *Sr62^NLR^* identified in Zahir-1644 by PacBio IsoSeq. The five transcripts (PB.3537.1 to PB.3537.5) were recovered from 9,264,998 IsoSeq reads obtained from a seedling leaf transcriptome (table S5). Black and white boxes represent predicted translated and untranslated exons, respectively, while black connecting lines represent introns. The start (ATG) and stop (TGA) codons are marked with black arrows.

**Fig. S9.**
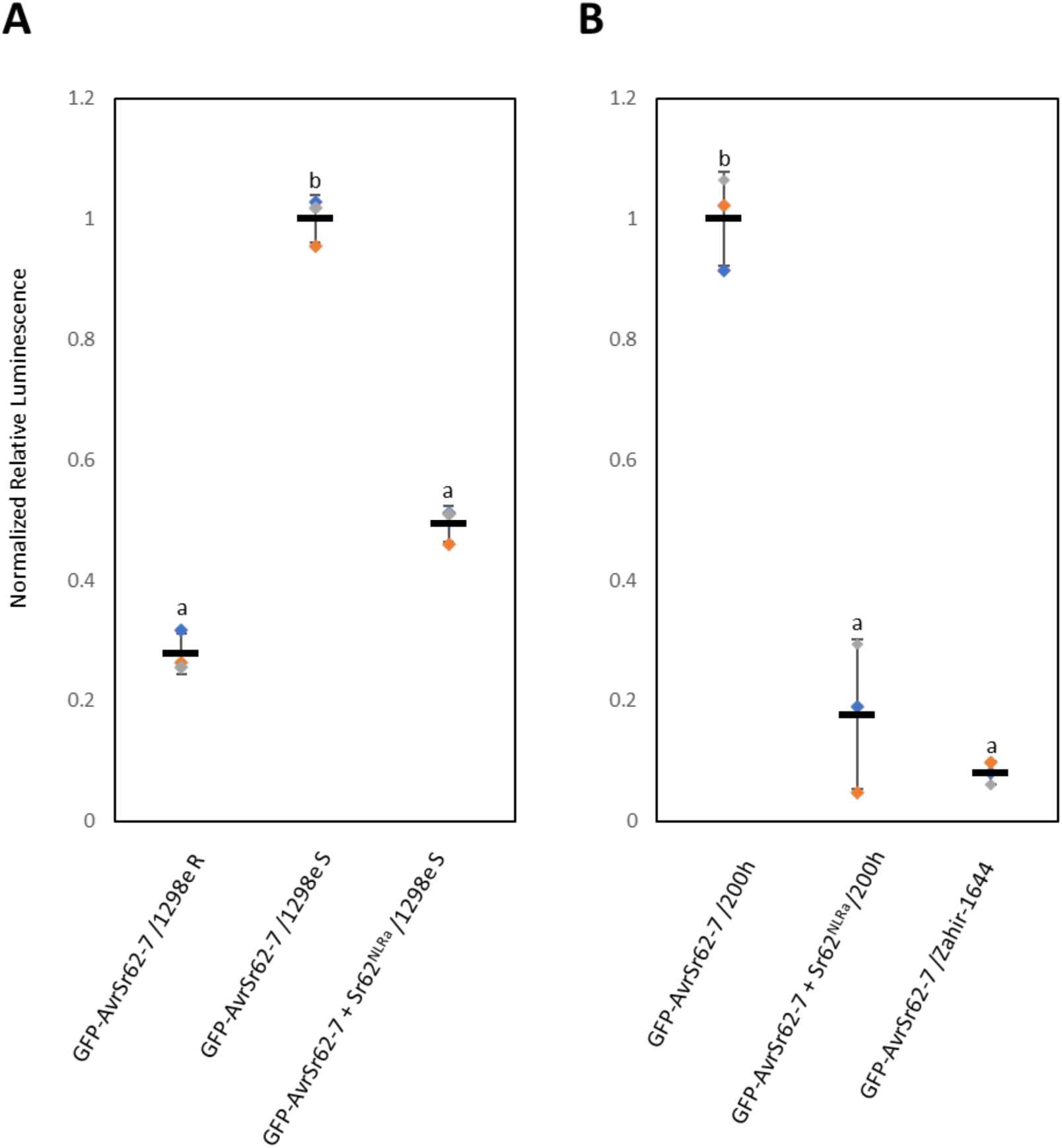
Co-expression of *Sr62^NLR^*and *AvrSr62-7* in Zahir-1644 mutant protoplasts. Complementation of *AvrSr62*-induced cell death in Zahir-1644 *Sr62^NLR^* 1298e and 200h mutant wheat protoplasts by co-transfection with *Sr62^NLR^*. The resistant Zahir-1644 parent (right graph) or a homozygous resistant BC1F2 segregant (1298e R, left graph) were used as a positive control. Luciferase activity was measured after 16 hours. The relative luminescence signal of each sample was normalized to GFP-AvrSr62-7 /1298e S (**A**), or GFP-AvrSr62-7 /200h (**B**). Results represent the means of three biological replicates (dots) with error bars indicating the SE. Samples marked by identical letters in the plots do not differ significantly (P < 0.05; ANOVA Tukey test).

**Fig. S10.**
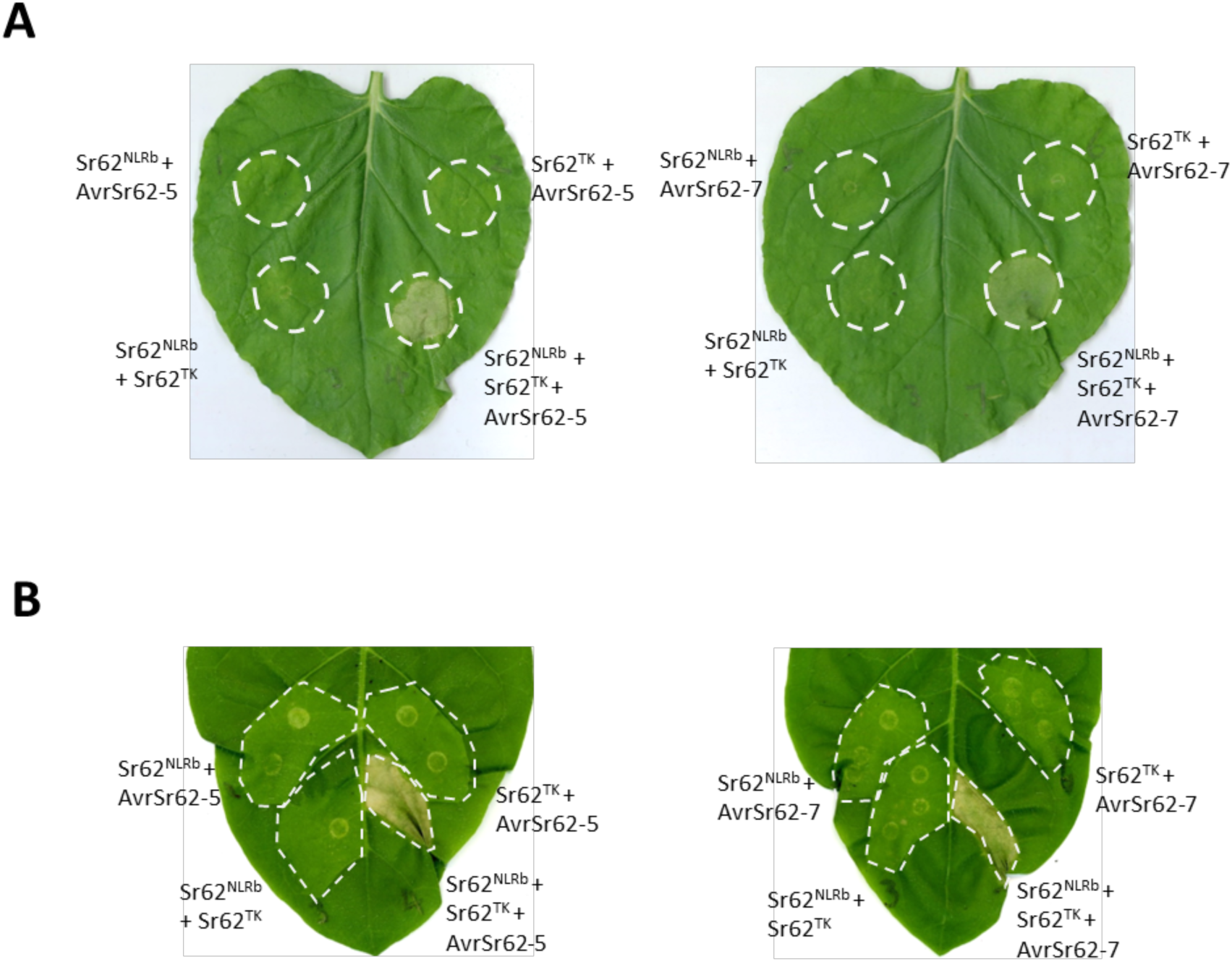
Sr62^NLR^ induces cell death with Sr62^TK^ and AvrSr62 in tobacco leaves. (**A**) *N. benthamiana*. (**B**) *N. tabacum*. *Sr62^NLRb^*and *Sr62^TK^* with C-terminal HA tags and AvrSr62-5 and AvrSr62-7 with N-terminal YFP-tags were co-expressed in leaves by Agroinfiltration in the indicated combinations. Leaves were visualized at three days post infiltration.

**Fig. S11.**
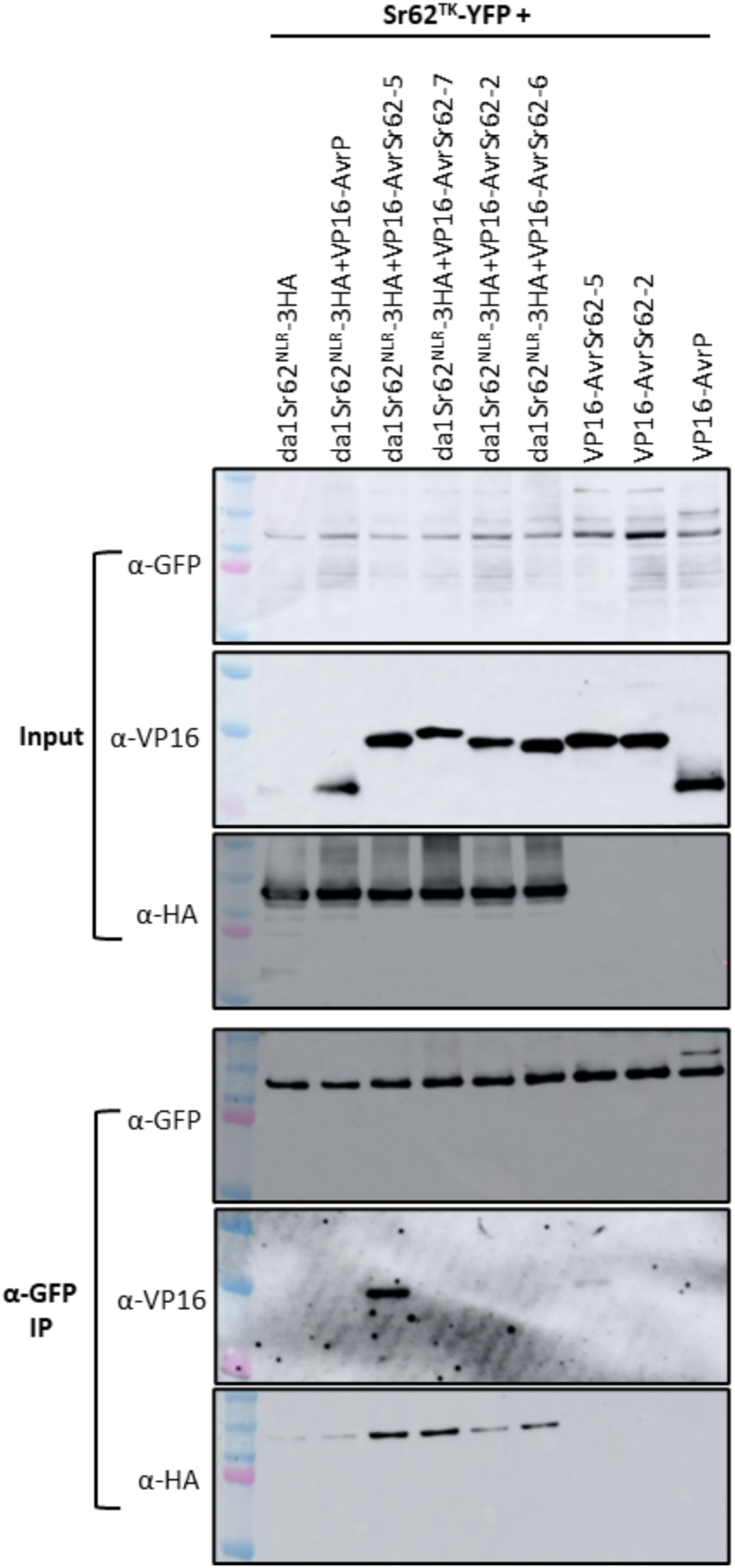
AvrSr62 enhances the interaction between Sr62^TK^ and Sr62^NLR^ in Co-IP. YFP-tagged *Sr62^TK^* was transiently co-expressed in *N. benthamiana* with HA-tagged dα1*Sr62^NLRb^*, VP16-tagged *AvrSr62-2*, *AvrSr62-5*, *AvrSr62-6*, and *AvrSr62-7*. Total protein was extracted and subjected to immunoprecipitation with anti-GFP beads, followed by immunoblotting with anti-HA, anti-VP16, and anti-GFP antibodies.

**Fig. S12.**
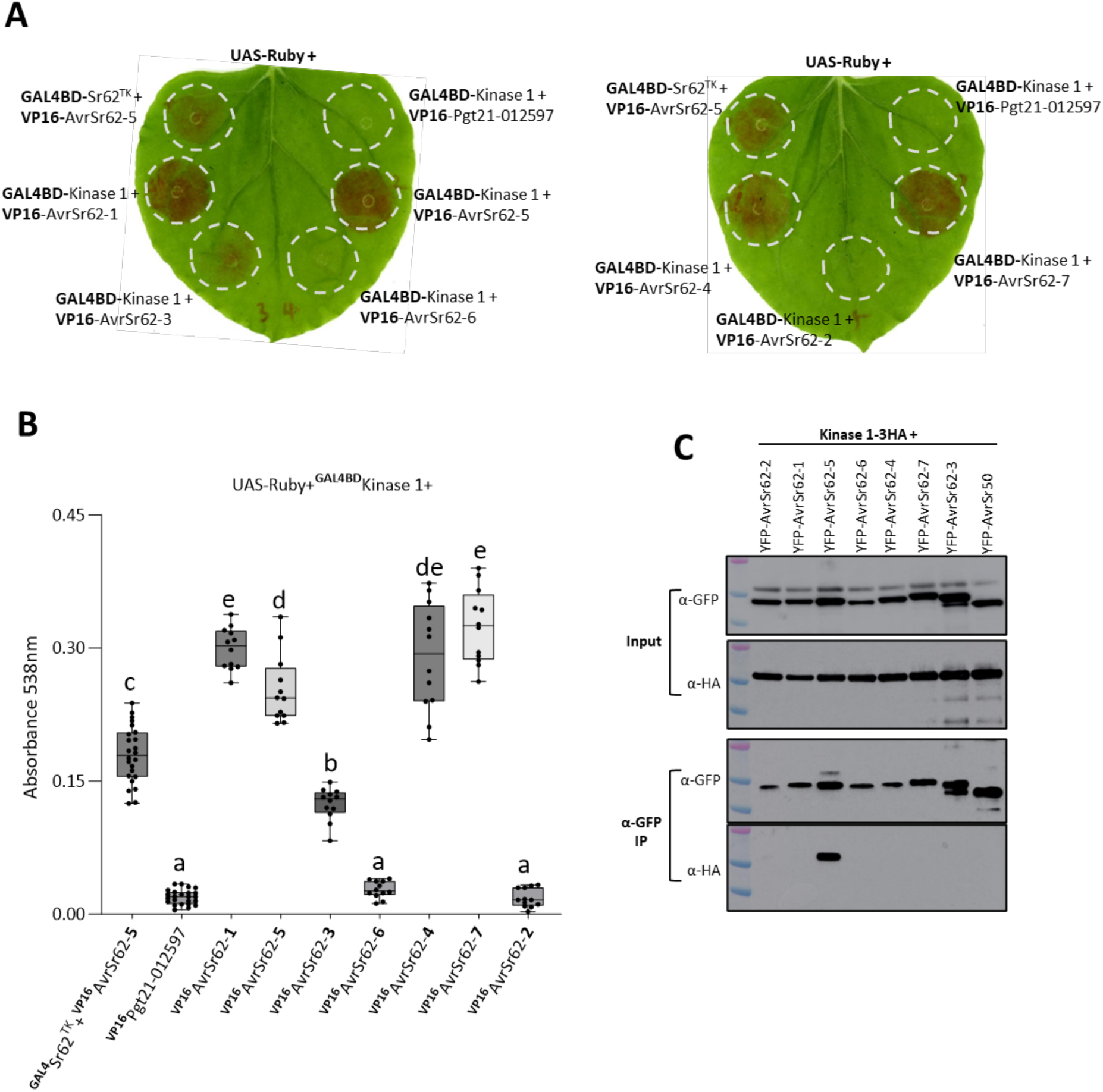
Sr62^TK^ Kinase 1 domain interacts with AvrSr62 effectors. (**A**) Betalain accumulation was observed in leaves expressing UAS-Ruby with Kinase 1 fused to BD and different AvrSr62 fused to VP16. Sr62^TK^ fused to BD co-expressed with AvrSr62-5 fused to VP16 was used as a positive control. Pgt21-012397 fused to VP16 co-expressed with Kinase 1 was used as a negative control. (**B**) Betalain accumulation of each co-expression pair in (A) was quantified and graph indicates the results as box and whisker plots with at least five biological replicates (dots). Samples marked by identical letters in the plots do not differ significantly (P < 0.05; ANOVA Tukey test). (**C**) HA tagged Sr62^TK^ Kinase 1 domain was transiently co-expressed in *N. benthamiana* with YFP tagged AvrSr62 variants 1 to 7. Total protein was extracted and subjected to immunoprecipitation with anti-GFP beads, followed by immunoblotting with anti-HA and anti-GFP antibodies. YFP-tagged AvrSr50 was used as a negative control.

**Fig. S13.**
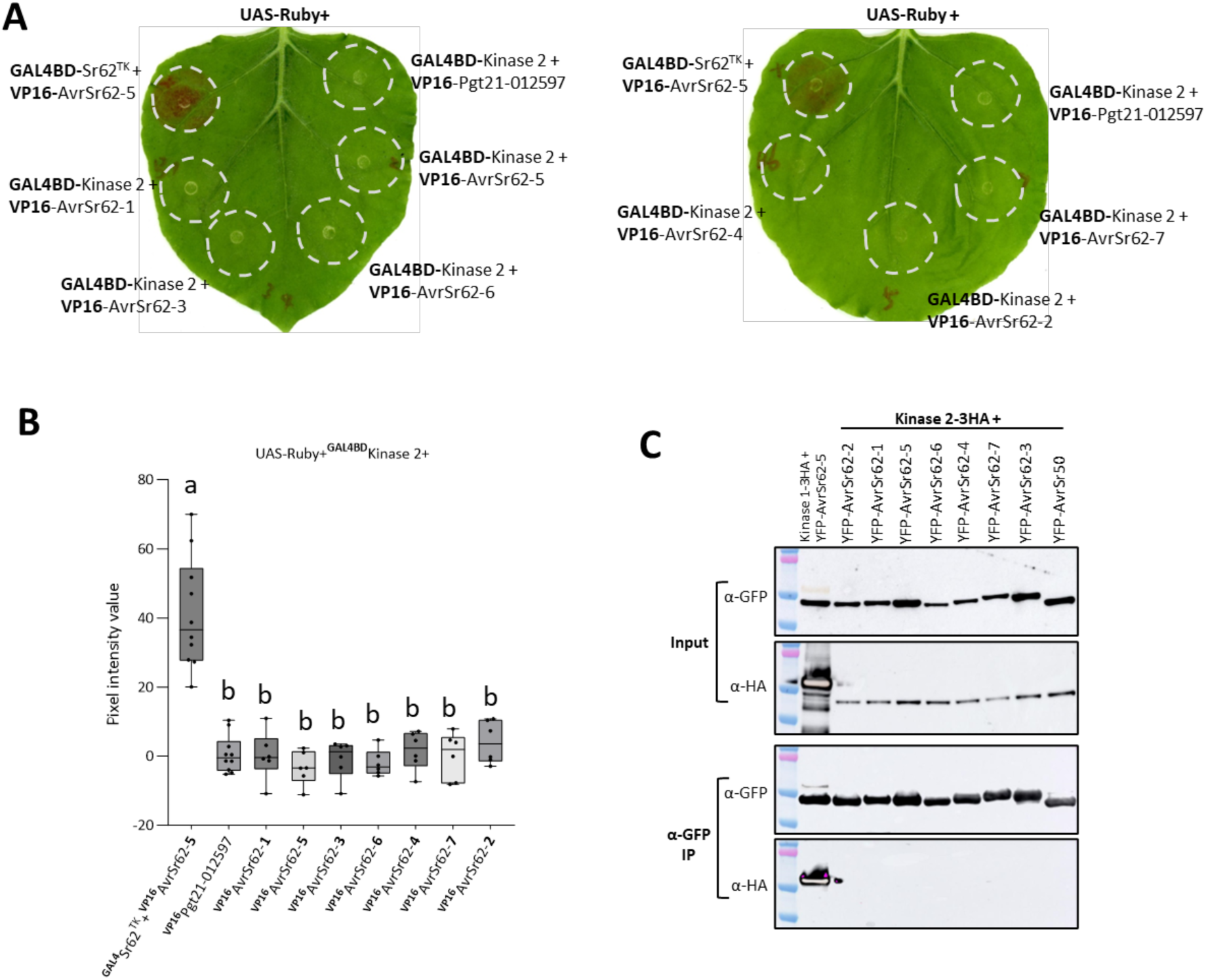
Sr62^TK^ Kinase 2 domain does not interact with AvrSr62. (**A**) Betalain accumulation was observed in leaves expressing UAS-Ruby with Kinase 2 fused to BD and AvrSr62 variants 1 to 7 fused to VP16. Sr62^TK^ fused to BD co-expressed with AvrSr62-5 fused to VP16 was used as a positive control. Pgt21-012397 fused to VP16 co-expressed with Kinase 2 was used as a negative control. (**B**). Betalain accumulation of each co-expression pair in (A) was quantified by pixel intensity values calculated via the ImageJ integrated density function as described (*1*). Graph shows results as box and whisker plots with at least five biological replicates (dots). Samples marked by identical letters in the plots do not differ significantly (P < 0.05; ANOVA Tukey test). (**C**) HA-tagged Sr62^TK^ Kinase 2 domain was transiently co-expressed in *N. benthiamiana* with YFP-tagged AvrSr62 variants 1 to 7. Total protein was extracted and subjected to immunoprecipitation with anti-GFP beads, followed by immunoblotting with anti-GFP and anti -HA antibodies. YFP tagged AvrSr50 was used as a negative control. Co-IP of Kinase 1 fused to HA tag and AvrSr62-5 fused with YFP was used as a positive control.

**Fig. S14.**
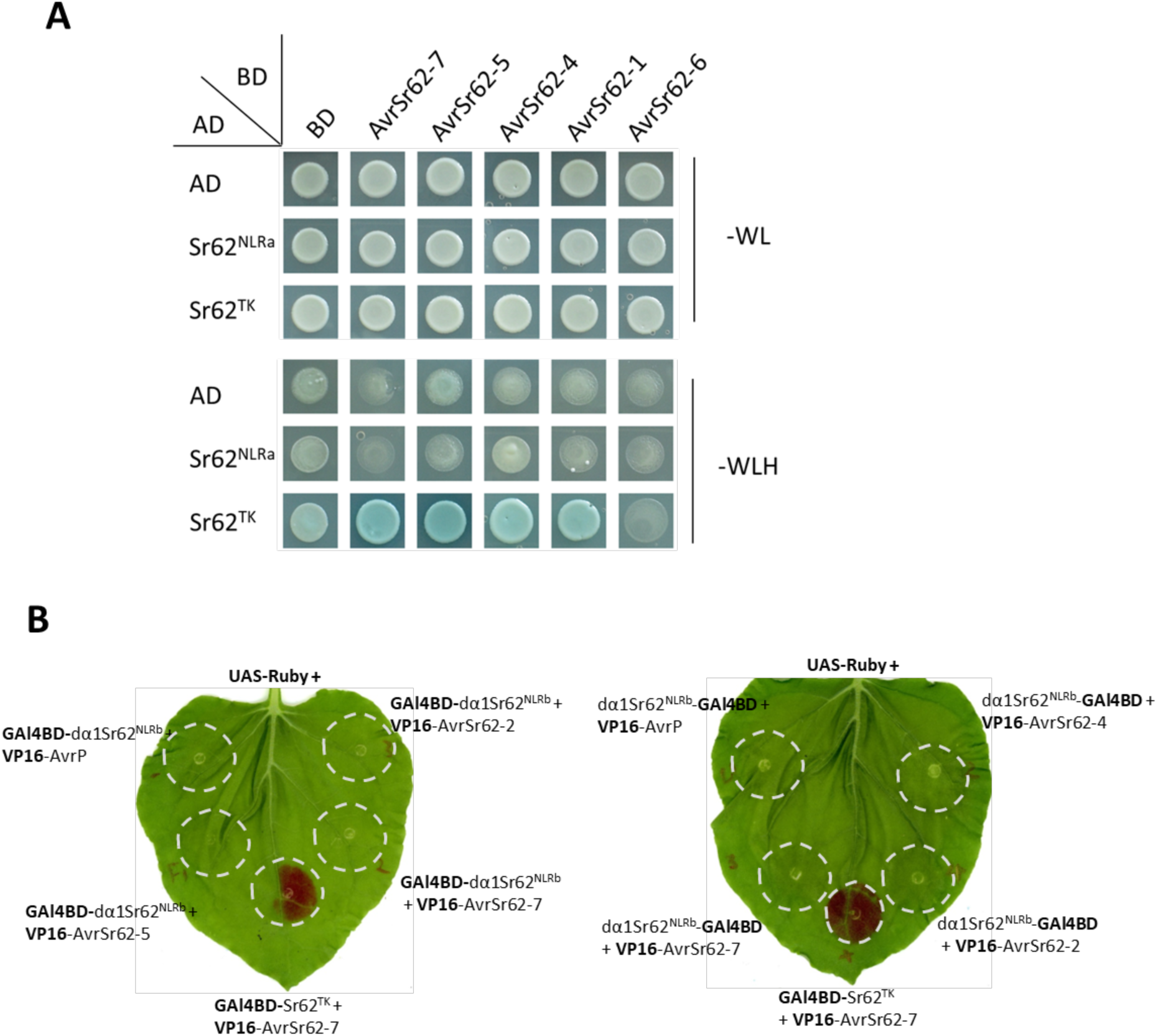
Sr62^NLR^ does not interact with AvrSr62 effectors. (**A**) The coding sequences of *Sr62^TK^* and *Sr62^NLRa^* were fused to the Gal4 transactivation domain (AD). The coding sequences of *AvrSr62-4*, *AvrSr62-5, AvrSr62-6,* and *AvrSr62-7* were fused to the Gal4 DNA binding domain (BD). Different pairs of constructs were co-transformed into AH109. A 10 μL suspension (OD_600_ = 0.5) of each co-transformant was dropped onto synthetic dropout (SD) medium lacking histidine (H) and tryptophan (W), and SD medium lacking histidine (H), leucine (L), and tryptophan (W) with X-Gal. Photographs were taken 3 days after plating. Sr62^TK^ was used as a positive control. (**B**) Betalain accumulation was observed in leaves expressing UAS-Ruby with dα1Sr62^NLRb^ fused to BD and different AvrSr62 variants fused to VP16. Sr62^TK^ fused to BD co-expressed with AvrSr62-7 fused to VP16 was used as a positive control.

**Fig. S15.**
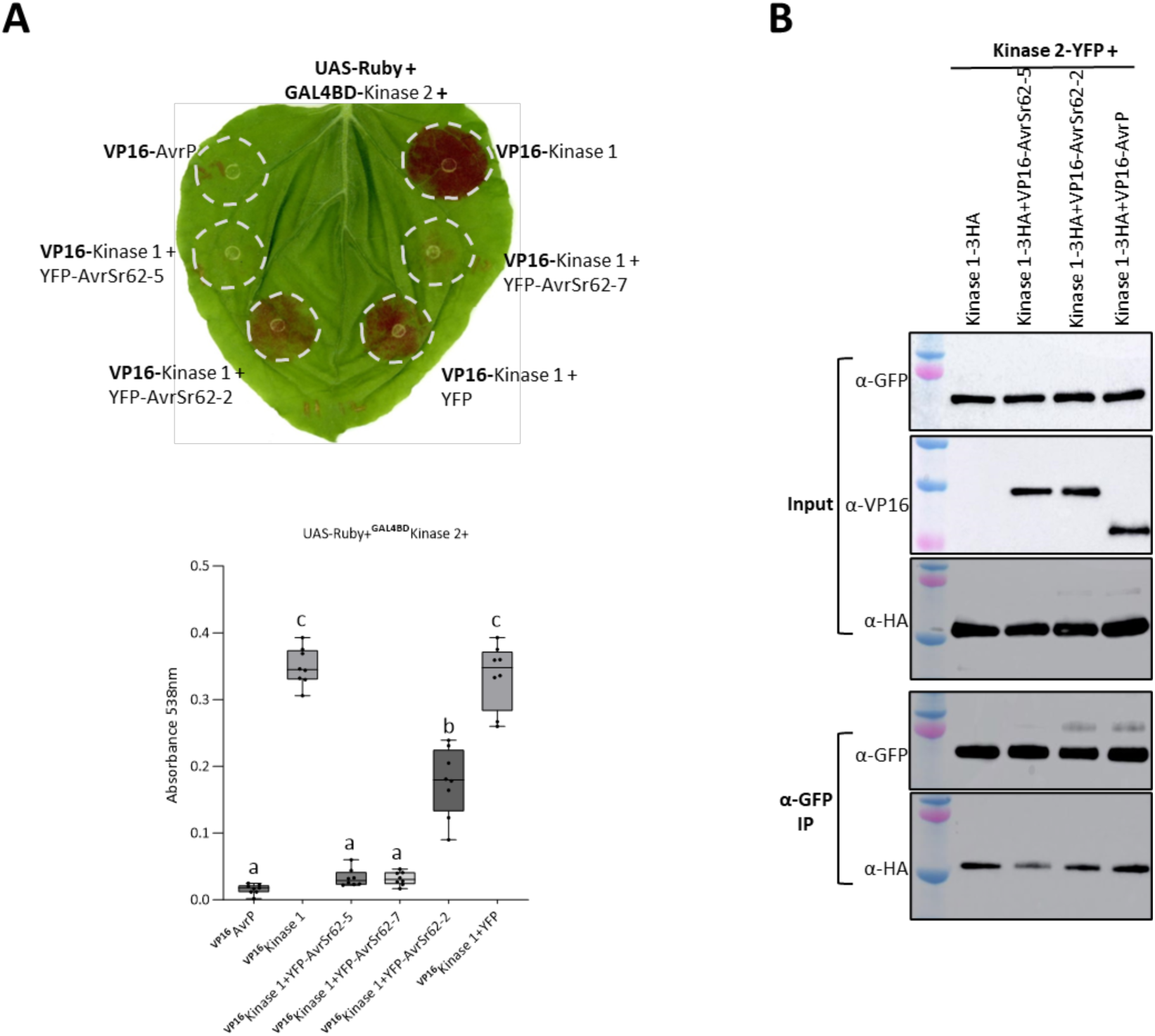
AvrSr62 effectors inhibit the interaction between Kinase 1 and Kinase 2 domains. (**A**) Betalain accumulation was observed in leaves expressing UAS-Ruby with Kinase 2 domain fused to BD and Kinase 1 domain fused to VP16 with the expression with different AvrSr62 effectors. Graph indicates betalain accumulation as box and whisker plots with at least five biological replicates (dots). Samples marked by identical letters in the plots do not differ significantly (P < 0.05; ANOVA Tukey test). (**B**) YFP-tagged Sr62^TK^ Kinase 2 domain was transiently co-expressed in *N. benthamiana* with HA-tagged Sr62^TK^ Kinase 1 and VP16-tagged AvrSr62-5, -7 or AvrP from flax rust as a negative control. The *Agrobacterium* concentration of Kinase 1-3HA was adjusted to OD_600_ = 0.1 to eliminate its high expression. Total protein was extracted and subjected to immunoprecipitation with an anti-GFP beads, followed by immunoblotting with anti-GFP, anti-HA and anti-VP16 antibodies.

**Fig. S16.**
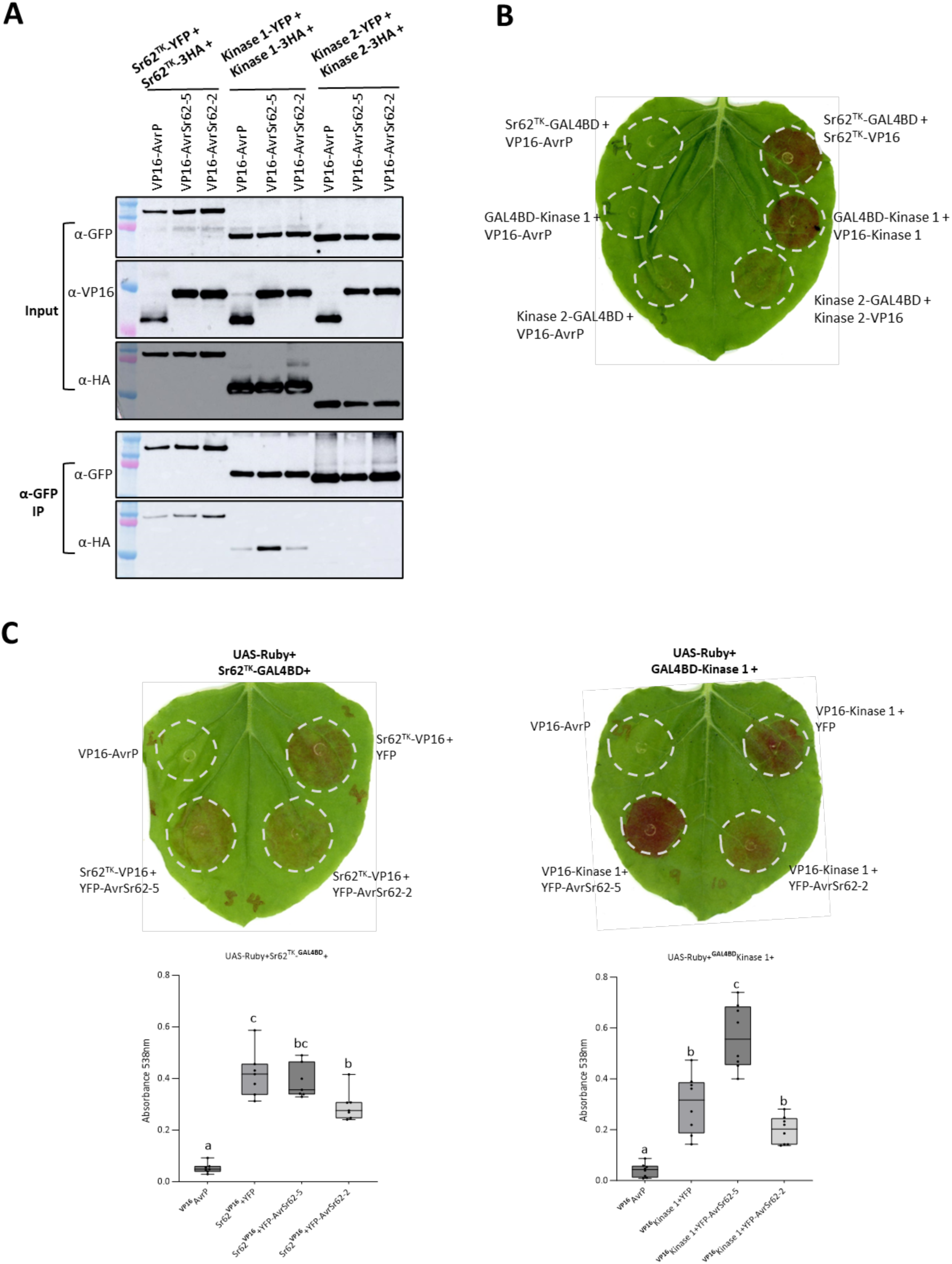
Self-association of Sr62^TK^ and Kinase 1 domains. (**A**). Sr62^TK^, Kinase 1 and Kinase 2 domains were expressed as both -YFP and -HA tagged versions along with VP16-tagged AvrSr62-6, -2 or AvrP from flax rust. Total protein was extracted and subjected to immunoprecipitation with anti-GFP beads, followed by immunoblotting with anti-GFP, anti-HA and anti-VP16 antibodies. The *Agrobacterium* concentration of Kinase 1-3HA was adjusted to OD_600_ = 0.1 to eliminate its high expression. (**B**). Betalain accumulation was observed in leaves expressing UAS-Ruby with Sr62^TK^, Kinase 1 and Kinase 2 domains fused to GAL4-BD and VP16-tagged AvrP (negative control), Sr62^TK^, Kinase 1 or Kinase 2 as indicated. (**C**) Betalain accumulation was observed in leaves expressing UAS-Ruby with Sr62^TK^ (left) or Kinase 1 (right) fused to GAL4-BD plus VP16-tagged AvrP (negative control) or Sr62^TK^ (left) or Kinase 1 (right) with or without YFP-fused AvrSr62-2, AvrSr62-5 or YFP as indicated. Graph indicates betalain accumulation as box and whisker plots with at least five biological replicates (dots). Samples marked by identical letters in the plots do not differ significantly (P < 0.05; ANOVA Tukey test).

**Fig. S17.**
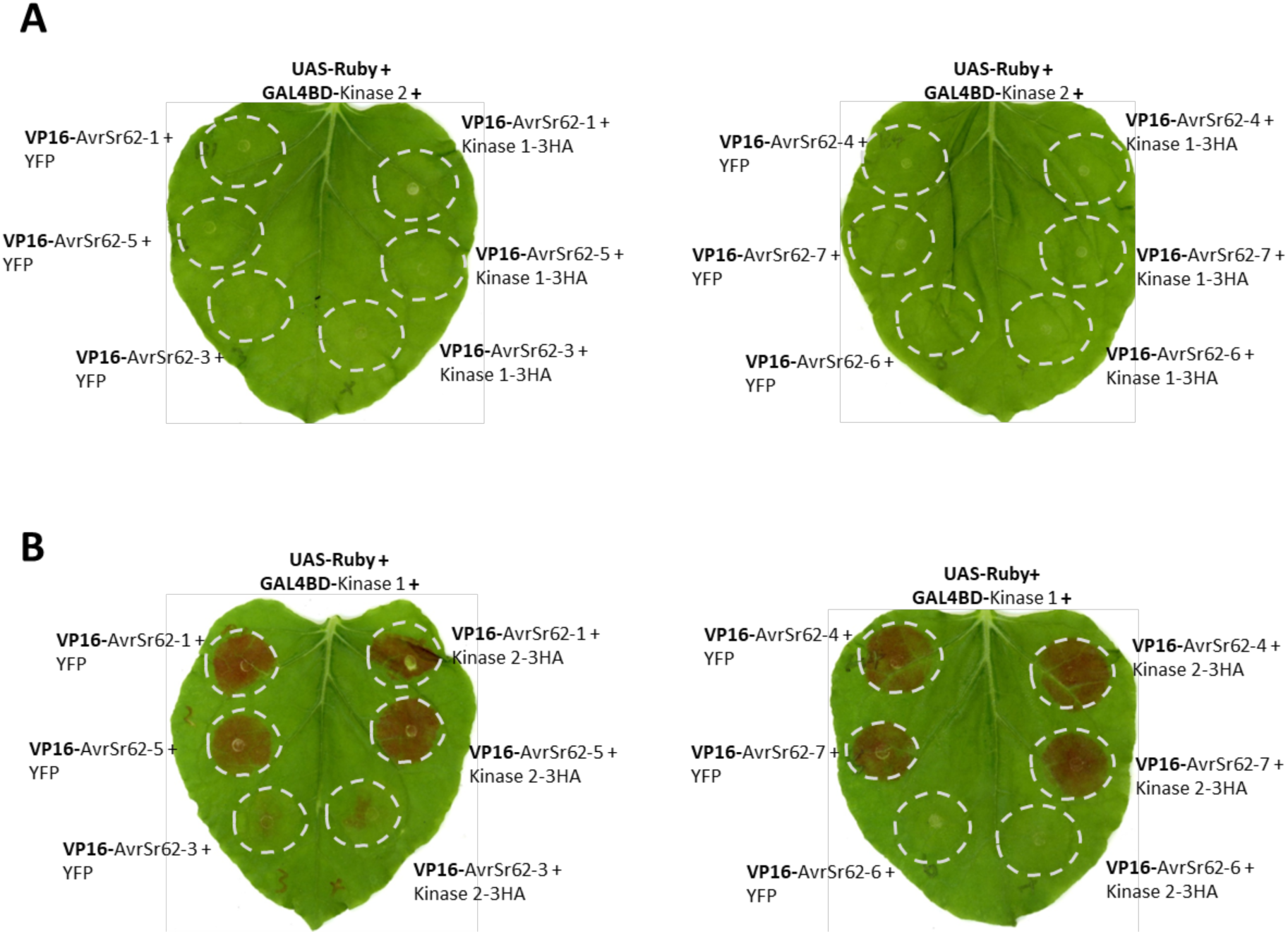
Comparison of Kinase 1, Kinase 2 and AvrSr62 interactions. (**A**) Betalain accumulation was observed in leaves expressing Sr62^TK^ Kinase 2 domain fused to GAL4-BD withVP16-fused AvrSr62 variants with or without Kinase 1 fused to HA. No Ruby expression was detected indicating that Kinase 1 does not mediate an indirect interaction between kinase 2 and AvrSr62 in a three-way complex. This suggests that the Kinase 1 interactions with Kinase 2 and with AvrSr62 are mutually exclusive. (**B**) Betalain accumulation was observed in leaves expressing Sr62^TK^ Kinase 1 domain fused to GAL4-BD with VP16-fused AvrSr62 variants with or without Kinase 2 fused to HA. The presence of Kinase 2 did not affect the interaction observed between Kinase 1 and the AvrSr62 proteins. This is in contrast to the results in Fig 3D and S15, where AvrSr62 expression inhibits the interaction between Kinase 1 and Kinase 2 domains, suggesting that the Kinase 1-AvrSr62 interaction is favored over the Kinase 1-Kinase 2 interaction.

**Fig. S18.**
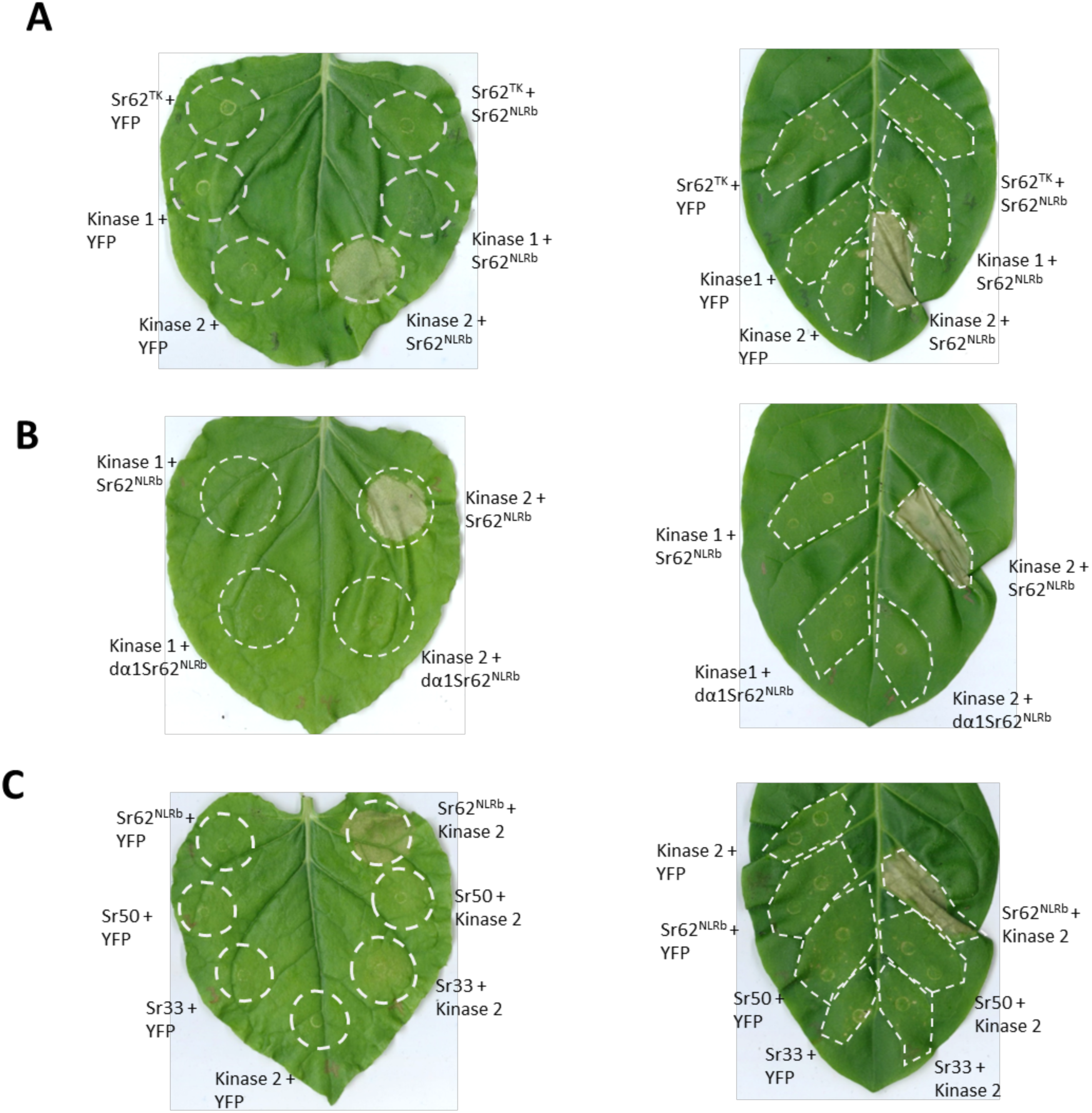
Sr62^TK^ Kinase 2 domain is sufficient to activate Sr62^NLR^. (**A**) Sr62^TK^, or its Kinase 1 or Kinase 2 domains alone fused to HA (C-terminal) were co-expressed with YFP or HA (C-terminal) tagged Sr62^NLRb^ in *N. benthamiana* (left) and *N. tabacum* (right). Leaves were visualized after 3 days. (**B**) Sr62^TK^ Kinase 1 or Kinase 2 domains fused to HA (C-terminal) were co-expressed with HA tagged Sr62^NLRb^ or a mutant version with the N-terminal alpha-1 helix deleted (dα1Sr62^NLRb^). Leaves were visualized after 3 days. (**C**) HA-tagged Sr62^NLRb^, and YFP tagged Sr50 or Sr33 were co-expressed with YFP or Sr62^TK^ Kinase 2 domain (HA-tagged). Kinase 2 did not trigger cell death in combination with Sr33 or Sr50, indicating that this response is specific to Sr62^NLR^.

**Fig. S19.**
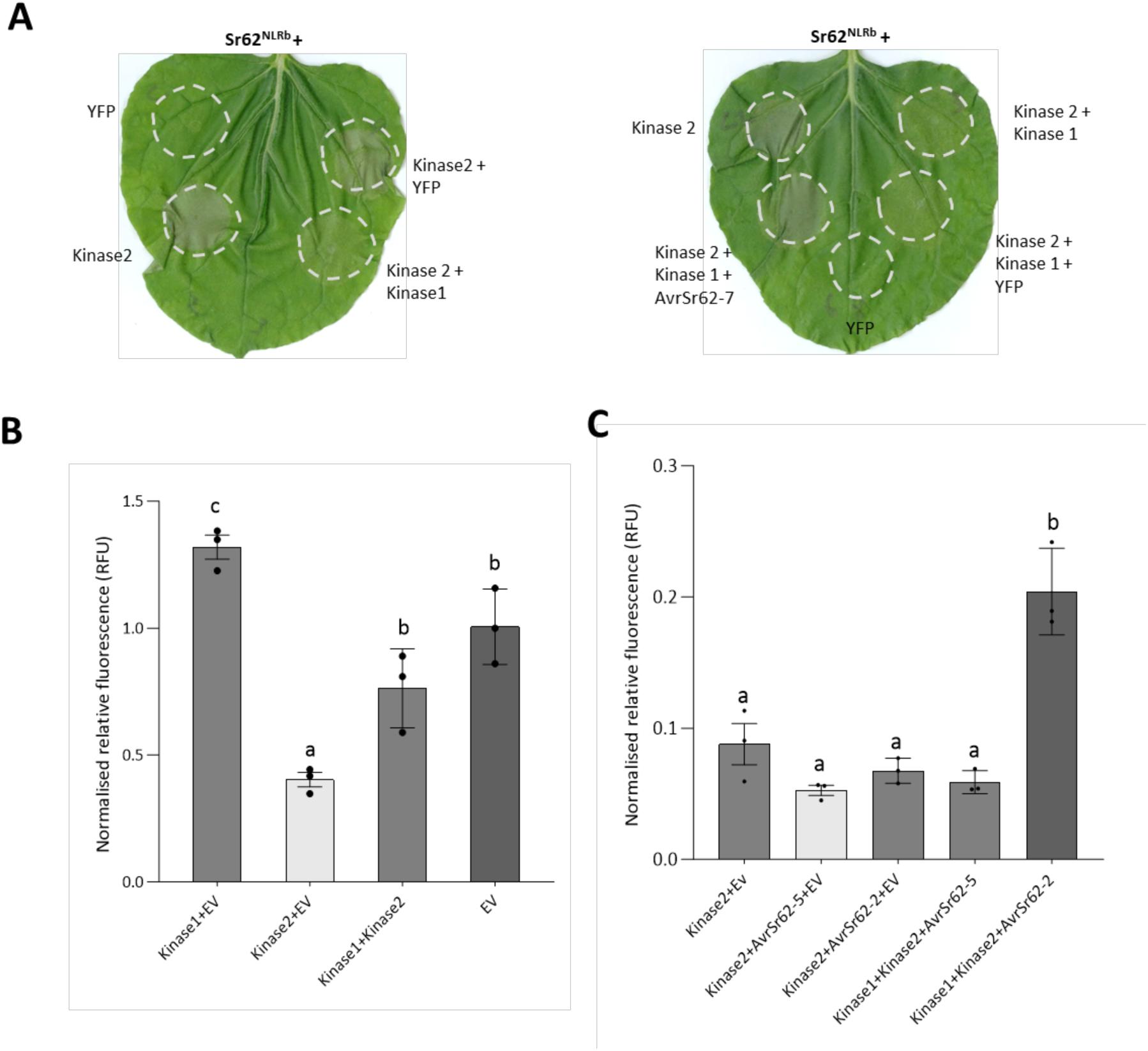
Kinase 1 domain inhibits cell death caused by Kinase 2 domain and Sr62^NLRb^, whereas co-expression of AvrSr62 releases the inhibition. (**A**) Sr62^NLRb^ was co-expressed in *N. benthamiana* with Kinase 1, Kinase 2 domains, YFP and/or AvrSr62-7 in the indicated combinations. Sr62^NLRb^ and Sr62^TK^ Kinase1 and Kinase 2 domains are fused to a C terminal 3xHA tag. AvrSr62-7 has a YFP tag at the N terminus. (**B**) and (**C**) Sr62^NLRb^, Sr62^TK^, Kinase 1 and Kinase 2 domains, and AvrSr62-2 or -5 were co-transformed into wheat protoplasts in the indicated combinations along with a YFP reporter construct. YFP signals were measured at 16 hpt. Results represent the means of three biological replicates (dots) with error bars indicating the standard error. Samples marked by identical letters in the plots do not differ significantly (P < 0.05; ANOVA Tukey test). The amount of the Kinase 1 construct in the transformation was 3000 fmol, while 375 fmol of the Kinase 2 construct was used.

**Fig. S20.**
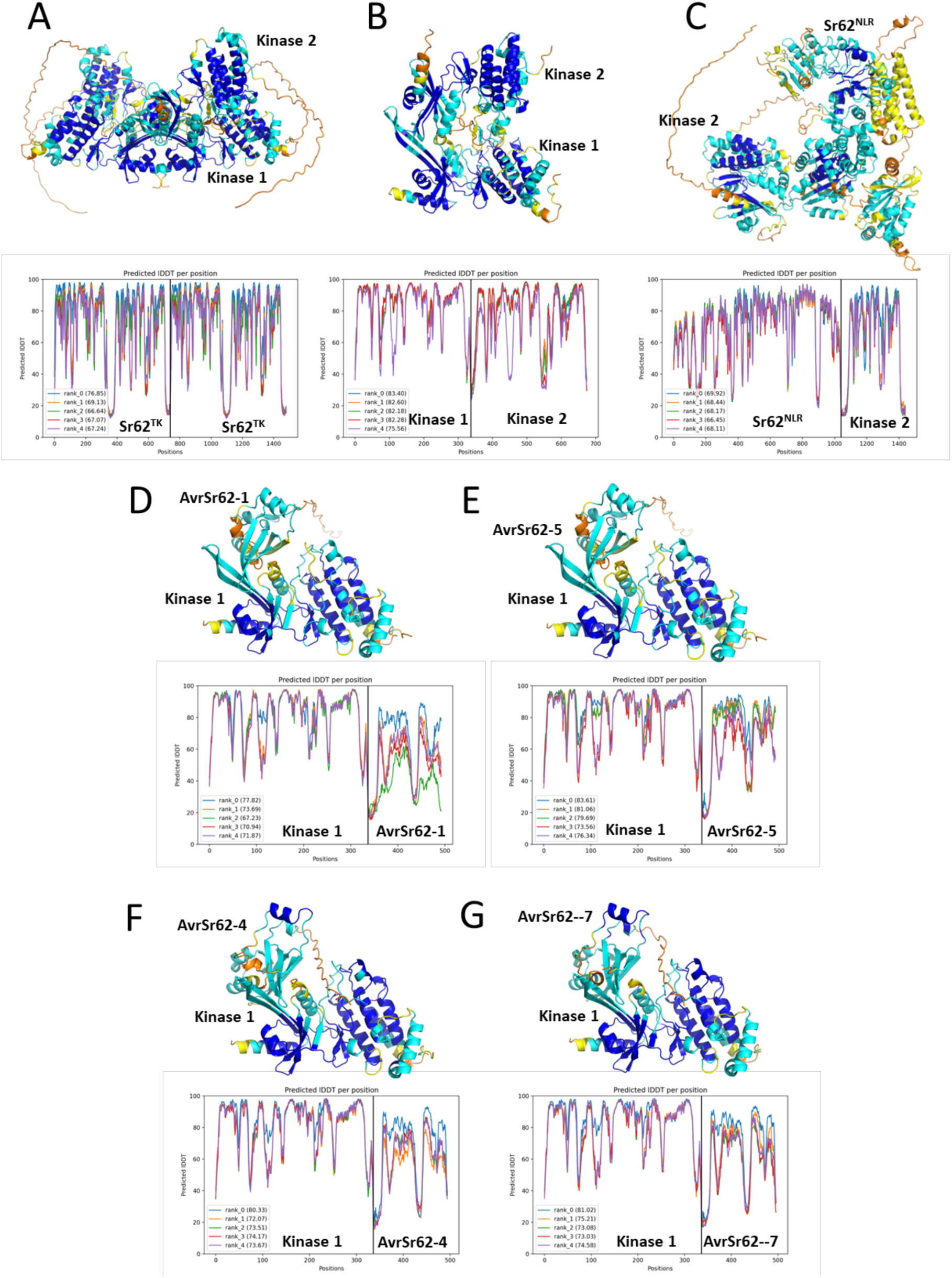
AlphaFold predicted local Distance Difference Test pLDDT scores per residue. Upper panels: molecular models colored by pLDDT, with dark blue: plDDT > 90 (very high confidence); light blue: 90 > pLDDT > 70 (high); yellow: 70 > pLDDT > 50 (low); orange pLDDT < 50 (very low). Lower panels: Plots of pLDDT vs residue number for 5 independent prediction runs per complex. (**A**) Sr62^TK^ dimer. (**B**) Sr62^TK^ Kinase 1 and Kinase 2 (taken as independent chains). (**C**) Sr62^NLR^ in complex with Sr62^TK^ Kinase 2 (complex as in fig. S20C); (**D** to **G**) Effectors bound to Sr62^TK^ Kinase 1.

**Fig. S21.**
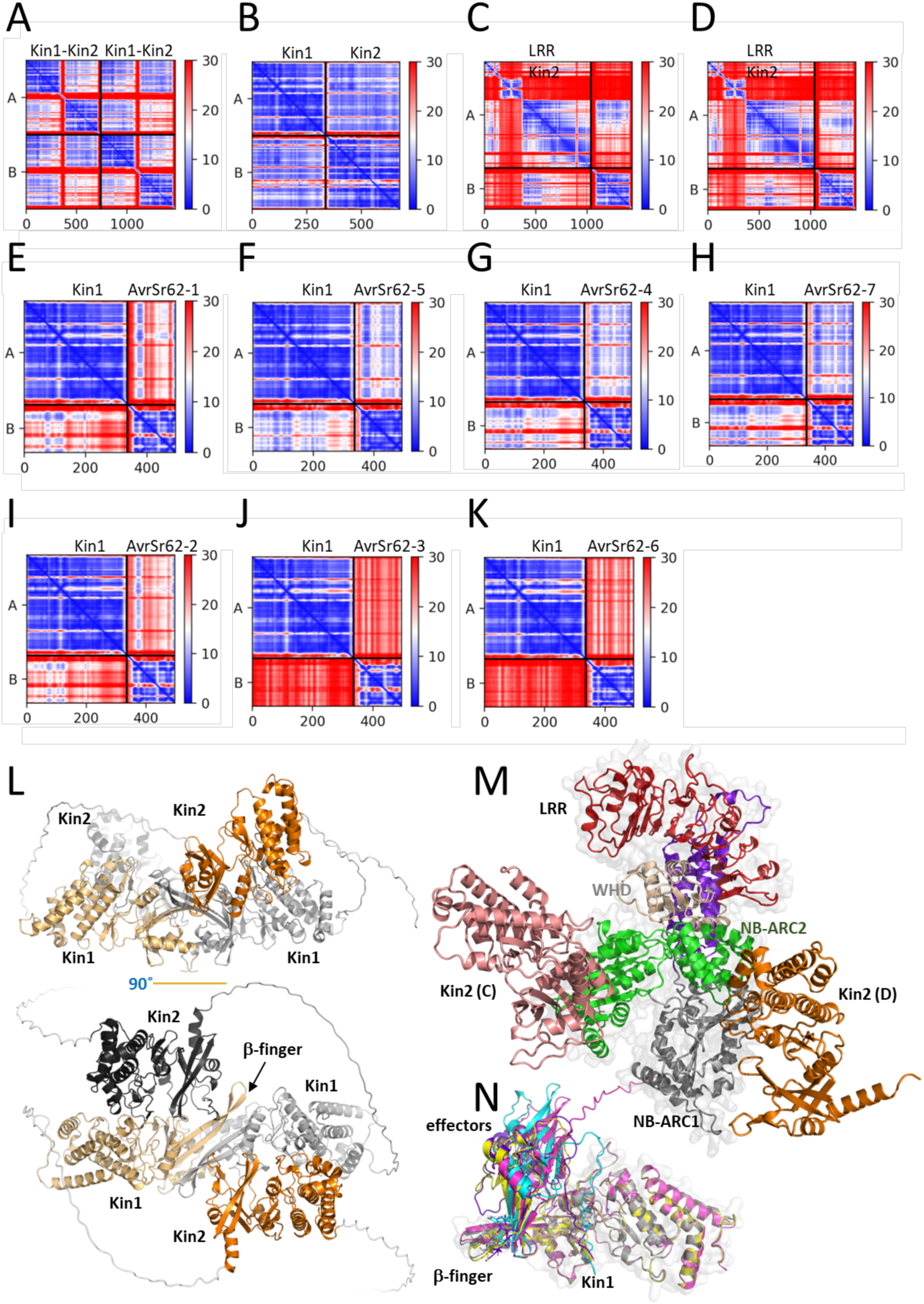
Protein interaction predictions and AlphaFold models. (**A-K**) Predicted aligned error (PAE). Coloration is color-ramped blue to red from 0 to 30 Å. The sequences corresponding to the fields are shown. (**A**) Sr62^TK^ dimer. (**B**) Sr62^TK^ Kinase 1 vs Kinase 2. (**C**) Sr62^NLR^ vs Sr62^TK^ Kinase 2 binding to the NB of the NLR NB-ARC2. (**D**) Sr62^NLR^ vs Sr62^TK^ Kinase 2 binding to the ARC of the Sr62^NLR^ NB-ARC2. (**E**) Sr62^TK^ Kinase 1 vs AvrSr62-1. (**F**) Sr62^TK^ Kinase 1 vs AvrSr62-5. (**G**) Sr62^TK^ Kinase 1 vs AvrSr62-4. (**H**) Sr62^TK^ Kinase 1 vs AvrSr62-7. (**I**) Sr62^TK^ Kinase 1 vs AvrSr62-2. (**J**) Sr62^TK^ Kinase 1 vs AvrSr62-3. (**K**) Sr62^TK^ Kinase 1 vs 1059. (**L-N**) AlphaFold models. (**L**) the Sr62^TK^ self-association. Kinase 1 and Kinase 2 of both chains are coloured bright and pale orange (chain 1) or light and dark gray (chain 2), respectively. Side and top views are shown. (**M**) Association between Sr62^NLR^ (domains are color-coded) and Sr62^TK^ Kinase 2 (orange). Shown are the two Kinase 2 positions predicted at the NB domain of NB-ARC2 [labelled Kin2 (C), corresponding to the PAE shown in C] and at the ARC domain of NB-ARC2 [labelled Kin2 (D), corresponding to the PAE shown in D]. (**N**) Superimposition of effectors bound to Sr62^TK^ Kinase 1. Shown are models for AvrSr62-5 (cyan, Kinase 2 in grey), AvrSr62-1 (magenta), AvrSr62-4 (yellow), AvrSr62-7 (purple). Kin1 and Kin2 denote Kinase 1 and Kinase 2.

**Fig. S22.**
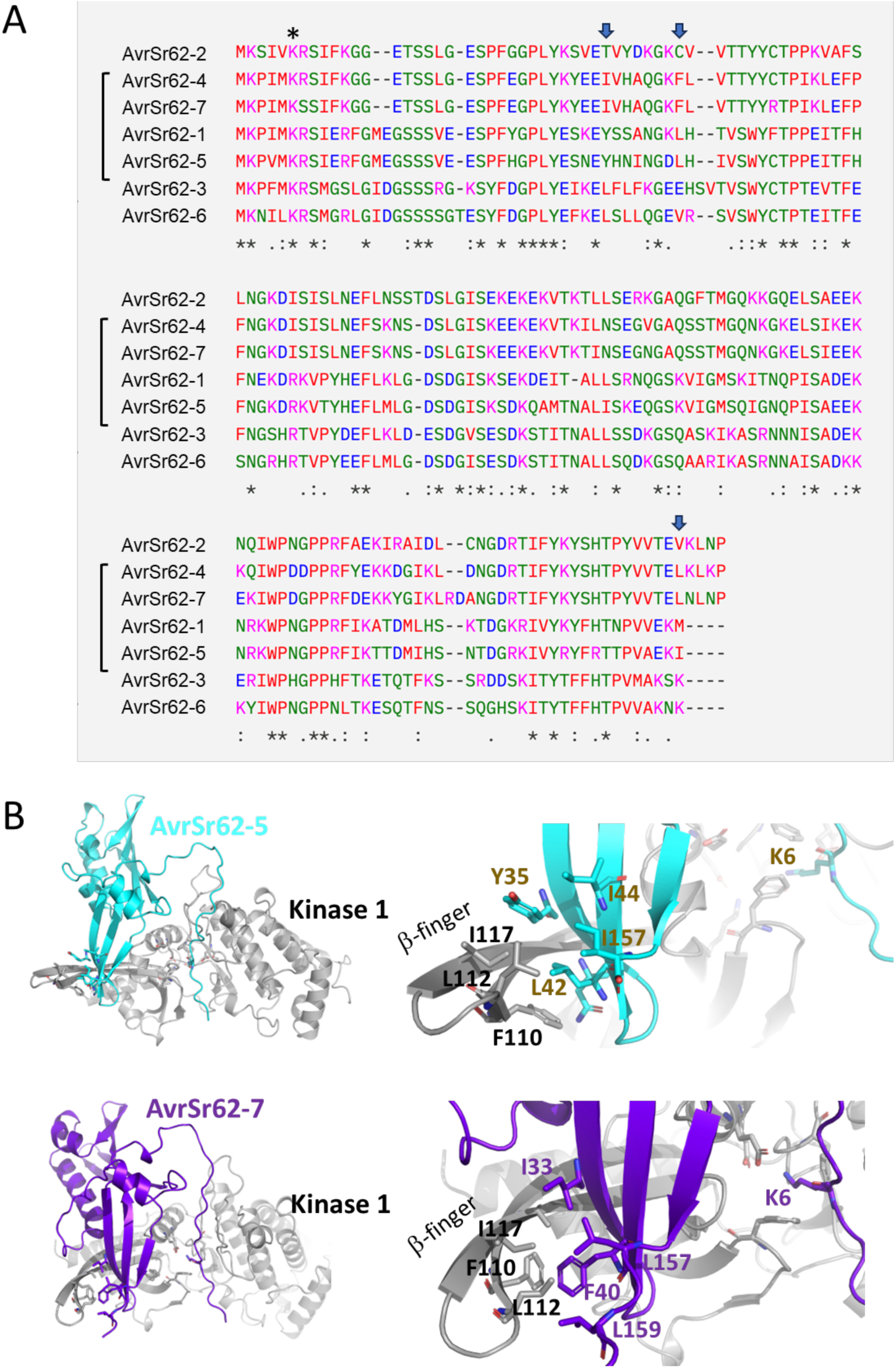
Interactions between Sr62^TK^ Kinase 1 and effectors. (**A**) Effector sequence alignment. Effectors experimentally shown to bind Sr62^TK^ are indicated by a bracket. Key positions of the effector β-sheet that correlate with Sr62^TK^ Kinase 1 binding are indicated by an arrow. The asterisk indicates the Lysine 6 that reaches into the Sr62^TK^ Kinase 1 active site. (**B**) Molecular AlphaFold models corresponding to the association between Sr62^TK^ Kinase 1 (gray) and two effectors (472, cyan; 490, purple). Right panel shows the zoom into the primary binding site.

**Fig. S23.**
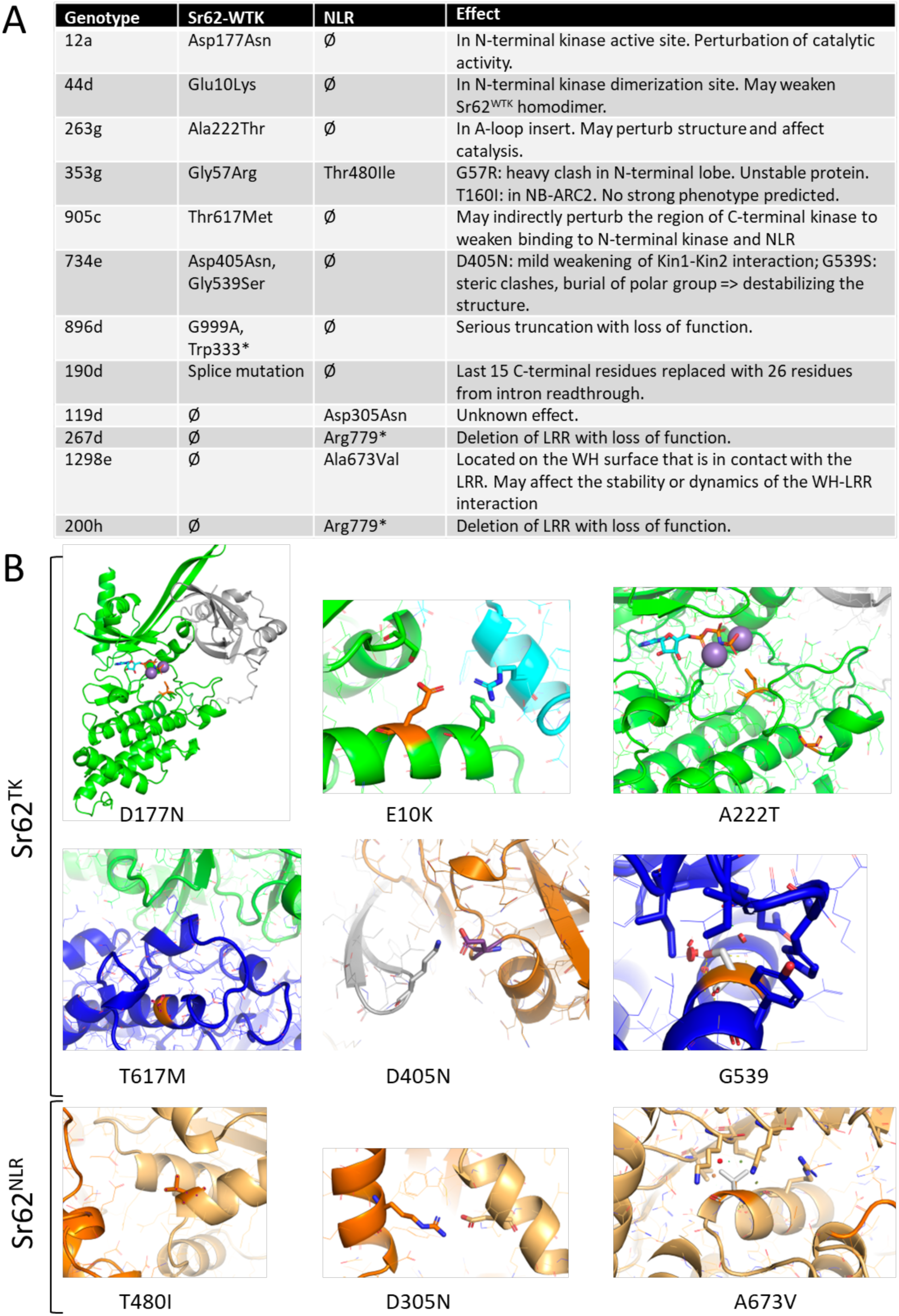
Mapping of the Sr62^TK^ and Sr62^NLR^ EMS-induced mutations onto the 3D structures. (A) Table showing the mutations and the effect predicted from the AlphaFold structural models. (B) Molecular visualization of the mutations in their 3D environment. The affected residues are shown in stick representation, with their carbon atoms colored in orange.

**Fig. S24.**
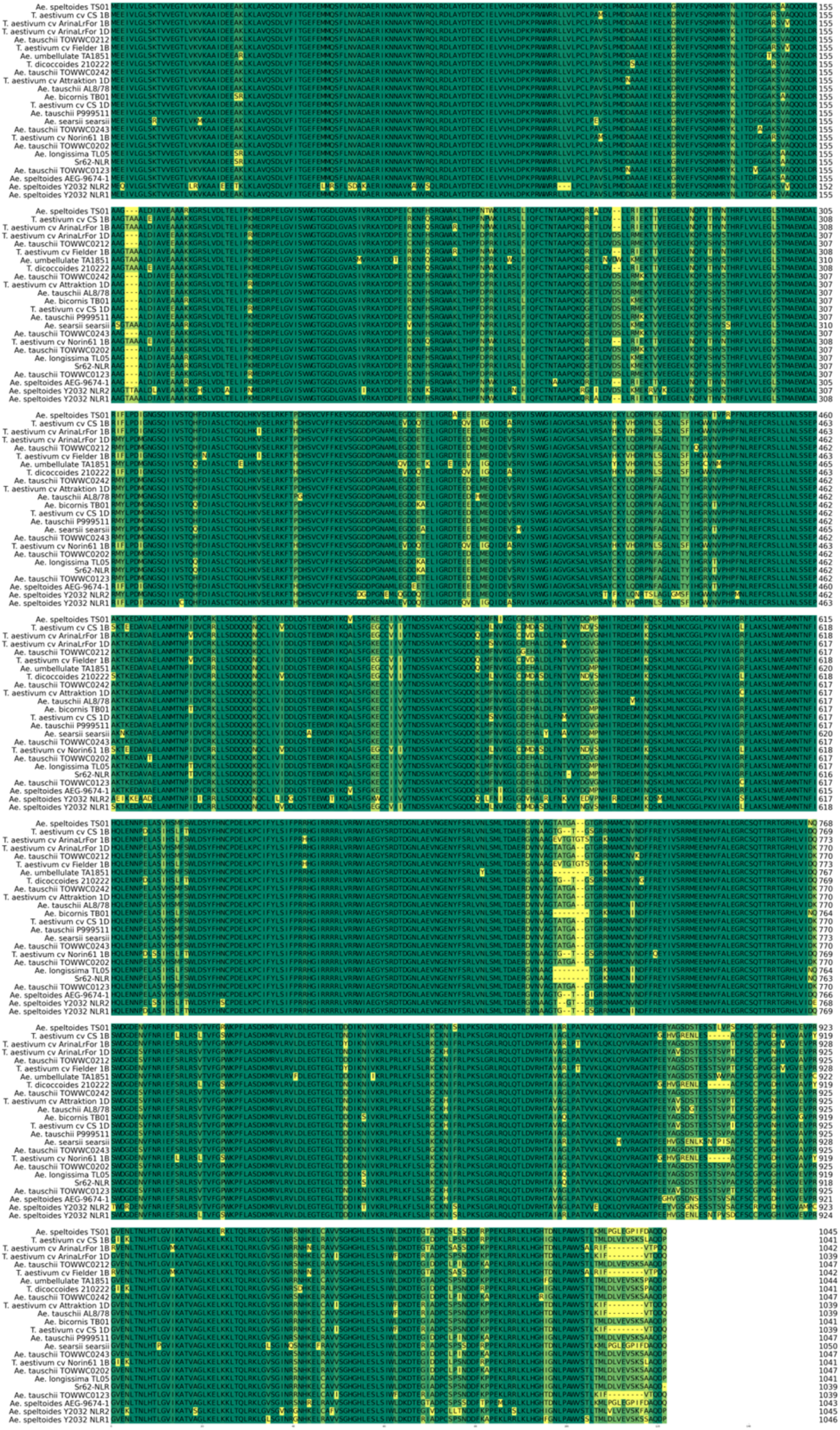
Alignment of Triticeae *Sr62^NLR^* homologues (non-redundant sequences).

**Fig. S25.**
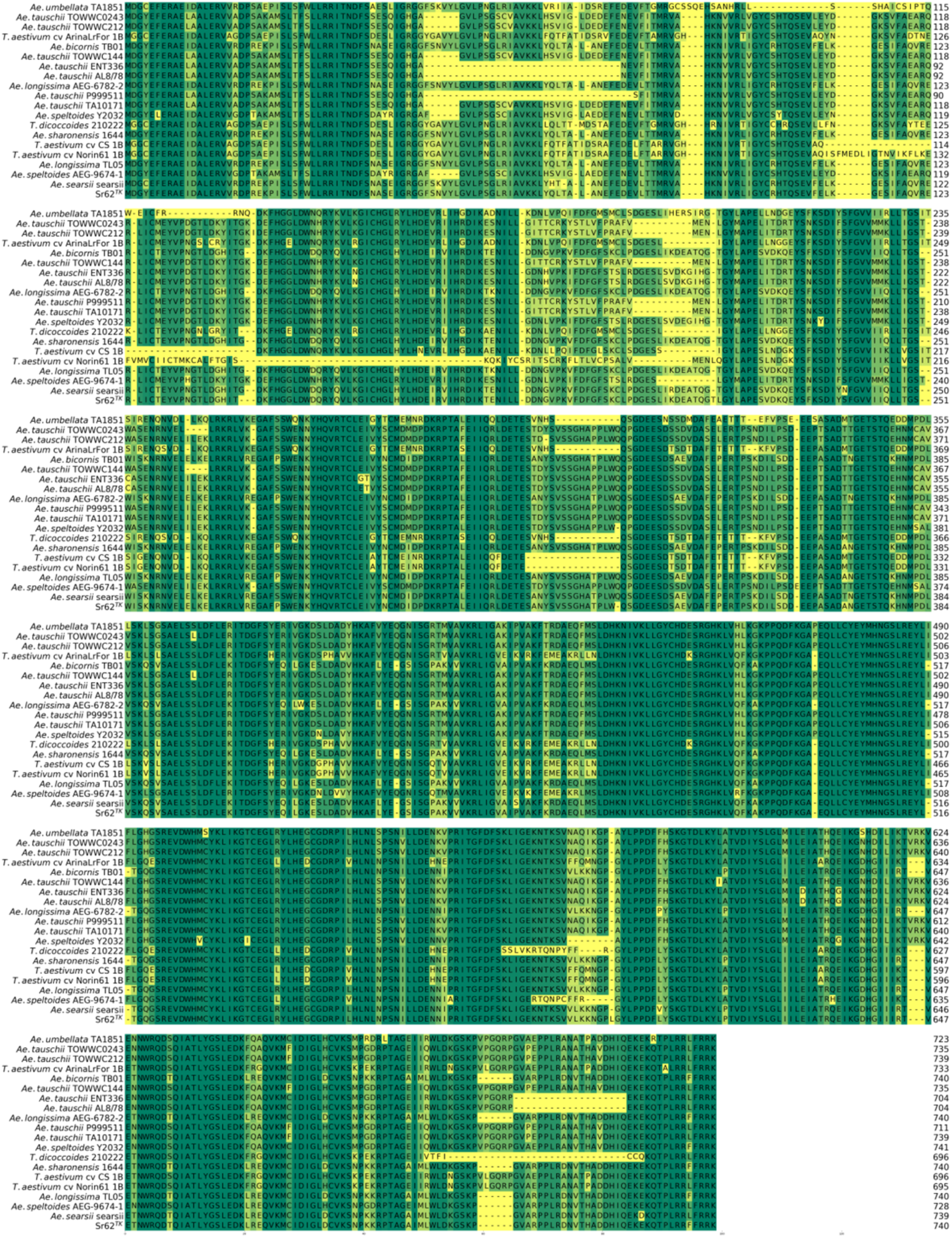
Alignment of Triticeae Sr62^TK^ homologues (non-redundant sequences).

## Supplementary Tables S1 to S12 are provided in a separate excel file as Data S2

**Table S1.** Wheat-*Aegilops sharonensis* Zahir-1644 PacBio sequencing and assembly metrics.

**Table S2.** Wheat-*Aegilops sharonensis* 1644 wildtype and mutant Illumina RNA-Seq and whole genome shotgun (WGS) sequencing metrics.

**Table S3.** Seedling infection types (ITs) of wheat-*Aegilops sharonensis* 1644 wildtype and mutants.

**Table S4.** RNA-Seq, WGS, and exome-capture sequence data analysis of EMS-derived stem rust susceptible mutants.

**Table S5.** Wheat-*Aegilops sharonensis* Zahir-1644 PacBio IsoSeq HiFi read metrics.

**Table S6.** Segregation of stem rust susceptible plants in BC_1_F_2_ populations derived from backcrossing EMS-derived susceptible mutants to the Zahir-1644 parental non-mutated line.

**Table S7.** Genotyping of *Sr62^NLR^* mutations in individual BC_1_F_2_ plants from populations 1298e (no. 2) and 267d by PCR and Sanger sequencing.

**Table S8.** Mutation overlap in gene loci from BC_1_F_2_ bulk susceptible exome capture data.

**Table S9**. Interaction scores for endogenous associations involving Sr62^TK^.

**Table S10.** Interaction scores between effectors and Sr62^TK^ Kinase 1.

**Table S11.** Orthologues of *Sr62^TK^* and *Sr62^NLR^* identified in *Triticum* and *Aegilops* genomes.

**Table S12.** Plasmids used in this study.

**Table S13.** Primers used in this study.

**Data S1. (separate file)**

PDF containining IGV genome browser screenhots of RNA, WGS, and exome-capture sequences of *Sr62^TK^*and *Sr62^NLR^* mutants aligned to the Zahir-1644 genome assembly.

**Data S2. (separate file)**

Excel file contains supplementary tables S1 to S12.

