## Supplementary material for "A wheat tandem kinase sensor activates an NLR helper to trigger immunity": Data S1.pptx

### Slide 1
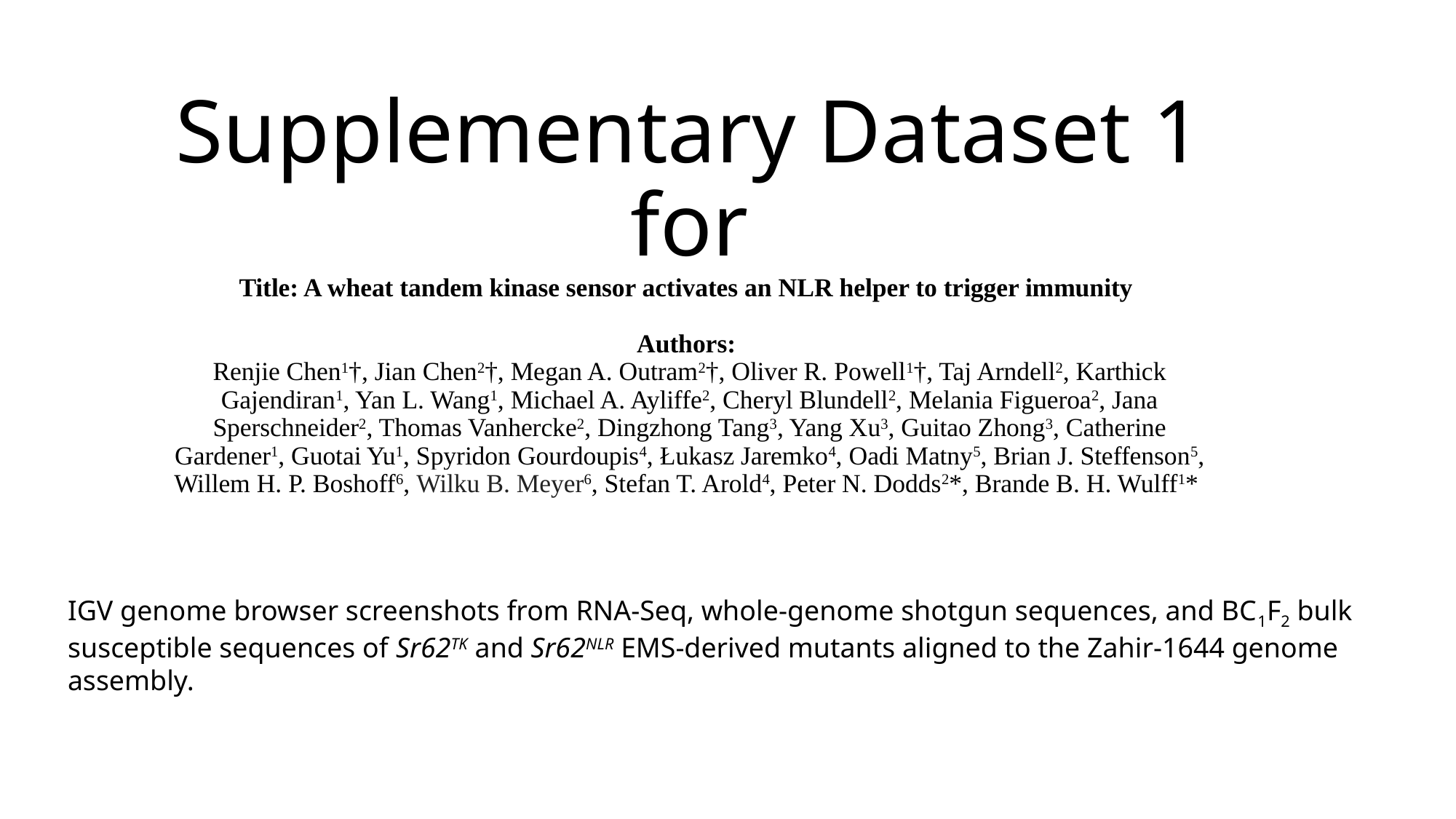

#### Supplementary Dataset 1 for Title: A wheat tandem kinase sensor activates an NLR helper to trigger immunity Authors: Renjie Chen^1^†, Jian Chen^2^†, Megan A. Outram^2^†, Oliver R. Powell^1^†, Taj Arndell^2^, Karthick Gajendiran^1^, Yan L. Wang^1^, Michael A. Ayliffe^2^, Cheryl Blundell^2^, Melania Figueroa^2^, Jana Sperschneider^2^, Thomas Vanhercke^2^, Dingzhong Tang^3^, Yang Xu^3^, Guitao Zhong^3^, Catherine Gardener^1^, Guotai Yu^1^, Spyridon Gourdoupis^4^, Łukasz Jaremko^4^, Oadi Matny^5^, Brian J. Steffenson^5^, Willem H. P. Boshoff^6^, Wilku B. Meyer^6^, Stefan T. Arold^4^, Peter N. Dodds^2^*, Brande B. H. Wulff^1^*
IGV genome browser screenshots from RNA-Seq, whole-genome shotgun sequences, and BC_1_F_2_ bulk susceptible sequences of Sr62^TK^ and Sr62^NLR^ EMS-derived mutants aligned to the Zahir-1644 genome assembly.

### Slide 2
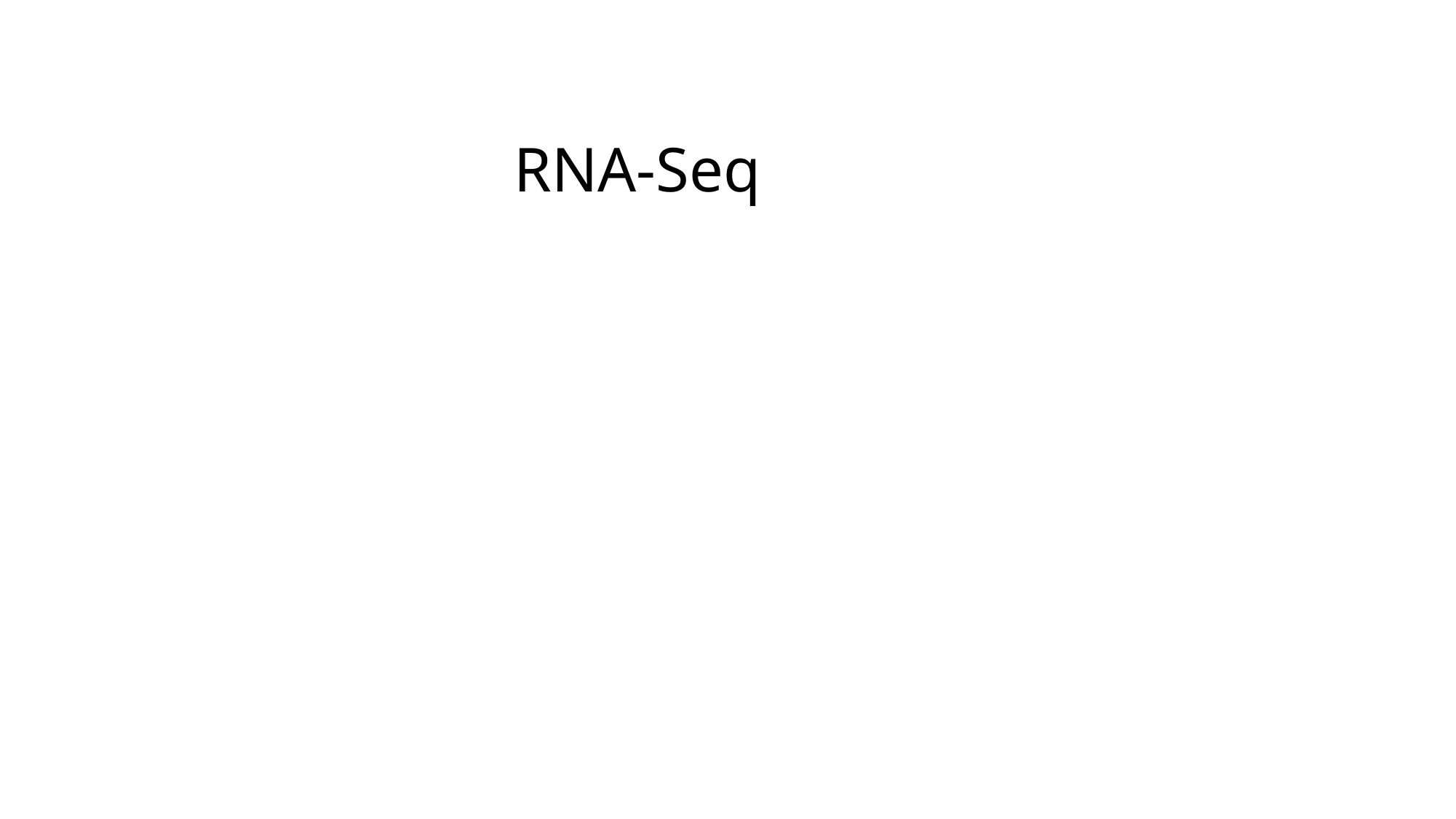

RNA-Seq

### Slide 3
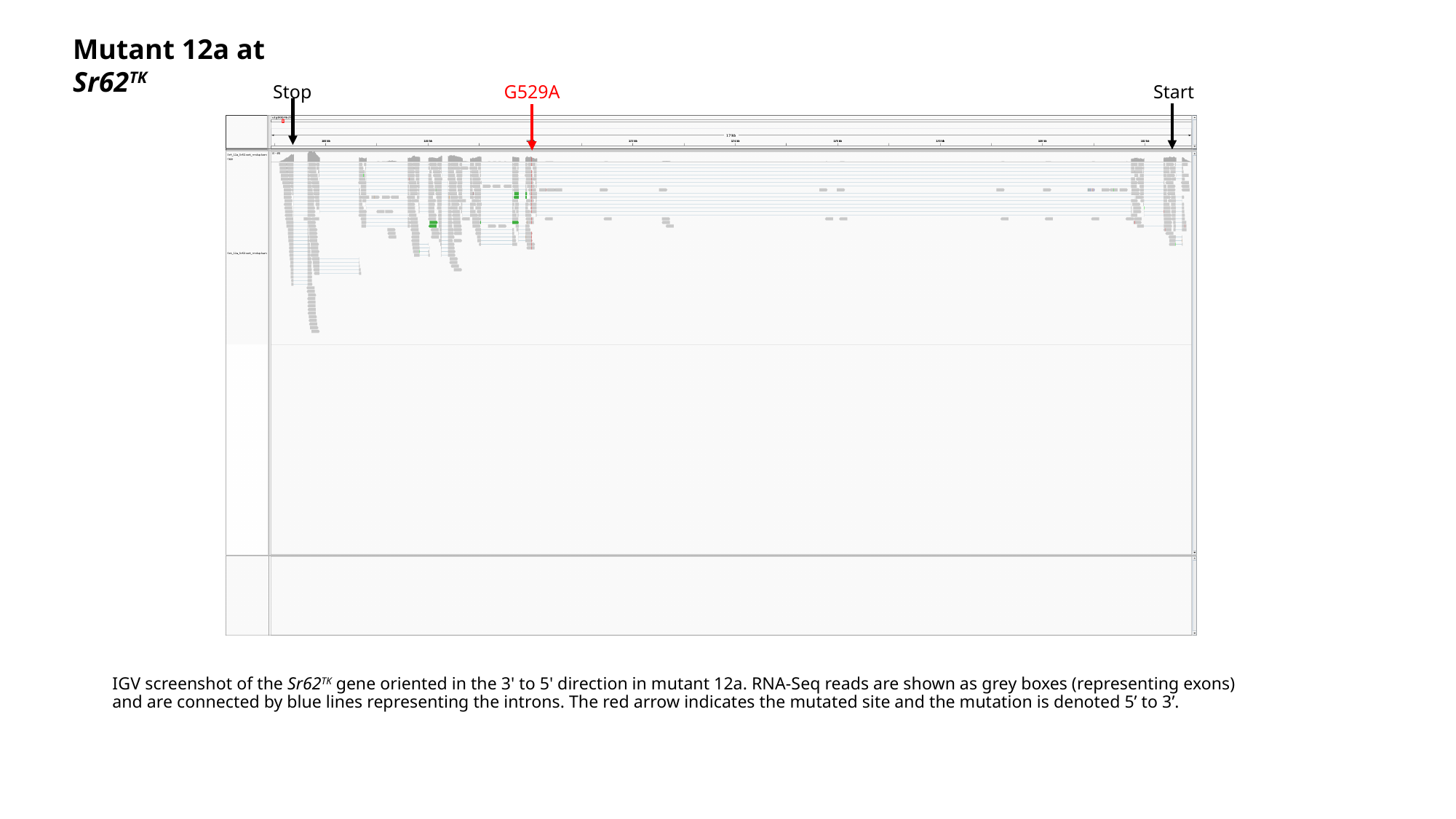

Mutant 12a at Sr62^TK^
Stop
G529A
Start
#### IGV screenshot of the Sr62^TK^ gene oriented in the 3' to 5' direction in mutant 12a. RNA-Seq reads are shown as grey boxes (representing exons) and are connected by blue lines representing the introns. The red arrow indicates the mutated site and the mutation is denoted 5’ to 3’.

### Slide 4
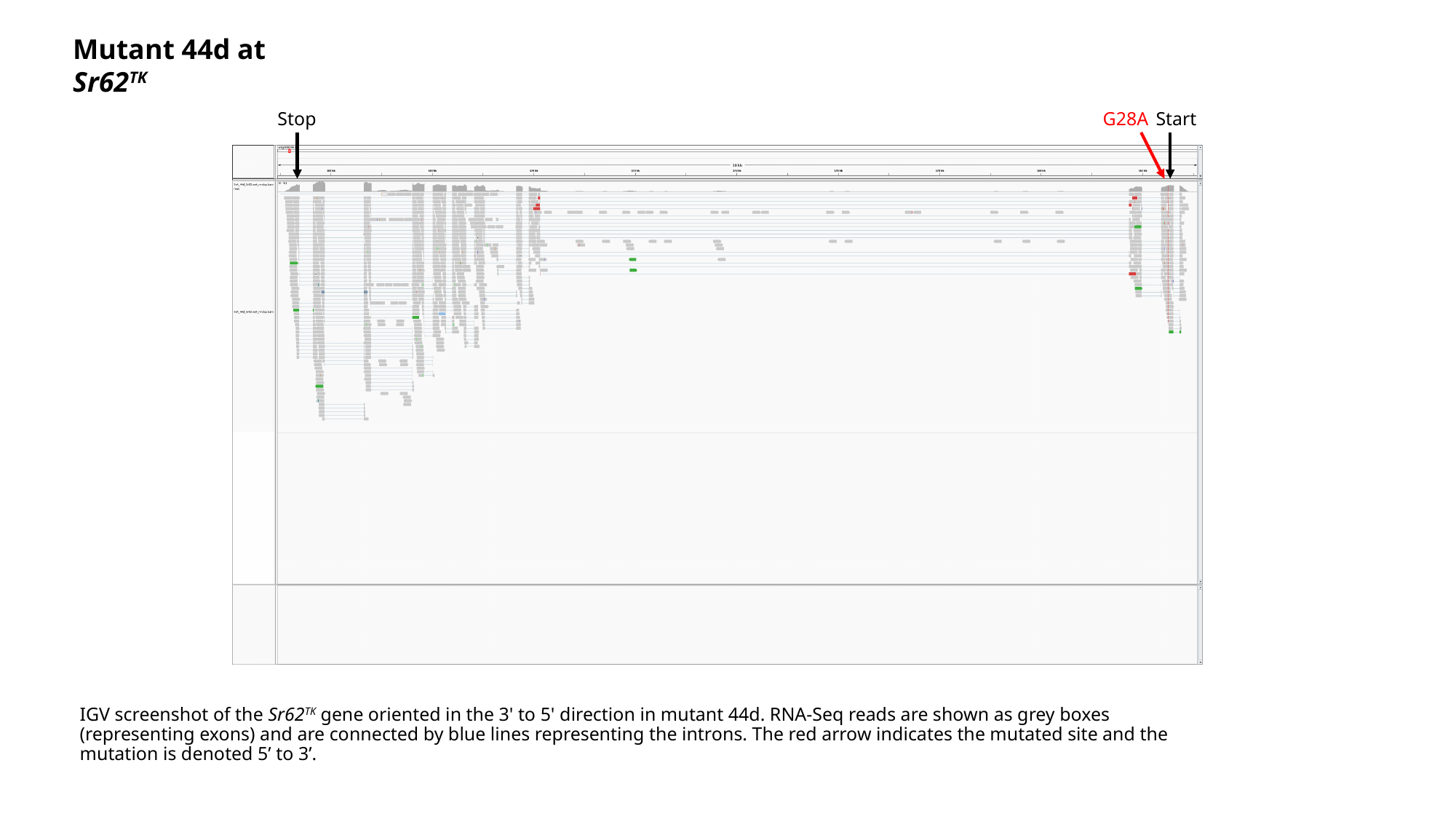

Mutant 44d at Sr62^TK^
Stop
G28A
Start
#### IGV screenshot of the Sr62^TK^ gene oriented in the 3' to 5' direction in mutant 44d. RNA-Seq reads are shown as grey boxes (representing exons) and are connected by blue lines representing the introns. The red arrow indicates the mutated site and the mutation is denoted 5’ to 3’.

### Slide 5
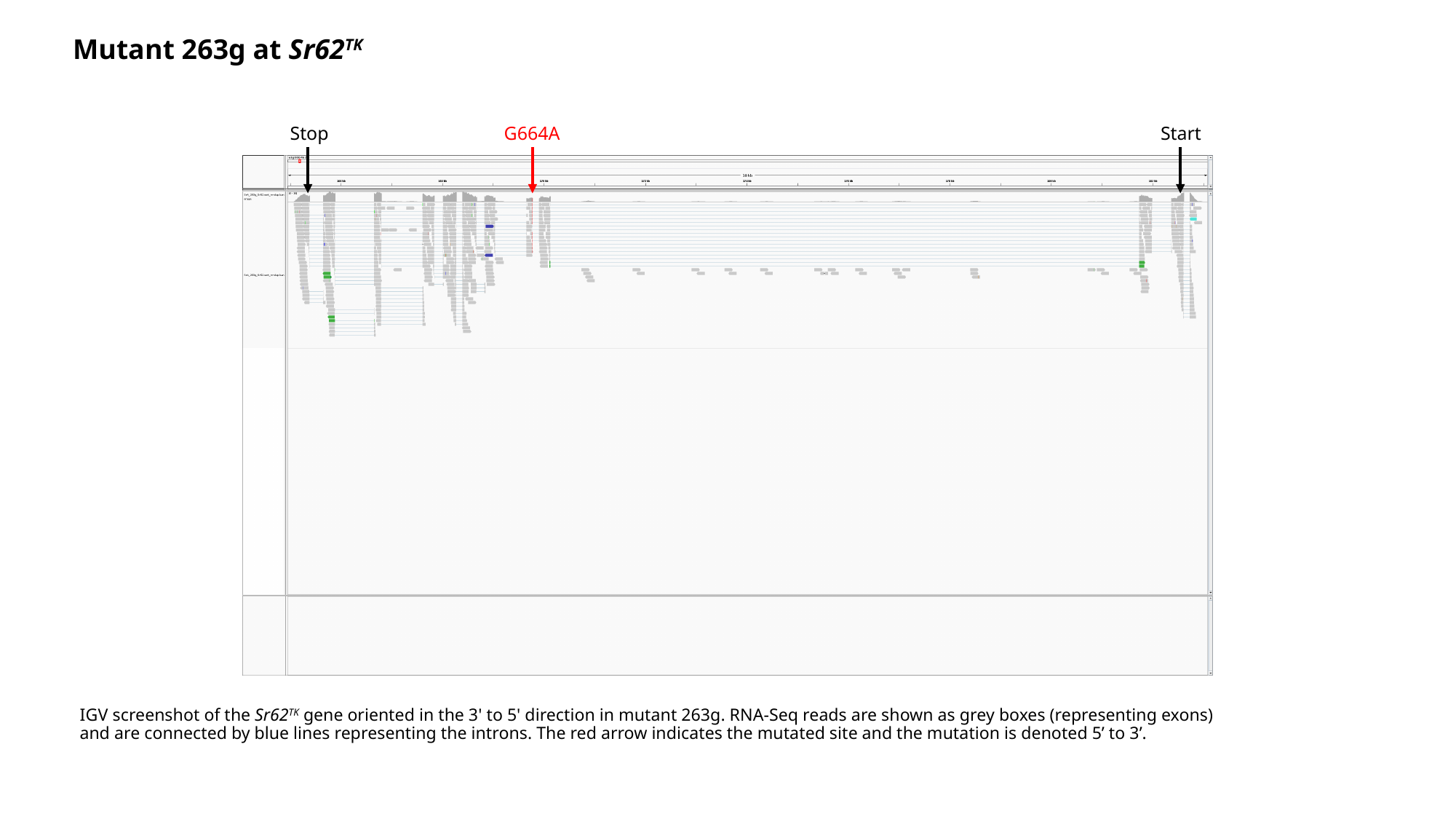

Mutant 263g at Sr62^TK^
Stop
G664A
Start
#### IGV screenshot of the Sr62^TK^ gene oriented in the 3' to 5' direction in mutant 263g. RNA-Seq reads are shown as grey boxes (representing exons) and are connected by blue lines representing the introns. The red arrow indicates the mutated site and the mutation is denoted 5’ to 3’.

### Slide 6
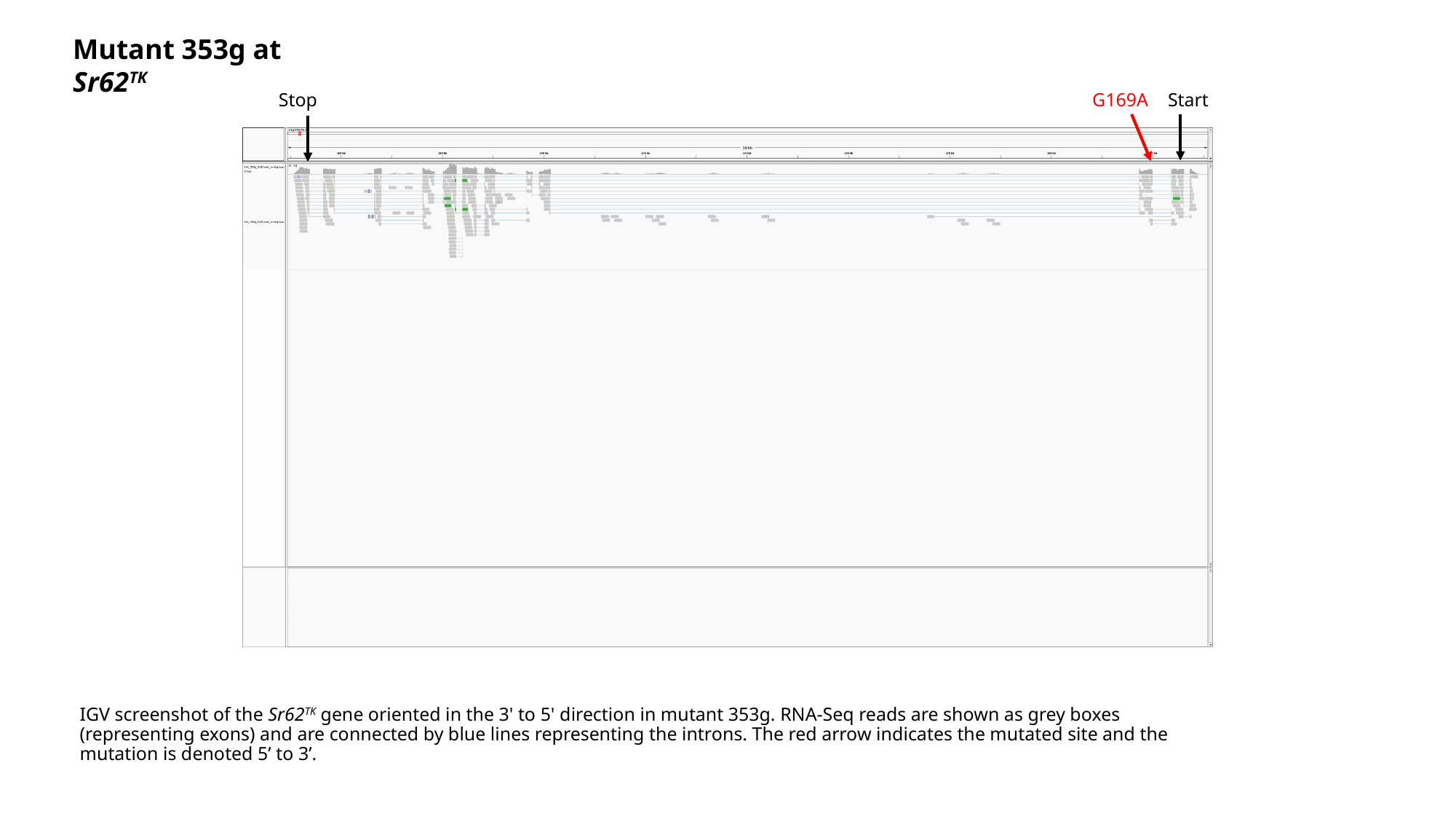

Mutant 353g at Sr62^TK^
Stop
G169A
Start
#### IGV screenshot of the Sr62^TK^ gene oriented in the 3' to 5' direction in mutant 353g. RNA-Seq reads are shown as grey boxes (representing exons) and are connected by blue lines representing the introns. The red arrow indicates the mutated site and the mutation is denoted 5’ to 3’.

### Slide 7
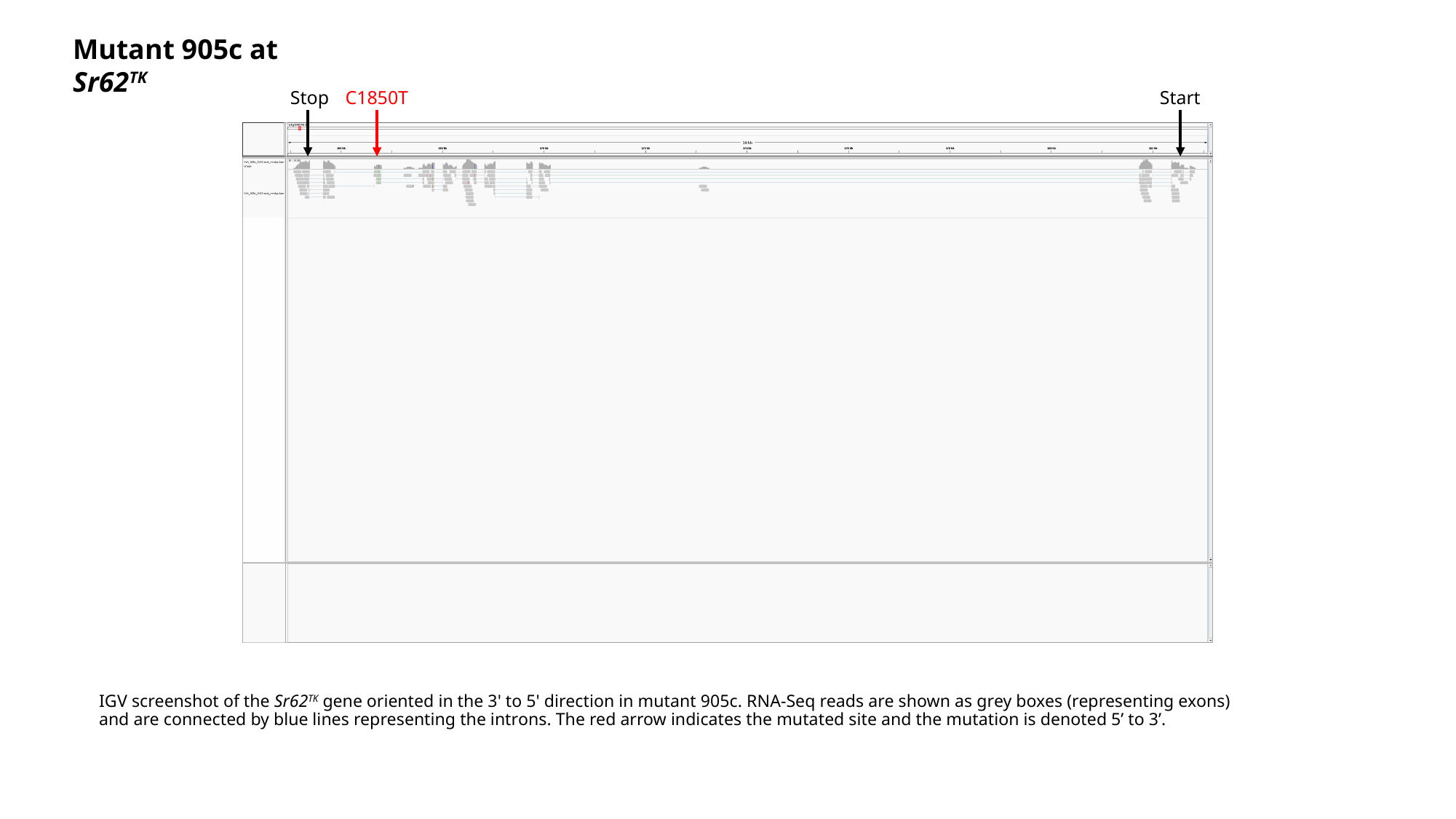

Mutant 905c at Sr62^TK^
Stop
C1850T
Start
IGV screenshot of the Sr62^TK^ gene oriented in the 3' to 5' direction in mutant 905c. RNA-Seq reads are shown as grey boxes (representing exons) and are connected by blue lines representing the introns. The red arrow indicates the mutated site and the mutation is denoted 5’ to 3’.

### Slide 8
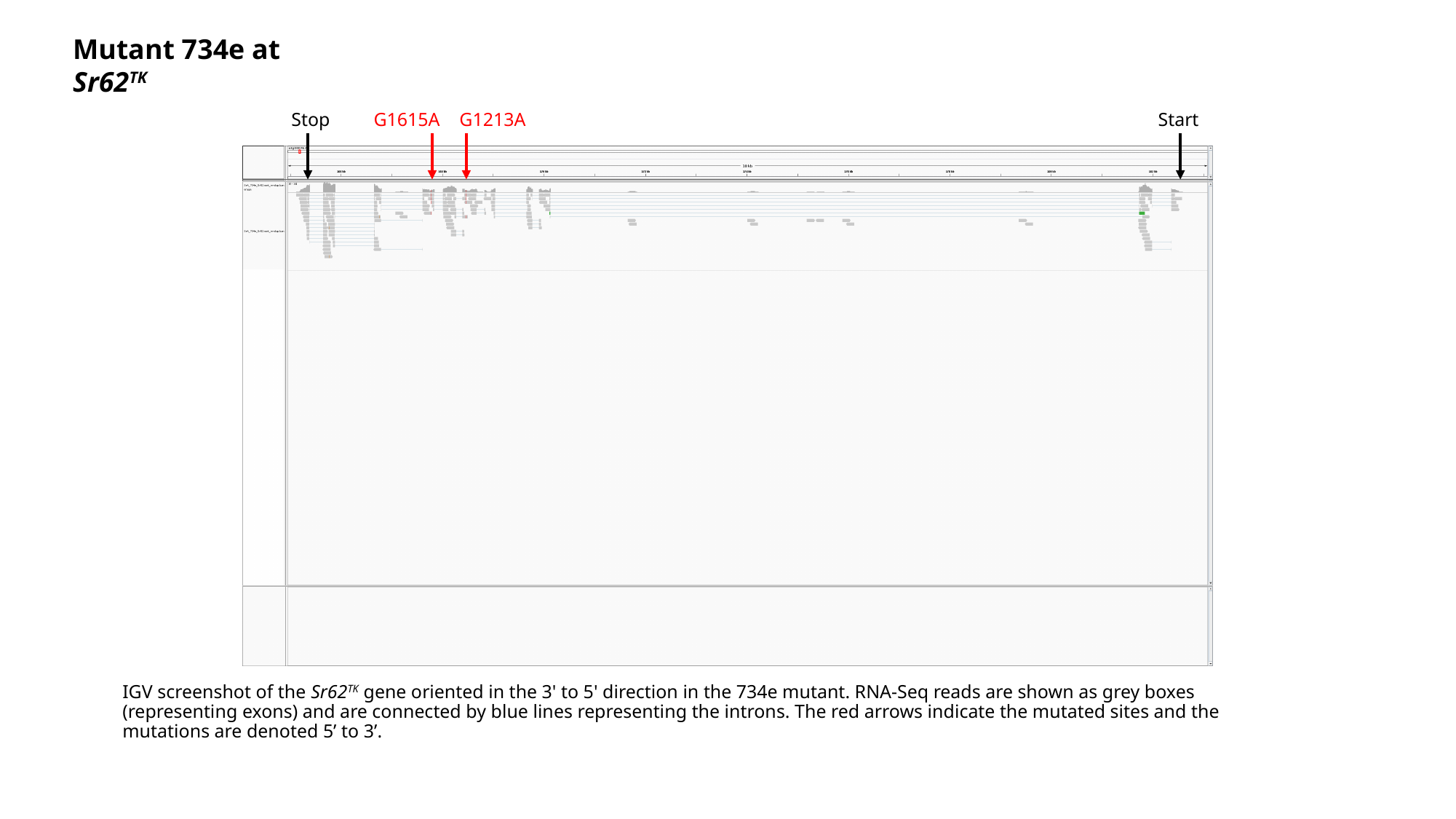

Mutant 734e at Sr62^TK^
Stop
G1615A
G1213A
Start
IGV screenshot of the Sr62^TK^ gene oriented in the 3' to 5' direction in the 734e mutant. RNA-Seq reads are shown as grey boxes (representing exons) and are connected by blue lines representing the introns. The red arrows indicate the mutated sites and the mutations are denoted 5’ to 3’.

### Slide 9
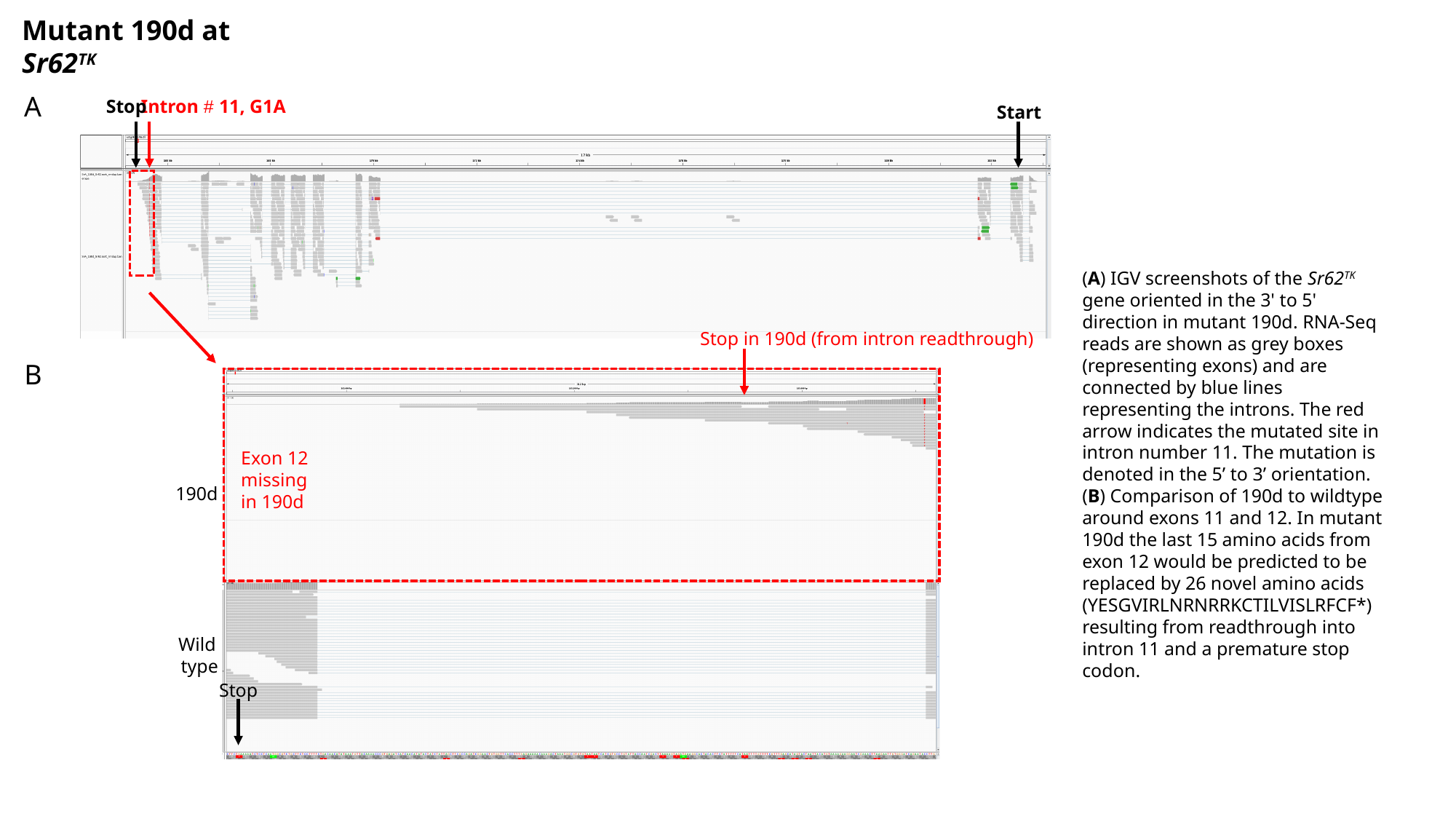

Mutant 190d at Sr62^TK^
A
Stop
Intron # 11, G1A
Start
(A) IGV screenshots of the Sr62^TK^ gene oriented in the 3' to 5' direction in mutant 190d. RNA-Seq reads are shown as grey boxes (representing exons) and are connected by blue lines representing the introns. The red arrow indicates the mutated site in intron number 11. The mutation is denoted in the 5’ to 3’ orientation. (B) Comparison of 190d to wildtype around exons 11 and 12. In mutant 190d the last 15 amino acids from exon 12 would be predicted to be replaced by 26 novel amino acids (YESGVIRLNRNRRKCTILVISLRFCF*) resulting from readthrough into intron 11 and a premature stop codon.
Stop in 190d (from intron readthrough)
B
Exon 12 missing
in 190d
190d
Wild
type
Stop

### Slide 10
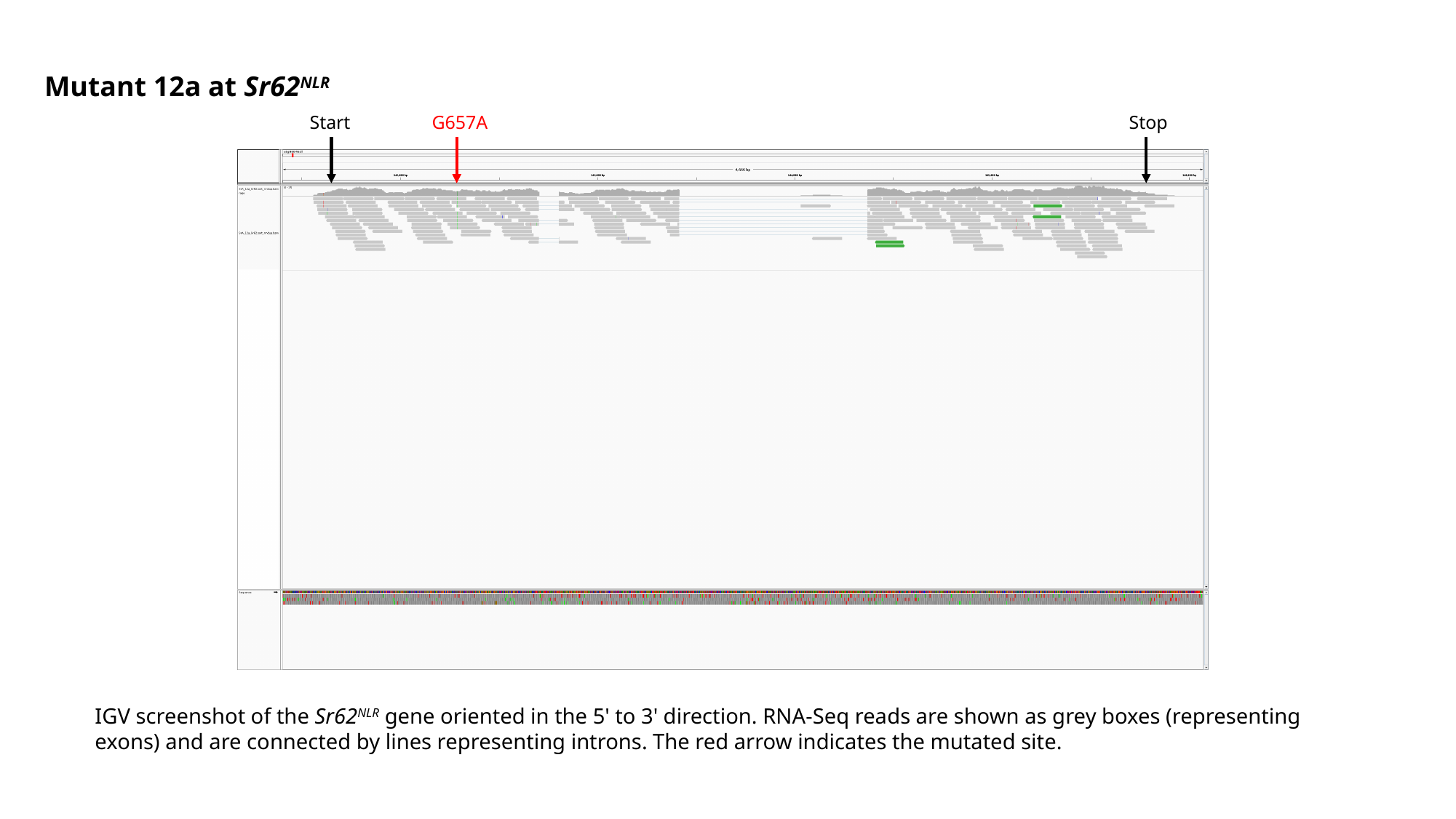

Mutant 12a at Sr62^NLR^
Start
G657A
Stop
IGV screenshot of the Sr62^NLR^ gene oriented in the 5' to 3' direction. RNA-Seq reads are shown as grey boxes (representing exons) and are connected by lines representing introns. The red arrow indicates the mutated site.

### Slide 11
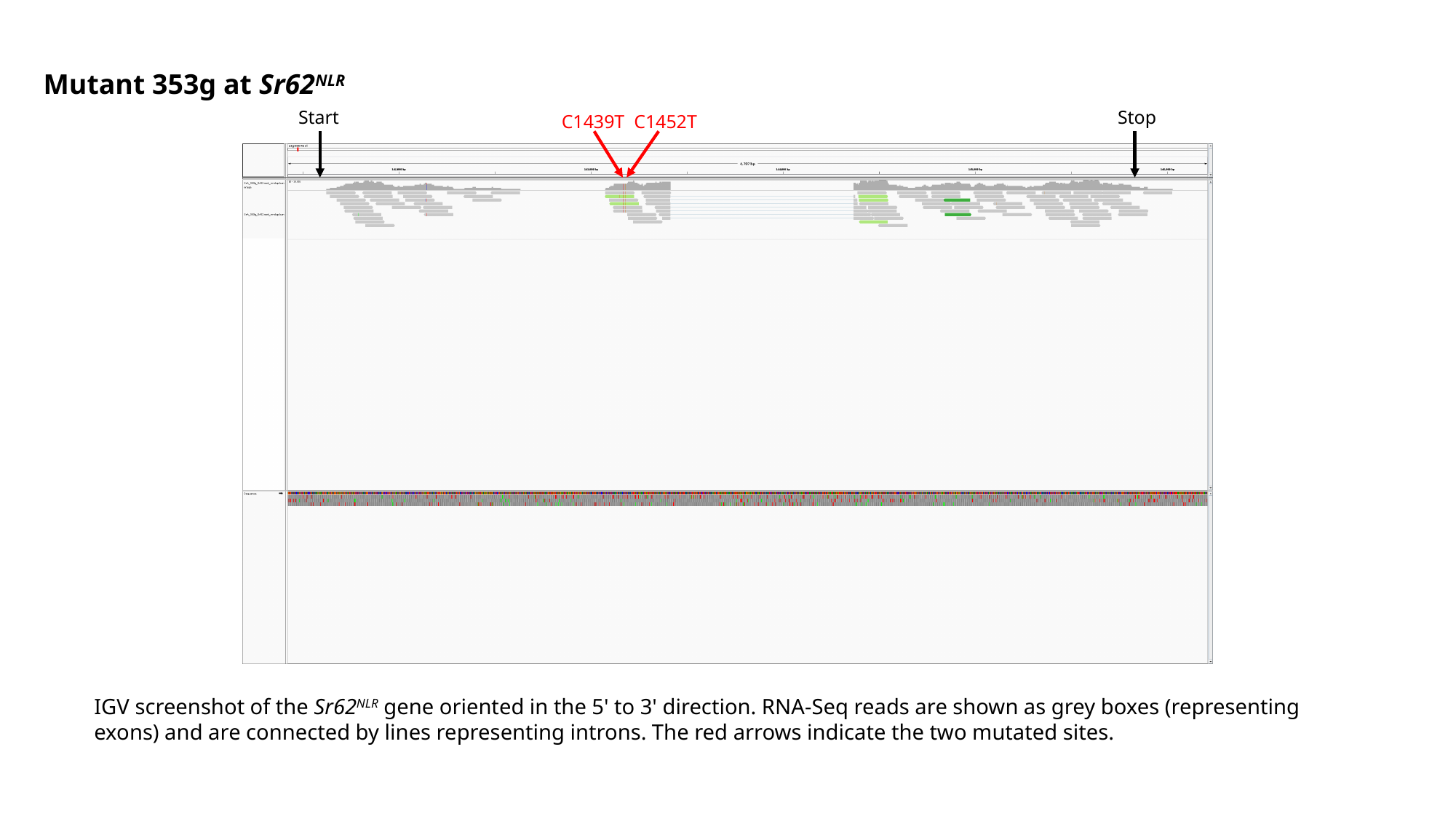

Mutant 353g at Sr62^NLR^
Start
Stop
C1439T C1452T
IGV screenshot of the Sr62^NLR^ gene oriented in the 5' to 3' direction. RNA-Seq reads are shown as grey boxes (representing exons) and are connected by lines representing introns. The red arrows indicate the two mutated sites.

### Slide 12
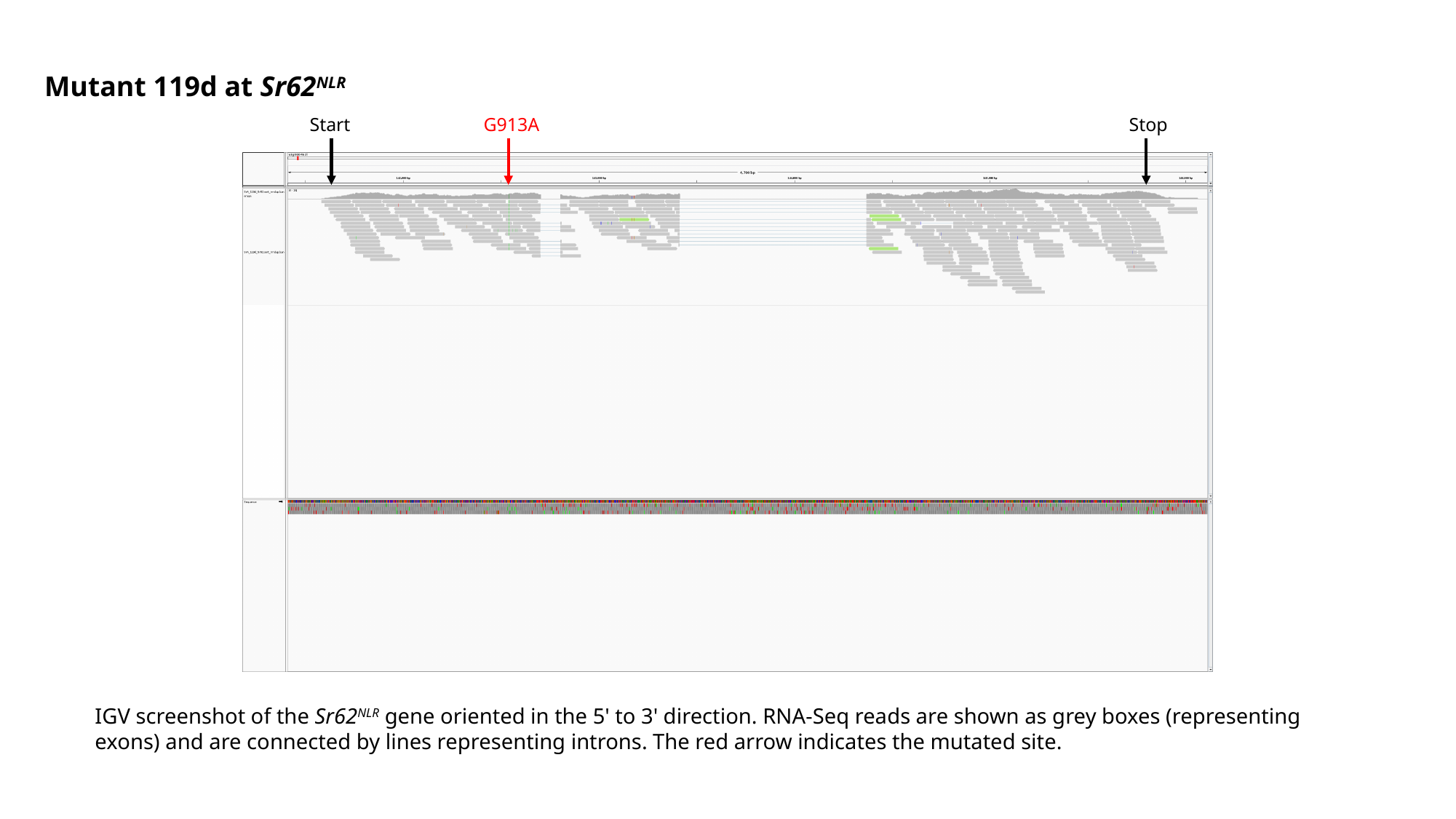

Mutant 119d at Sr62^NLR^
Start
G913A
Stop
IGV screenshot of the Sr62^NLR^ gene oriented in the 5' to 3' direction. RNA-Seq reads are shown as grey boxes (representing exons) and are connected by lines representing introns. The red arrow indicates the mutated site.

### Slide 13
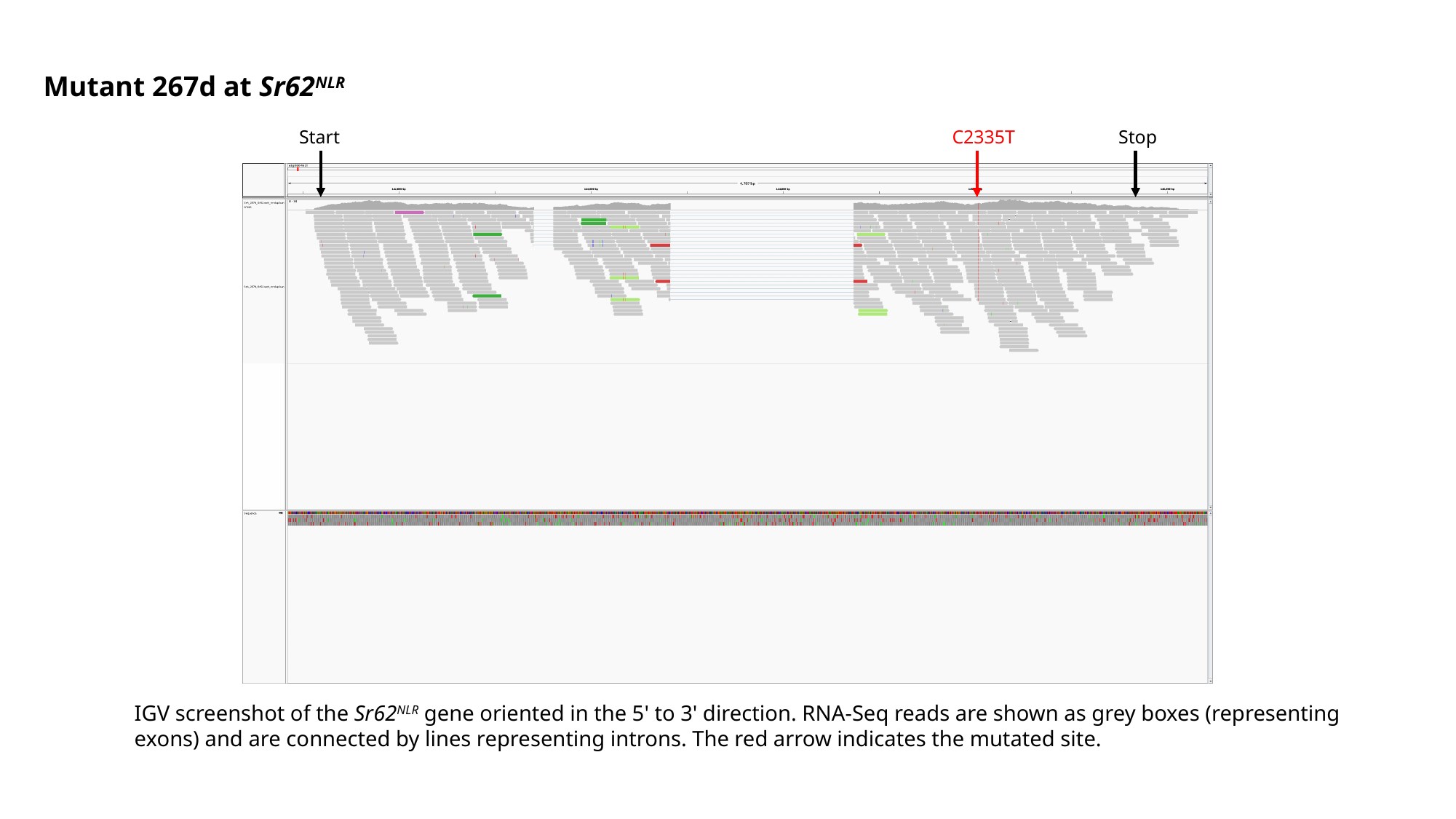

Mutant 267d at Sr62^NLR^
Start
C2335T
Stop
IGV screenshot of the Sr62^NLR^ gene oriented in the 5' to 3' direction. RNA-Seq reads are shown as grey boxes (representing exons) and are connected by lines representing introns. The red arrow indicates the mutated site.

### Slide 14
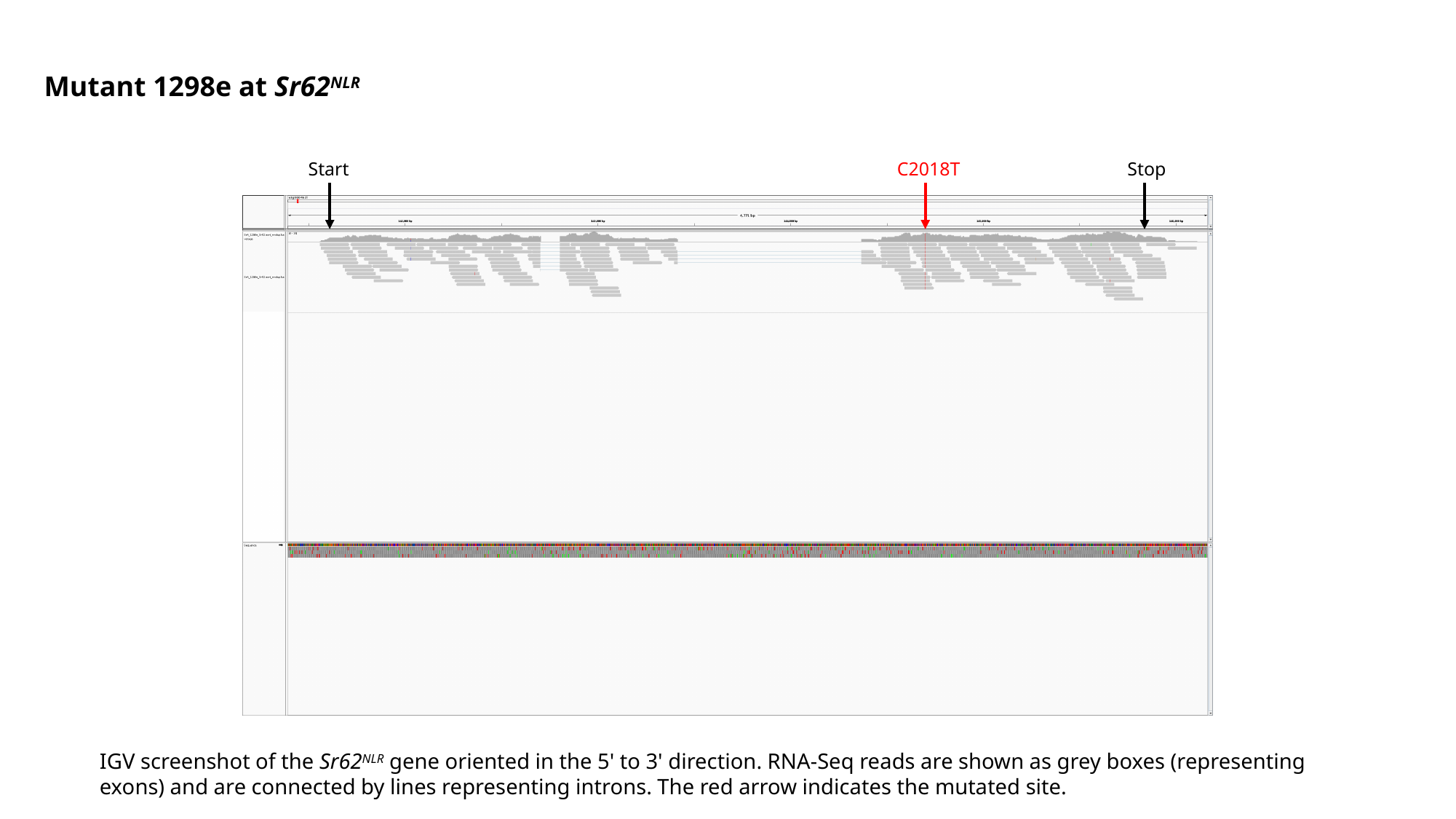

Mutant 1298e at Sr62^NLR^
Start
C2018T
Stop
IGV screenshot of the Sr62^NLR^ gene oriented in the 5' to 3' direction. RNA-Seq reads are shown as grey boxes (representing exons) and are connected by lines representing introns. The red arrow indicates the mutated site.

### Slide 15
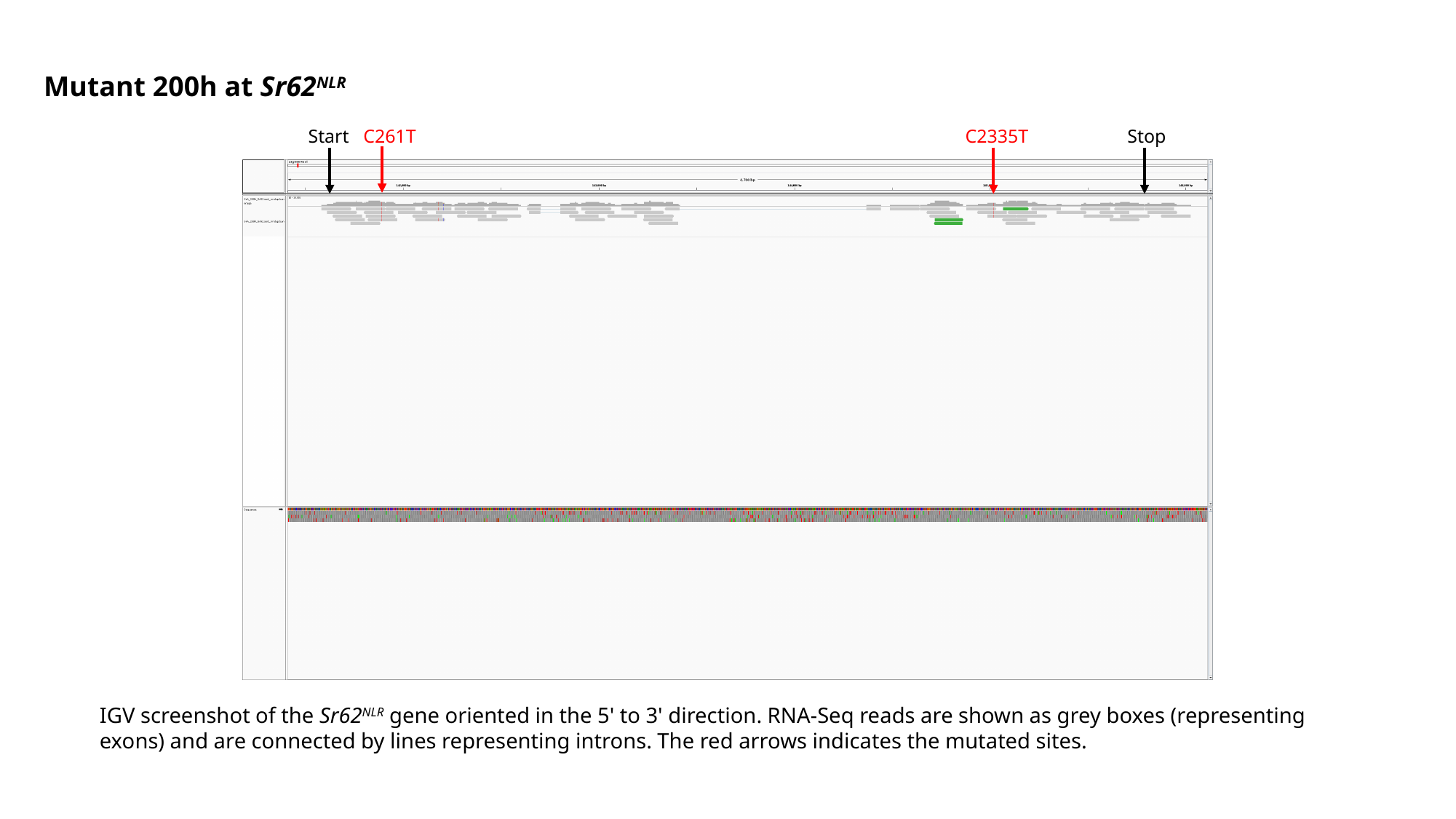

Mutant 200h at Sr62^NLR^
Start
C261T
C2335T
Stop
IGV screenshot of the Sr62^NLR^ gene oriented in the 5' to 3' direction. RNA-Seq reads are shown as grey boxes (representing exons) and are connected by lines representing introns. The red arrows indicates the mutated sites.

### Slide 16
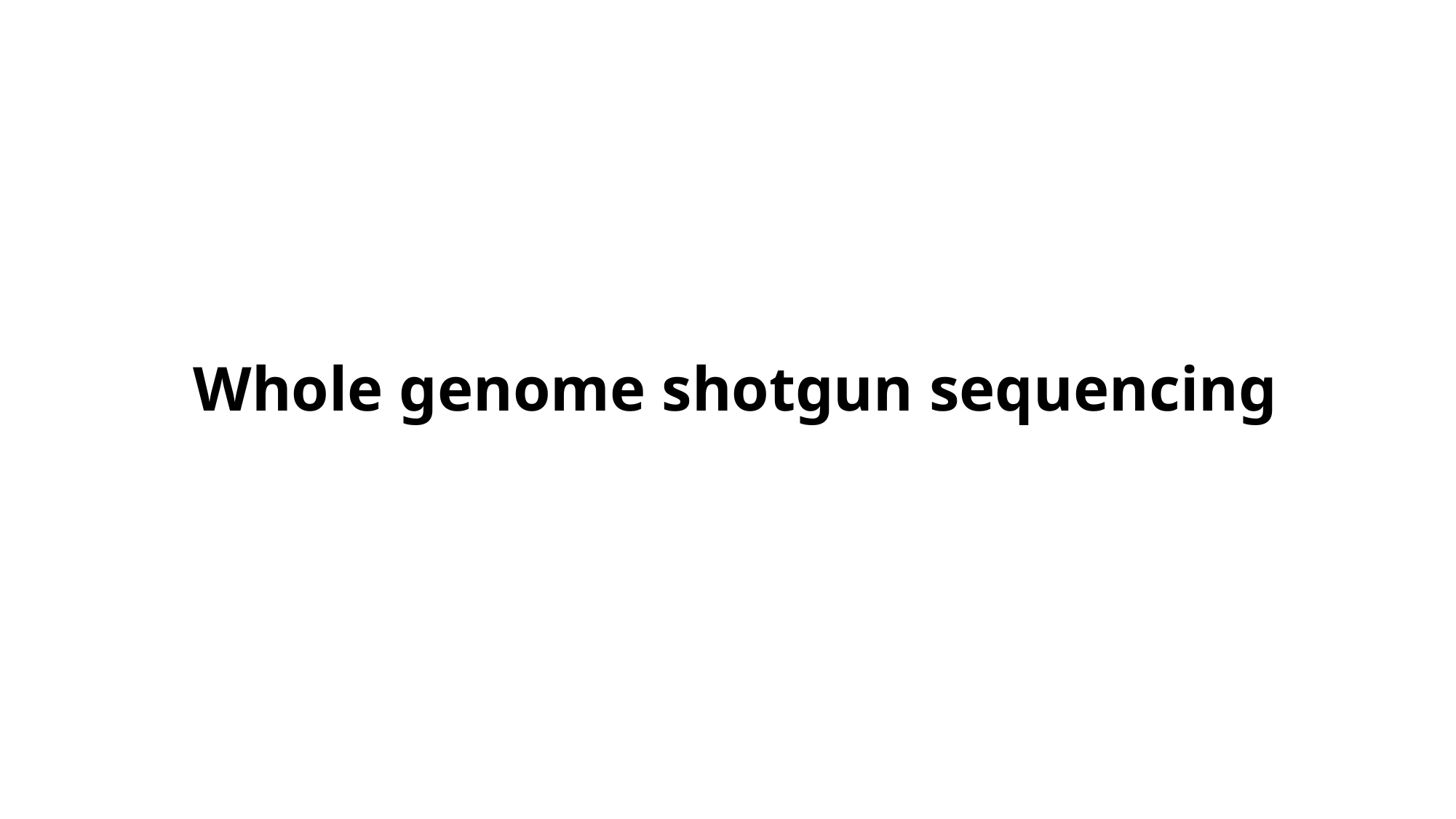

Whole genome shotgun sequencing

### Slide 17
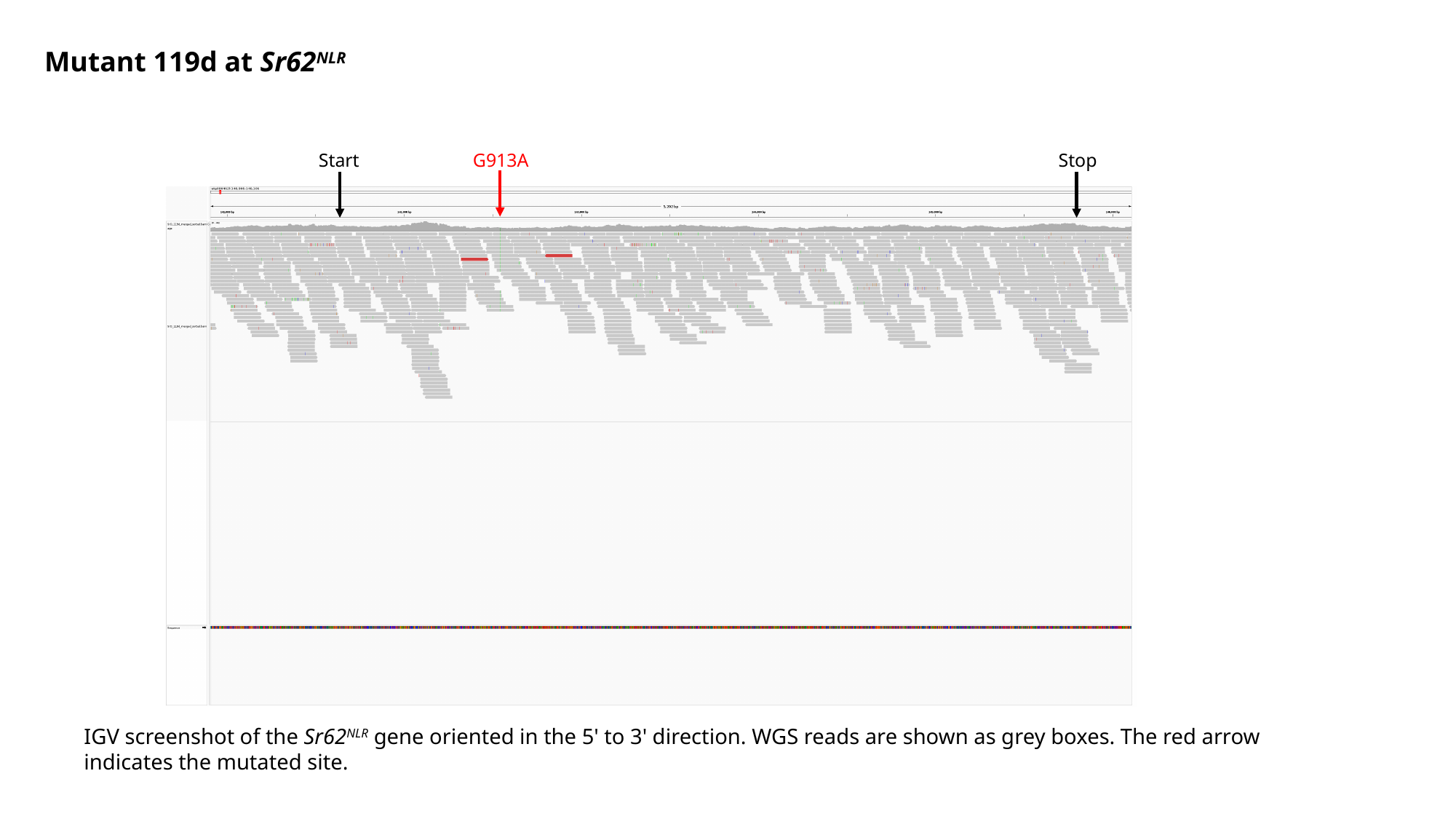

Mutant 119d at Sr62^NLR^
Start
G913A
Stop
IGV screenshot of the Sr62^NLR^ gene oriented in the 5' to 3' direction. WGS reads are shown as grey boxes. The red arrow indicates the mutated site.

### Slide 18
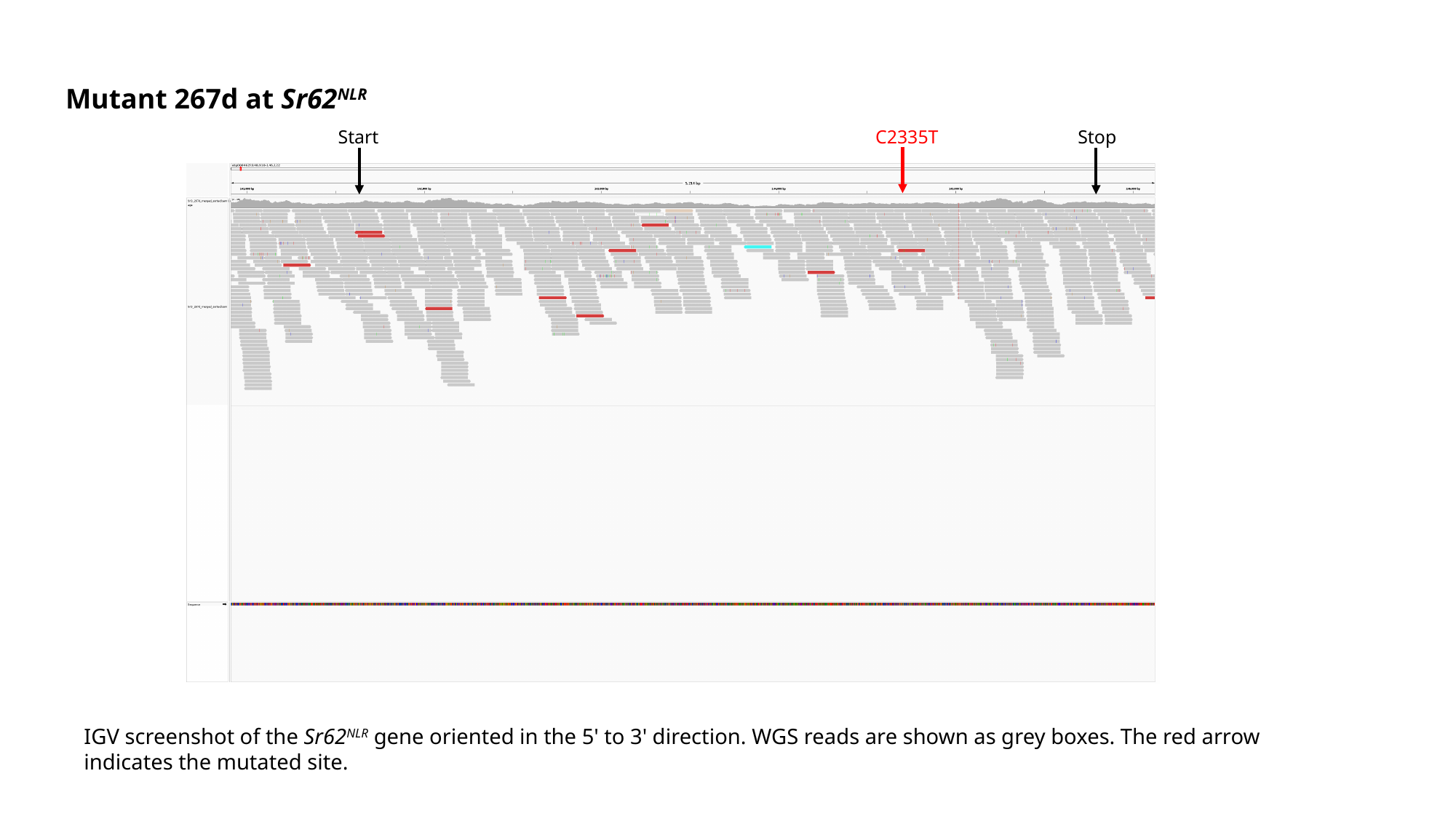

Mutant 267d at Sr62^NLR^
Start
C2335T
Stop
IGV screenshot of the Sr62^NLR^ gene oriented in the 5' to 3' direction. WGS reads are shown as grey boxes. The red arrow indicates the mutated site.

### Slide 19
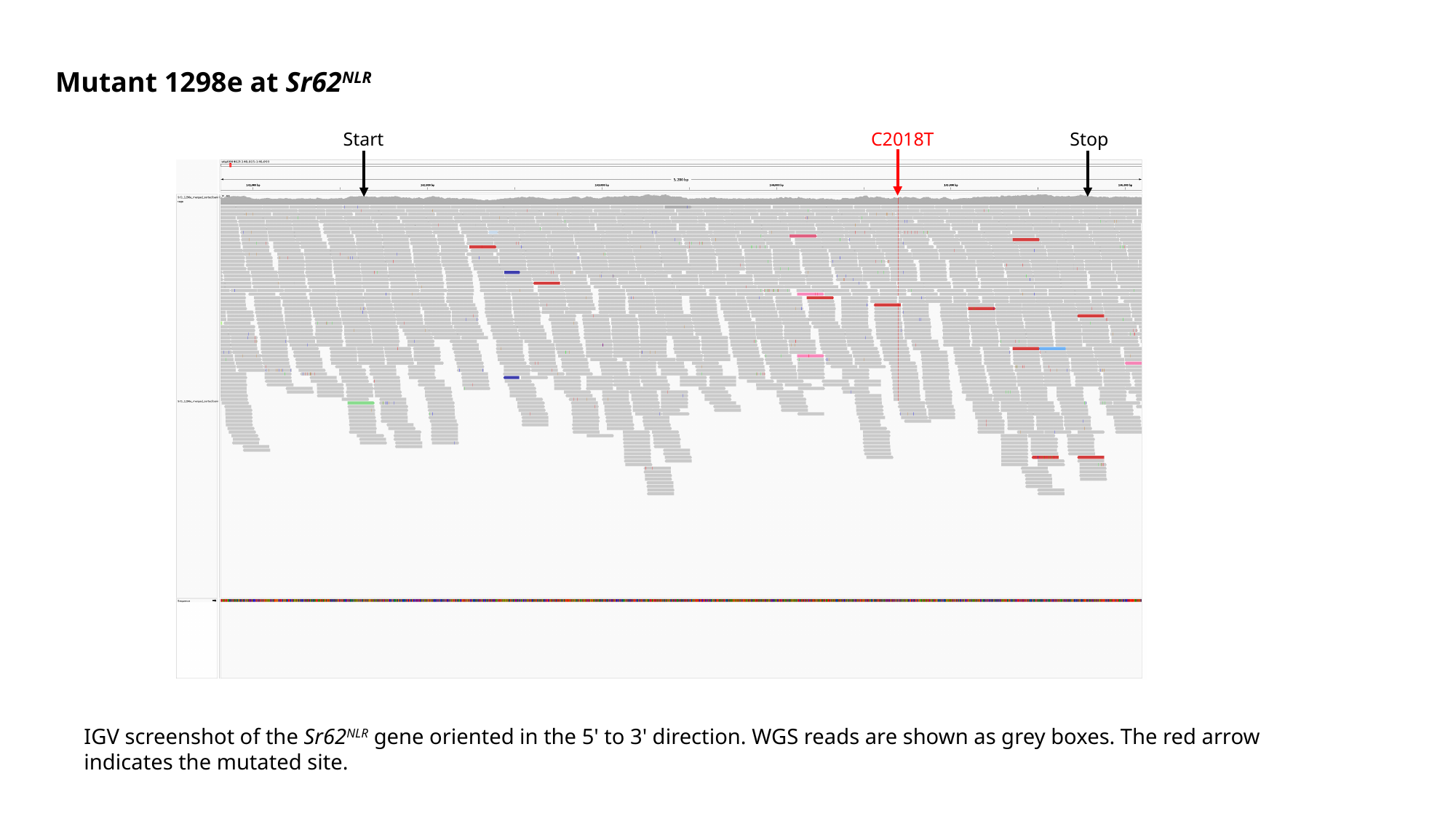

Mutant 1298e at Sr62^NLR^
Start
C2018T
Stop
IGV screenshot of the Sr62^NLR^ gene oriented in the 5' to 3' direction. WGS reads are shown as grey boxes. The red arrow indicates the mutated site.

### Slide 20
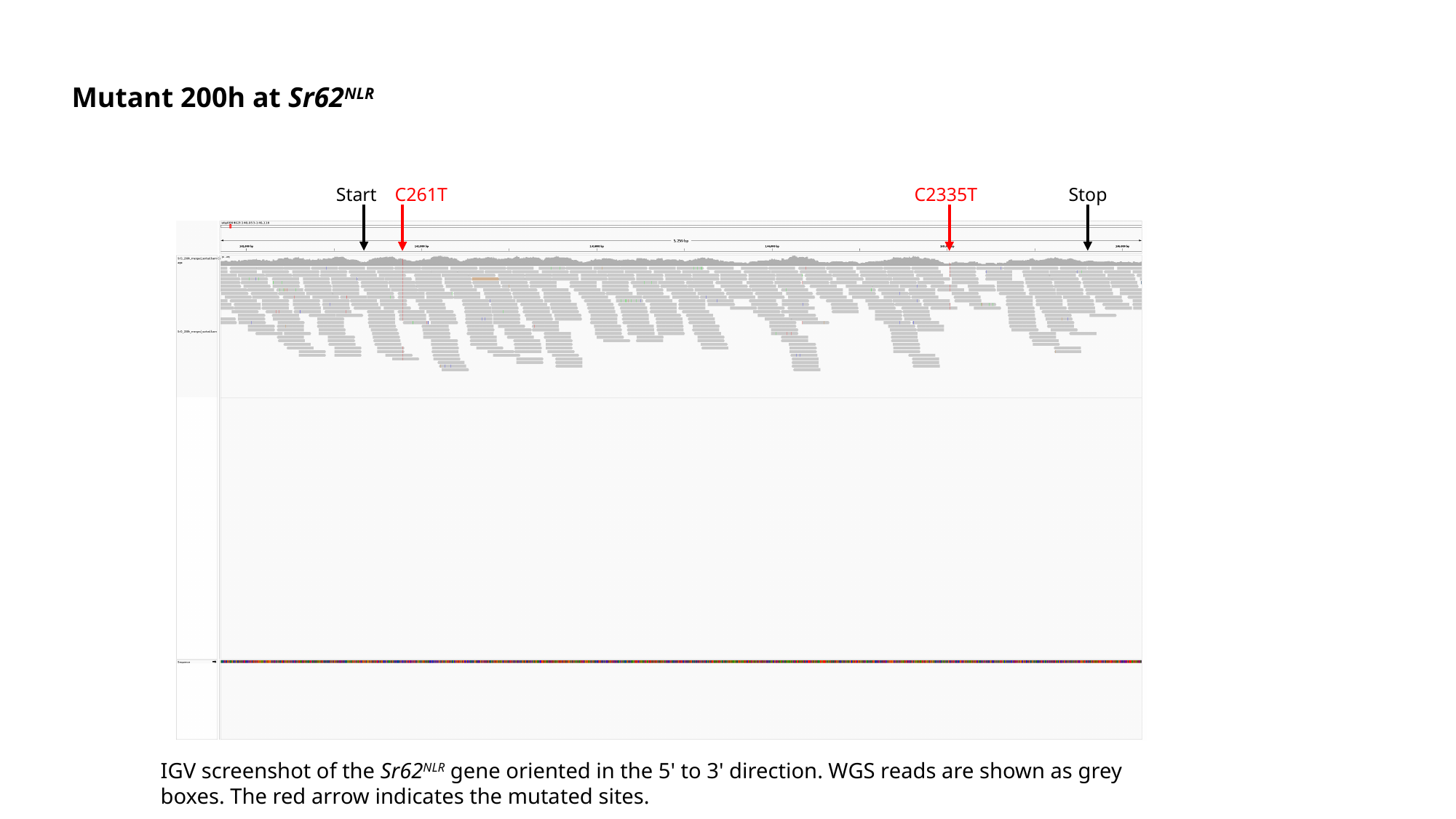

Mutant 200h at Sr62^NLR^
Start
C261T
C2335T
Stop
IGV screenshot of the Sr62^NLR^ gene oriented in the 5' to 3' direction. WGS reads are shown as grey boxes. The red arrow indicates the mutated sites.

### Slide 21
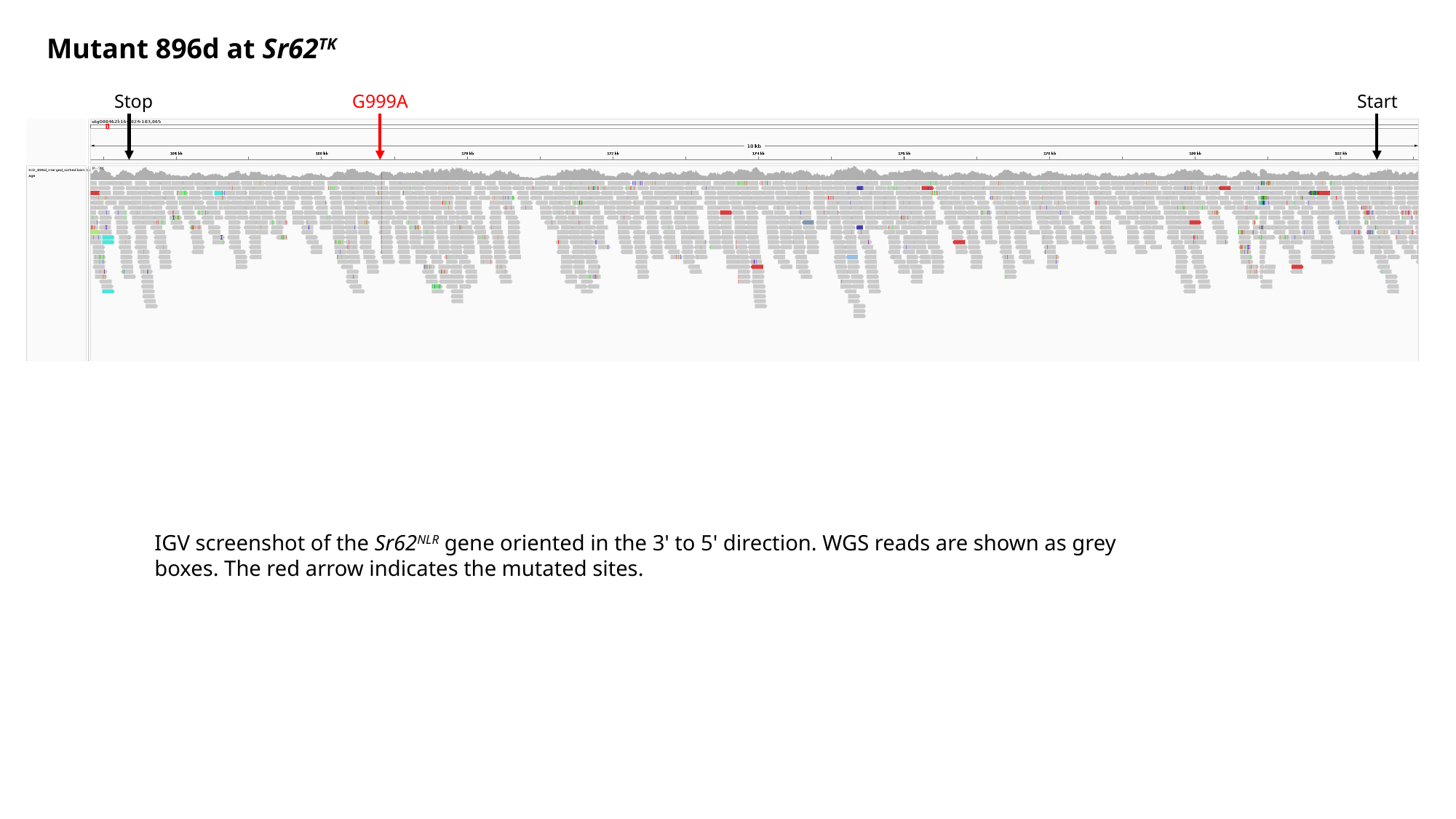

Mutant 896d at Sr62^TK^
Stop
G999A
Start
IGV screenshot of the Sr62^NLR^ gene oriented in the 3' to 5' direction. WGS reads are shown as grey boxes. The red arrow indicates the mutated sites.

### Slide 22
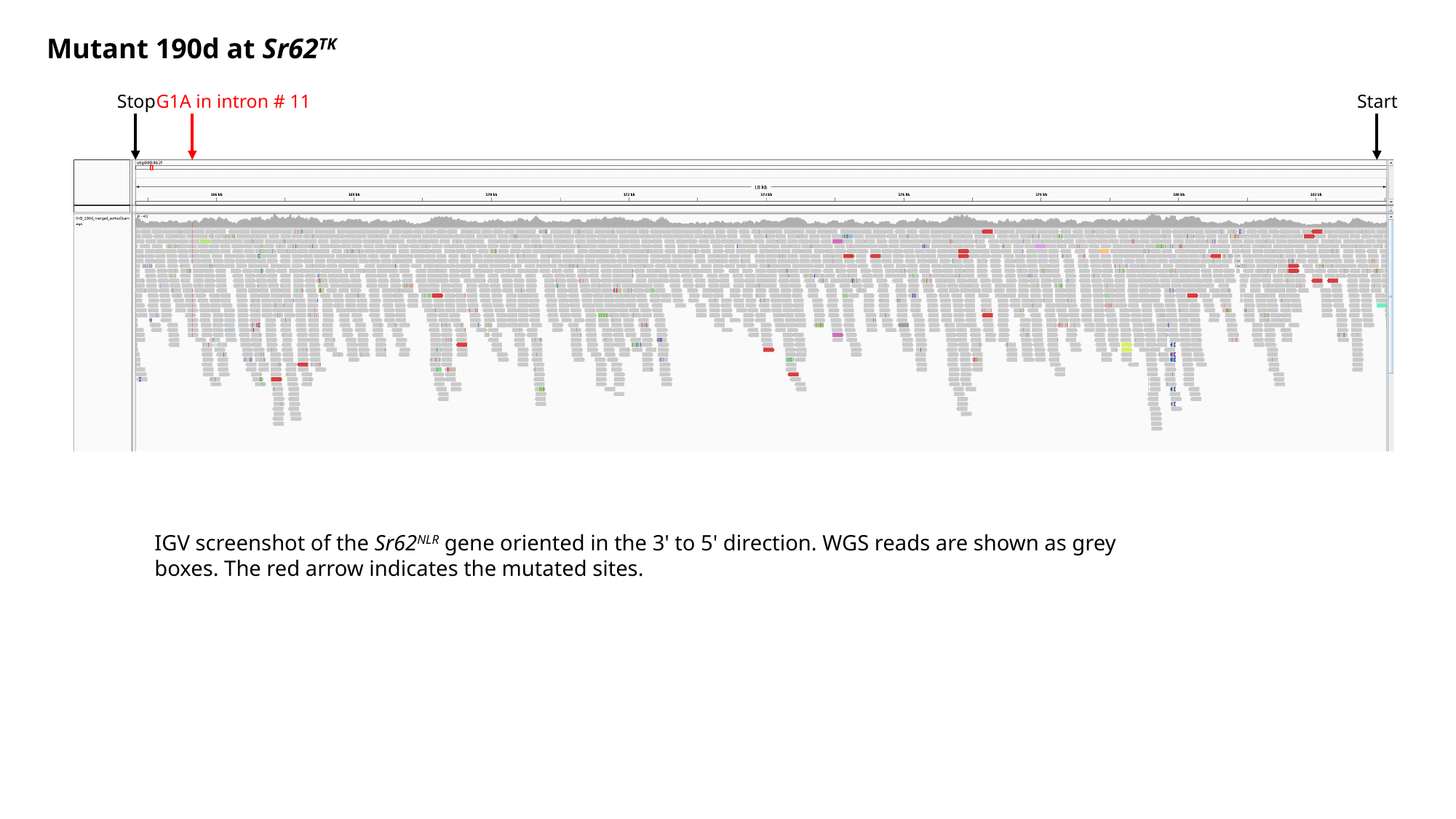

Mutant 190d at Sr62^TK^
Stop
G1A in intron # 11
Start
IGV screenshot of the Sr62^NLR^ gene oriented in the 3' to 5' direction. WGS reads are shown as grey boxes. The red arrow indicates the mutated sites.

### Slide 23
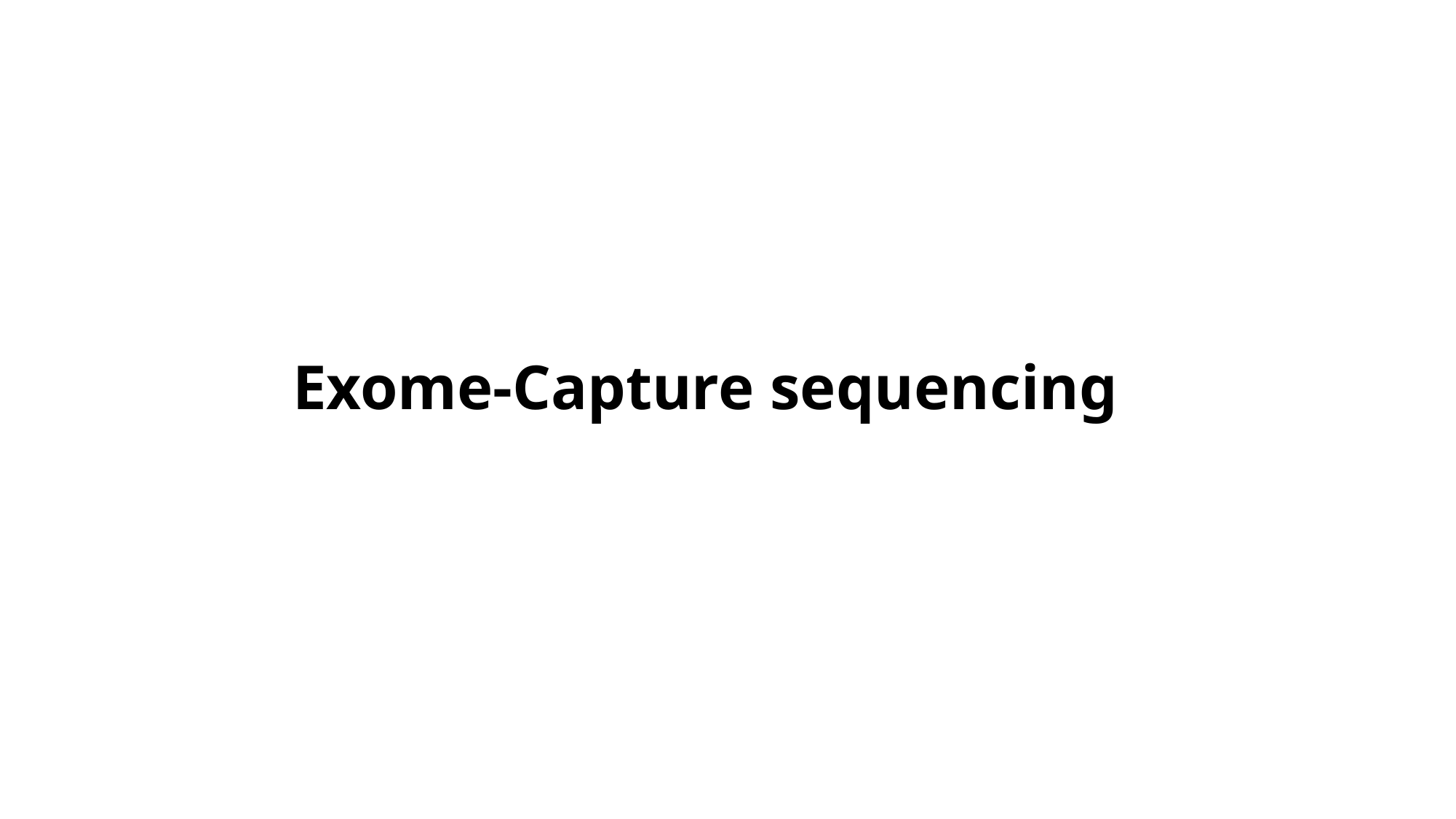

Exome-Capture sequencing

### Slide 24
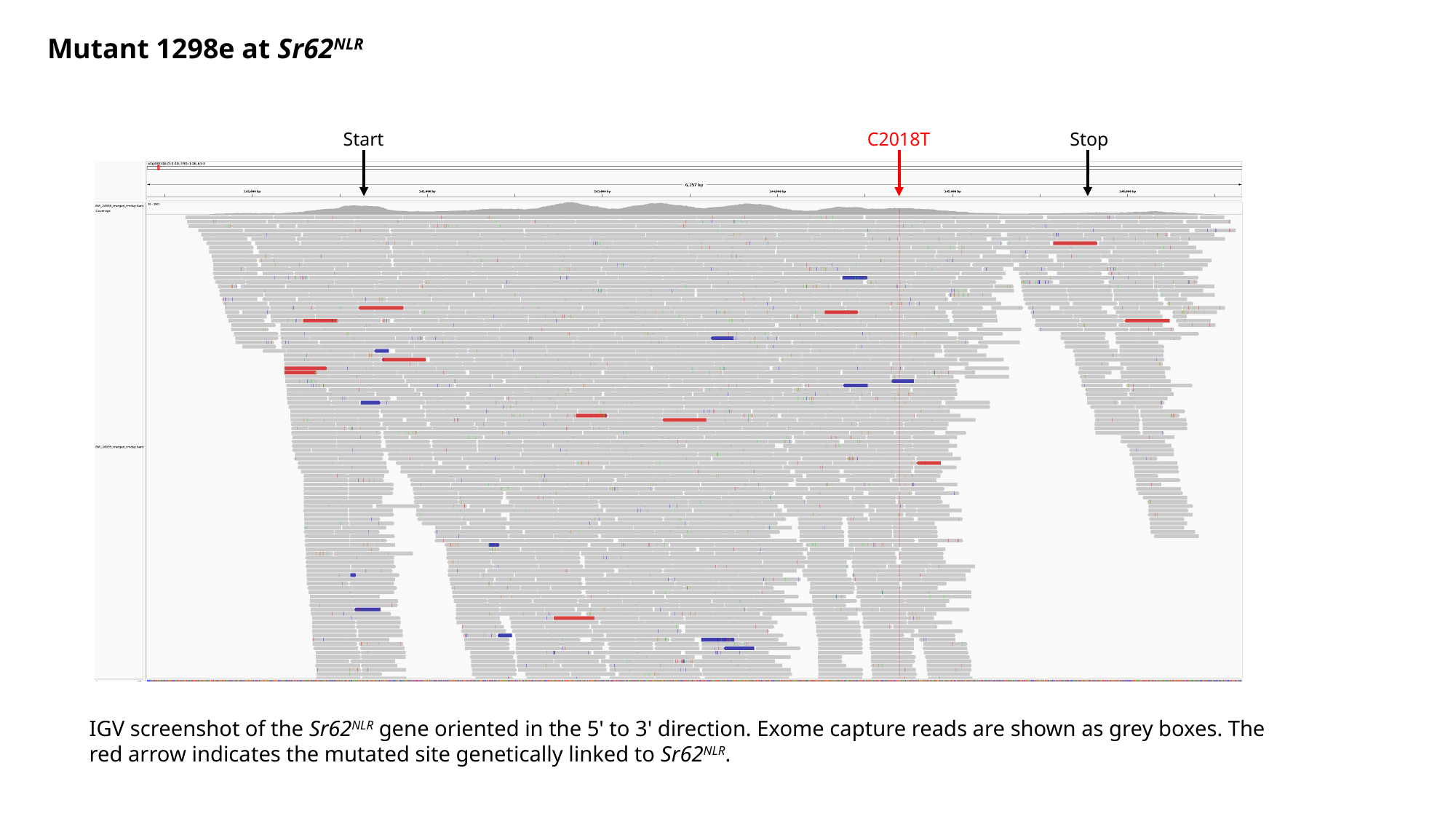

Mutant 1298e at Sr62^NLR^
Start
C2018T
Stop
IGV screenshot of the Sr62^NLR^ gene oriented in the 5' to 3' direction. Exome capture reads are shown as grey boxes. The red arrow indicates the mutated site genetically linked to Sr62^NLR^.

### Slide 25
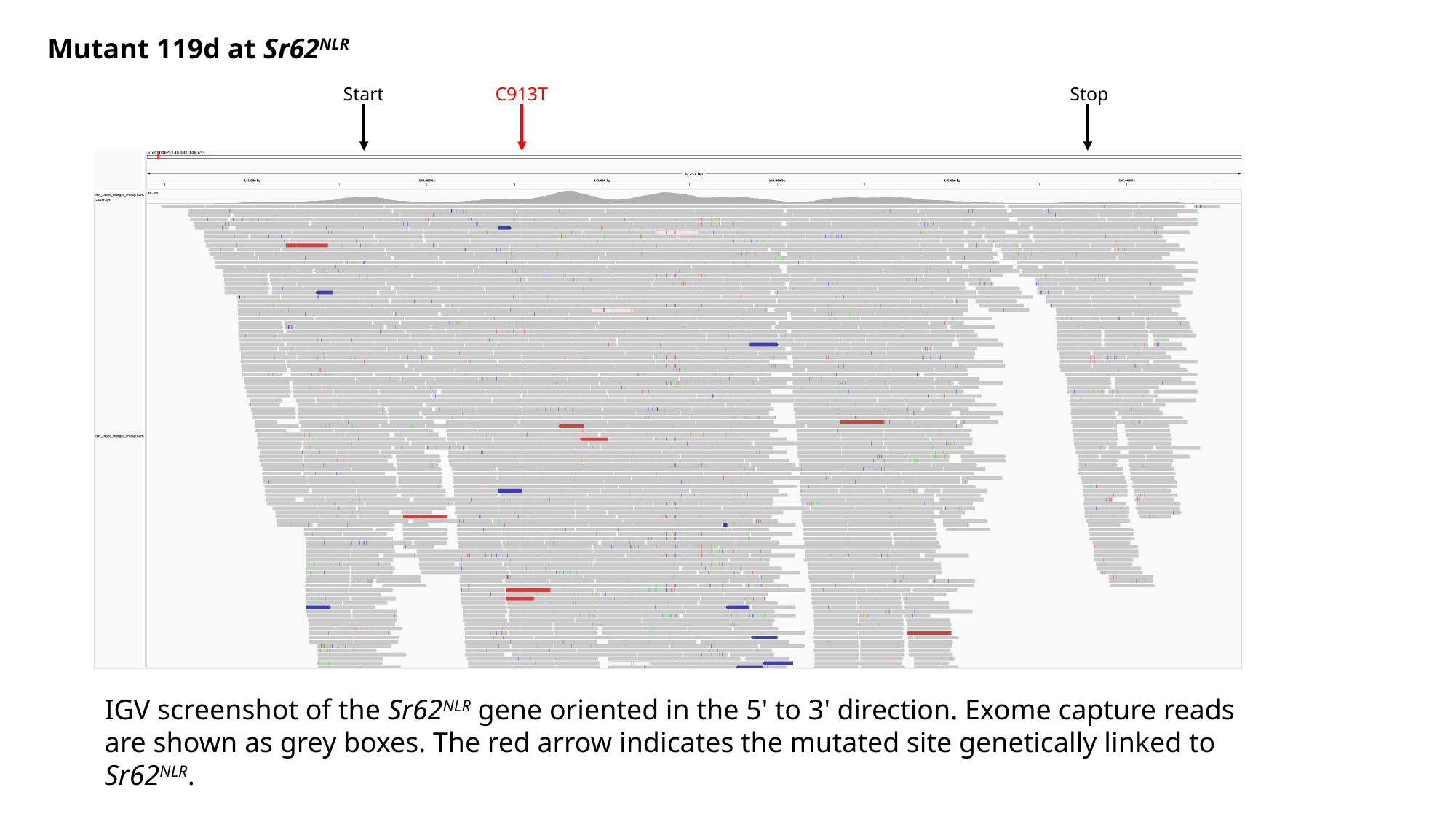

Mutant 119d at Sr62^NLR^
Start
C913T
Stop
IGV screenshot of the Sr62^NLR^ gene oriented in the 5' to 3' direction. Exome capture reads are shown as grey boxes. The red arrow indicates the mutated site genetically linked to Sr62^NLR^.

### Slide 26
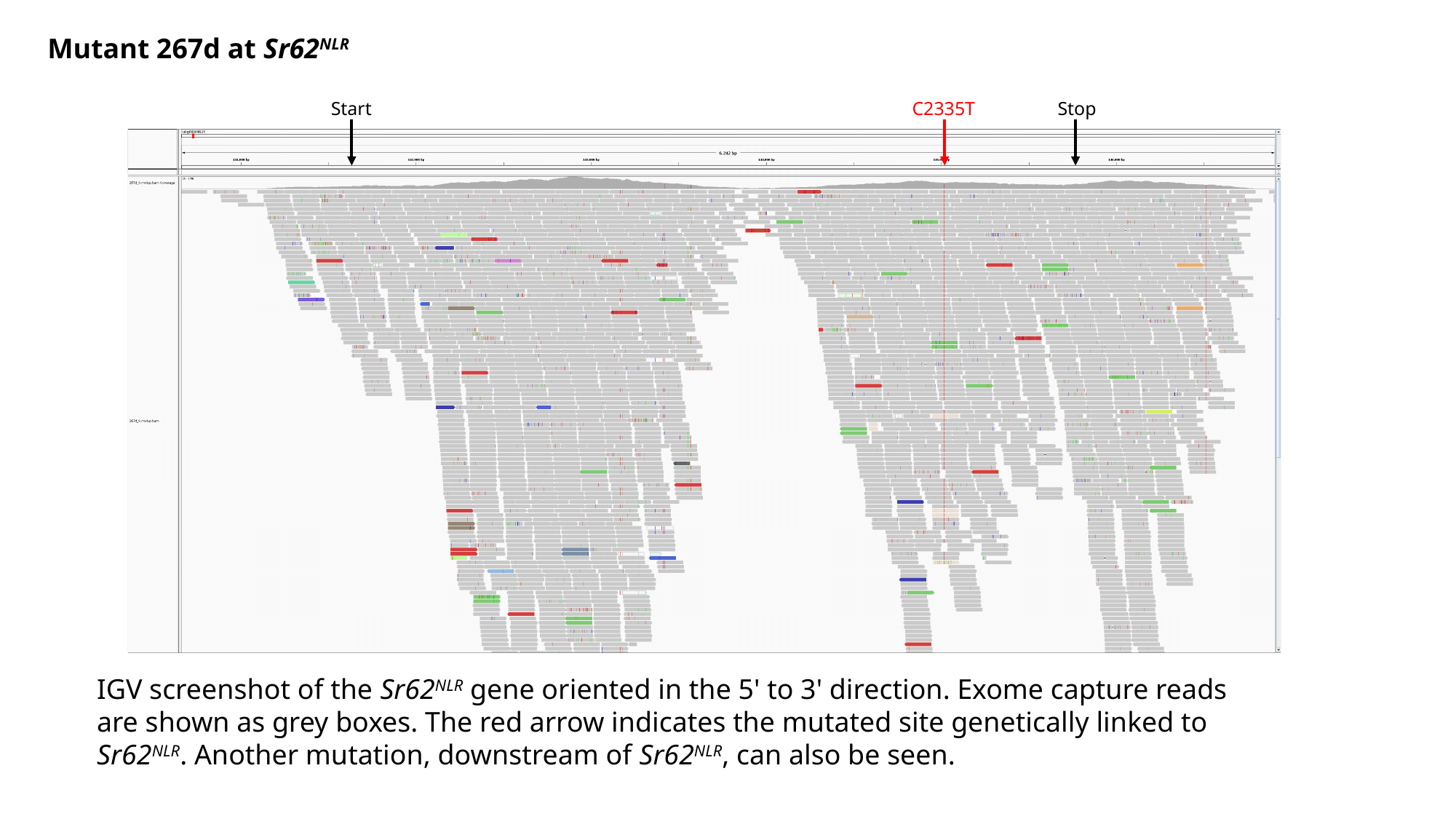

Mutant 267d at Sr62^NLR^
Start
C2335T
Stop
IGV screenshot of the Sr62^NLR^ gene oriented in the 5' to 3' direction. Exome capture reads are shown as grey boxes. The red arrow indicates the mutated site genetically linked to Sr62^NLR^. Another mutation, downstream of Sr62^NLR^, can also be seen.
